# Brief disruption of activity in a subset of dopaminergic neurons during consolidation impairs long-term memory by fragmenting sleep

**DOI:** 10.1101/2023.10.23.563499

**Authors:** Lin Yan, Litao Wu, Timothy D. Wiggin, Xiaojuan Su, Wei Yan, Hailiang Li, Lei Li, Zhonghua Lu, Fang Guo, Zhiqiang Meng, Yuantao Li, Fan Li, Leslie C. Griffith, Chang Liu

**Affiliations:** Department of Anesthesiology, Shenzhen Maternity and Child Healthcare Hospital, Shenzhen, 518028, China; Shenzhen-Hong Kong Institute of Brain Science, Shenzhen Institute of Advanced Technology, Chinese Academy of Sciences, Shenzhen, 518000, China; Faculty of Life and Health Science, Shenzhen University of Advanced Technology, Shenzhen, 518000, China; Department of Biology, Volen National Center for Complex Systems, Brandeis University, Waltham, MA 02454-9110, USA; School of Brain Science and Brain Medicine, Zhejiang University, Hangzhou, 310058, China; Shenzhen Key Laboratory of Drug Addiction, Shenzhen Institute of Advanced Technology, Chinese Academy of Sciences, Shenzhen 518000, China; Department of Pathophysiology, Shantou University Medical College, Shantou, 515041, China

## Abstract

Sleep disturbances are associated with poor long-term memory (LTM) formation, yet the underlying cell types and neural circuits involved have not been fully decoded. Dopamine neurons (DANs) are involved in memory processing at multiple stages. Here, using both male and female flies, *Drosophila melanogaster*, we show that, during the first few hours of memory consolidation, disruption of basal activity of a small subset of protocerebral anterior medial DANs (PAM-DANs), by either brief activation or inhibition of the two dorsal posterior medial (DPM) neurons, impairs 24 h LTM. Interestingly, these brief changes in activity using female flies result in sleep loss and fragmentation, especially at night. Importantly, pharmacological rescue of sleep after manipulation restores LTM. A specific subset of PAM-DANs (PAM-α1) that synapse onto DPM neurons specify the microcircuit that links sleep and memory. MBON-α1 also contributes to the integration of sleep and memory by acting as an additional parallel circuit. PAM-DANs, including PAM-α1, form functional synapses onto DPM mainly via two dopamine receptor subtypes. Dop1R1 primarily mediates the link between sleep and memory. This PAM-α1 to DPM microcircuit exhibits a synchronized, transient, post-training change in activity during the critical memory consolidation window, suggesting an effect of this microcircuit on maintaining the sleep necessary for LTM consolidation. Our results provide a new molecular and circuit basis for the complex relationship between sleep and memory.

## Introduction

Memory consolidation is a time-dependent process which, during a few hours post-encoding, facilitates maturation of newly formed but unstable memories (McGaugh, 2000; Dudai et al., 2015). Neurons undergo a cascade of events that actively remodel the connections at both synaptic and system levels to consolidate newly formed memories (Tonegawa et al., 2018). Systems consolidation, referring to the biological interaction between brain regions or circuits, results in establishment of an information flow for the neuronal ensemble during memory transformation (Dubnau and Chiang, 2013). Sleep, as one of the most important physiological processes that affects memory, has been shown to influence several different memory stages (Wu and Liu, 2023). Sleep deprivation (SD) before training compromises memory formation, while SD post-learning impairs memory consolidation (Li et al., 2009; Spira et al., 2014; Dag et al., 2019). The neural replay of memories during sleep, in particular, has been considered to be part of the active process of memory consolidation (Klinzing et al., 2019). However, the neural mechanisms that bridge sleep and memory consolidation are still largely unclear.

The fruit fly *Drosophila melanogaster* has proven to be a valuable model for uncovering the mechanistic underpinnings of memory formation, and studies in this organism have contributed to revealing the nature of the representations of memory traces formed in associative conditioning paradigms (Aso and Rubin, 2020; Adel, 2021). Classical olfactory conditioning, which is exemplified by studies on Pavlov’s dogs (Pavlov, 1927), forms a memory of a conditioned stimulus (CS, an odor) as a prediction of a pleasant or unpleasant unconditioned stimulus (US, a reward or a punishment) after pairing of the CS with the US (Quinn, 1974; Tempel, 1983; Tully, 1985).

The MB, a multi-functional unit roughly analogous to the hippocampus in mammals, encodes and integrates CS and the US information in a compartmentalized manner. Dopamine (DA) released from DANs acts directly on the MB intrinsic neurons via D1-type receptors (Kim et al., 2007; Qin et al., 2012). Functional observations revealed that ongoing activity in α’β’ neurons is required to maintain memory during the first hour after training (Krashes et al., 2007). In addition, MB αβ core neurons have been shown to act as a gate to permit LTM consolidation (Huang et al., 2012). Furthermore, a recurrent positive feedback loop between dopaminergic PAM-α1 neurons and the MB output neurons MBON-α1 has been shown to be required for the acquisition and consolidation of appetitive LTM (Ichinose, et al., 2015). These data imply that during memory consolidation, a DA signal cascade recruits circuit components at multiple levels to exert sequential modulation of these functional loops. While DANs are an indispensable part of the US pathway, they are also morphologically and functionally diverse (Mao and Davis, 2009; Aso et al., 2014a; Mohammad et al., 2024; Huang C et al., 2024). DANs convey reinforcement signals to spatially segregated subdomains of the mushroom body (MB) to form appetitive or aversive short-term memories (STM) (Aso et al., 2012; Burke et al., 2012; Liu et al., 2012) and also participate in several phases of LTM (Placais et al., 2012). DANs can also participate in post-encoding shaping of memories. Persistent activity in PAM-γ4, PPL1-γ1pedc and -γ2α’1 subtypes interferes with STM and anesthesia-resistant memory (ARM), and promotes forgetting (Berry et al., 2012; Placais et al., 2012; Berry et al., 2015; Lee et al., 2025), while PPL1-α2α’2 and PPL1-α3 mediate aversive LTM formation with persistent post-training activity (Feng et al., 2021). Delayed post-training activity in aSP13-DANs is required for LTM consolidation of courtship learning (Kruttner et al., 2015). The function of many of other subsets remains uncharacterized.

In addition to memory processes, DA also regulates wakefulness and sleep by acting on distinct downstream targets (Kume et al., 2005; Donlea, 2011; Van Swinderen and Andretic, 2011; Ueno et al., 2012; Tomita et al., 2021). Subsets of PAM-DANs have been suggested to signal wakefulness and project to wake-promoting compartments of the MB (Sitaraman et al., 2015; Driscoll et al., 2021).

Additional processing of memory takes place via activation of two pairs of extrinsic neurons, dorsal paired medial (DPM) neurons and anterior paired lateral (APL) neurons, which have been well characterized for their requirement in memory consolidation (Waddell, 2000; Keene et al., 2004; Keene et al., 2006). Furthermore, DPM neurons have been shown to promote sleep via inhibitory signaling (Haynes et al., 2015). The role of the activity of DAN-DPM connections in consolidation as well as the underlying signaling transmission have not been explored.

In this study, we show that PAM neurons form inhibitory connections on DPM neurons. Interestingly, activation of PAM neurons or inactivation of DPM neurons during the first few hours of memory consolidation results in an impairment of LTM. Furthermore, we observed sleep reduction and fragmentation after brief disruption of these neurons. Intriguingly, we identified a specific subset of PAM neurons that are sufficient and necessary to modulate memory consolidation via sleep regulation, and characterized their functional connection with DPM neurons. Maintenance of basal activity of this specific microcircuit during memory consolidation has a profound effect on LTM, through influencing sleep in an internal state-dependent manner. Additionally, we defined the distinct roles of two predominant DA receptors expressed in DPM in mediating sleep and memory processes, respectively. Finally, we characterized additional parallel MBON circuits that contribute to the coupling or segregation of sleep and memory regulation. Together, our findings reveal a microcircuit mechanism by which dopaminergic signaling coordinates sleep and memory consolidation.

## Results

### PAM neurons form functional synapses and inhibit downstream DPM neurons

PAM neurons project to the horizontal lobes of the MB where they participate in formation of appetitive memory (Burke et al., 2012; Liu et al., 2012) (Figure 1A). A pair of DPM neurons intensively innervates all lobes of the MB and is important for consolidation of memory (Yu et al., 2005; Keene et al., 2006) (Figure 1A). The electron microscope (EM) connectome database has shown that PAM and DPM neurons are connected by 4709 synapses (Zheng et al., 2018; Scheffer et al., 2020; Plaza et al., 2022). To determine if PAM and DPM neurons function together to regulate distinct memory processes, we first needed to confirm and characterized the connectome-predicted synapses. To investigate this, we use GFP Reconstitution Across Synaptic Partners (GRASP) and activity-dependent GRASP (Feinberg et al., 2008; Macpherson et al., 2015). Signals from GRASP and active GRASP, which detect physical proximity of neurons, were detected in the horizontal lobes of the MB in flies with split-GFP fragments (spGFP1-10/syb::spGFP1-10 and CD4::spGFP11) expressing in PAM neurons labeled by R58E02-GAL4 or -LexA and DPM neurons labeled by L0111-LexA or eyeless-GAL80;MB-GAL80;c316-GAL4, respectively, showing that PAM neurons likely to connect to DPM neurons functionally (Figure 1B). Further, using a restricted *trans*-Tango, a technique to probe the synaptic connections between GAL4-labeled upstream neurons and LexA-labeled targeted neurons (Sun et al., 2022), we confirmed that DPM neurons labeled by newly generated VT064246-LexA drivers are the downstream target of PAM neurons (Figure 1C-D).

**Figure 1.**
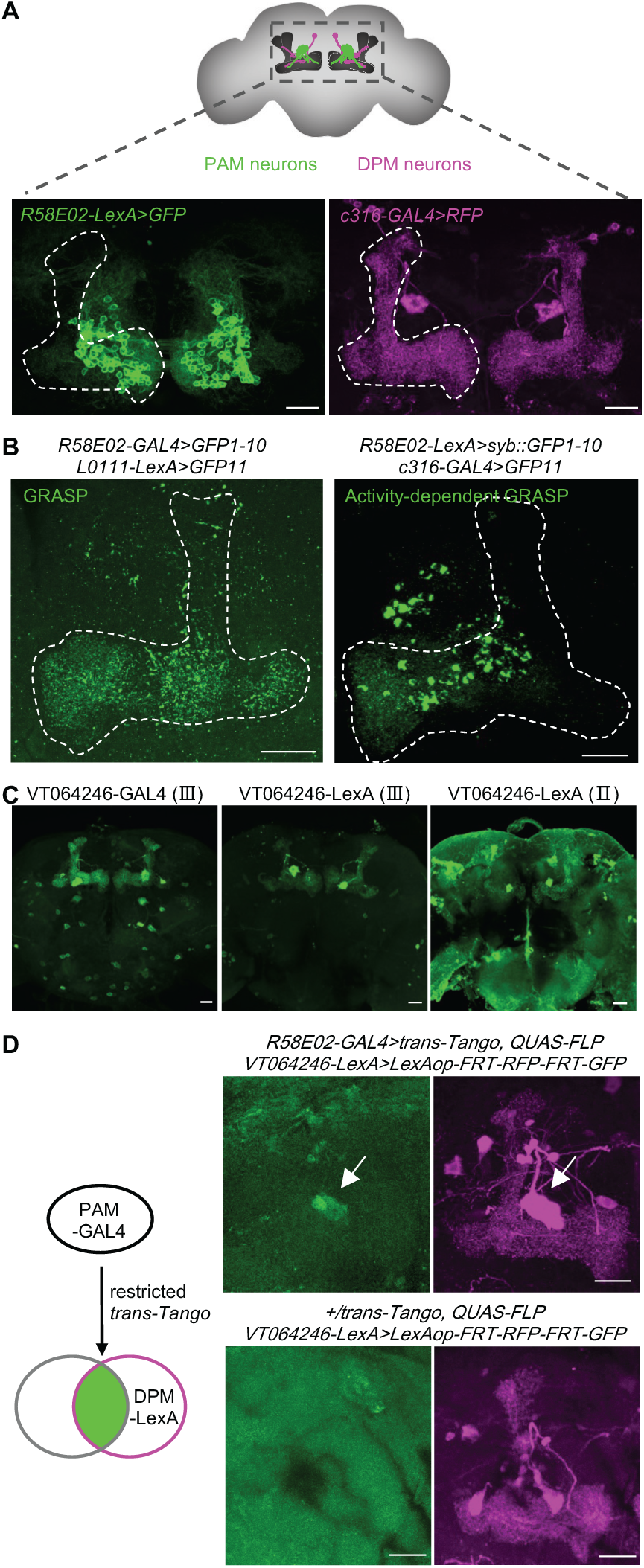
PAM neurons form functional synapses with DPM neurons. (A) The schematic and expression patterns of PAM and DPM neurons. PAM neurons labeled by R58E02-LexA are visualized by GFP (green), and DPM neurons labeled by eyeless-GAL80;MB-GAL80;c316-GAL4 are visualized by RFP (magenta). (B) Fragment of spGFP1-10 and nSyb-GFP1-10 is expressed in R58E02-GAL4 and LexA+ neurons, and fragment of spGFP11 is expressed L0111-LexA and eyeless-GAL80;MB-GAL80;c316-GAL4+ neurons, respectively. Reconstituted GFP signals are shown as puncta (green). The mushroom body is outlined with dashed lines. (C) Newly generated VT064246-LexA (II) and VT064246-LexA (III) recapitulate expression pattern of DPM neurons labeled with VT064246-GAL4. (D) Schematic of restricted *trans*-Tango for direct synaptic contacts between PAM-GAL4 (R58E02) labeled upstream neurons and DPM-LexA (VT064246) labeled targeted postsynaptic to PAM neurons. Anti-GFP-labeled neurons are the downstream of PAM neurons. Anti-RFP-labeled neurons are non-downstream neurons. Right lower panels, the control of restricted *trans*-Tango. Signals of downstream target of DPM neurons was not detected without PAM driver R58E02-GAL4. Scale bar: 20 μm.

To investigate the functional connection from PAM to DPM neurons, we expressed P2X_2_ receptors in PAM neurons using R58E02-LexA, and GCaMP6f in the DPM neurons using VT064246-GAL4. As previously described, we used the percent change in fluorescence over time as a ratio to the initial level, △F/F = (Fn-F_0_)/F_0_× 100% for quantification (Liu et al., 2019). We found that direct activation of PAM neurons by bath application of ATP resulted in a strong decrease in calcium in DPMs, consistent with inhibition of DPM activity (Figure 2A, see also Table 3). In addition, we conducted *in vivo* functional experiments. Optogenetic activation of PAM neurons using the red light-gated cation channel CsChrimson (Klapoetke NC et al., 2014) resulted in a significant reduction in GCaMP signal in DPM neurons (Figure 2B, see also Table 3). Together, these results provide direct evidence that PAM neurons exert inhibitory control over DPM neurons.

**Figure 2.**
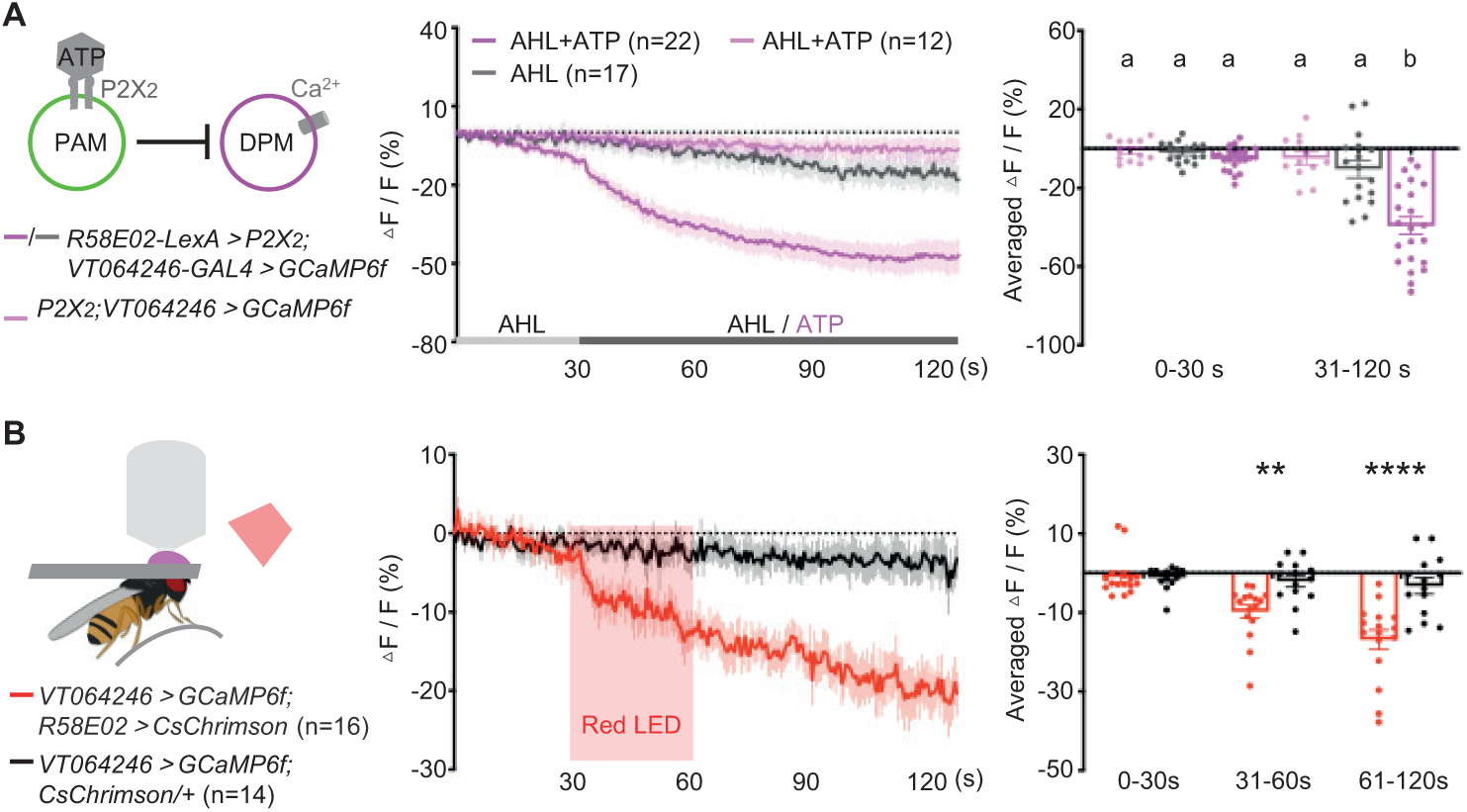
PAM neurons exert direct inhibitory control over DPM neurons. (A) *Ex vivo* functional connection between PAM neurons and DPM neurons. Averaged GCaMP traces and quantification of DPM neurons in response to activation of P2X_2_ expressing PAM neurons by application of 2.5 mM ATP. Experimental genotype under the condition of AHL+ATP, n =22; experimental genotype under the condition of AHL, n = 17; control genotype under the condition of AHL+ATP, n = 12. Groups that are statistically no differences using the same letter, while distinct letters denote statistically significant differences between groups. (B) *In vivo* calcium imaging for functional connection between PAM neurons and DPM neurons. Averaged GCaMP traces and quantification of DPM neurons in response to activation of CsChrimson expressing PAM neurons by red light. The red box indicates the 30-s, 2-Hz, 627-nm light pulse. n = 14-16. **p<0.01; ****p<0.0001. Curves and bar graph are represented as mean ± SEM with individual values. See also Table 3.

### PAM and DPM neurons both influence memory consolidation

To understand the biological function of the inhibitory connection from PAM to DPM neurons, we wanted to ask whether PAM and DPM neurons participate in distinct memory processes or whether they are part of a single pathway. We first confirmed their roles in learning or consolidation, employing an appetitive olfactory conditioning paradigm which is able to induce LTM with a single session of training (Krashes and Waddell, 2008). Blocking the PAM neurons during training using shibire^ts1^ (shi^ts1^), which inhibits neuronal output at high temperature (Krashes et al., 2009), resulted in 24 h appetitive memory defects (Figure 3A, see also Table 1), which is consistent with previous findings (Yamagata et al., 2015).

**Figure 3.**
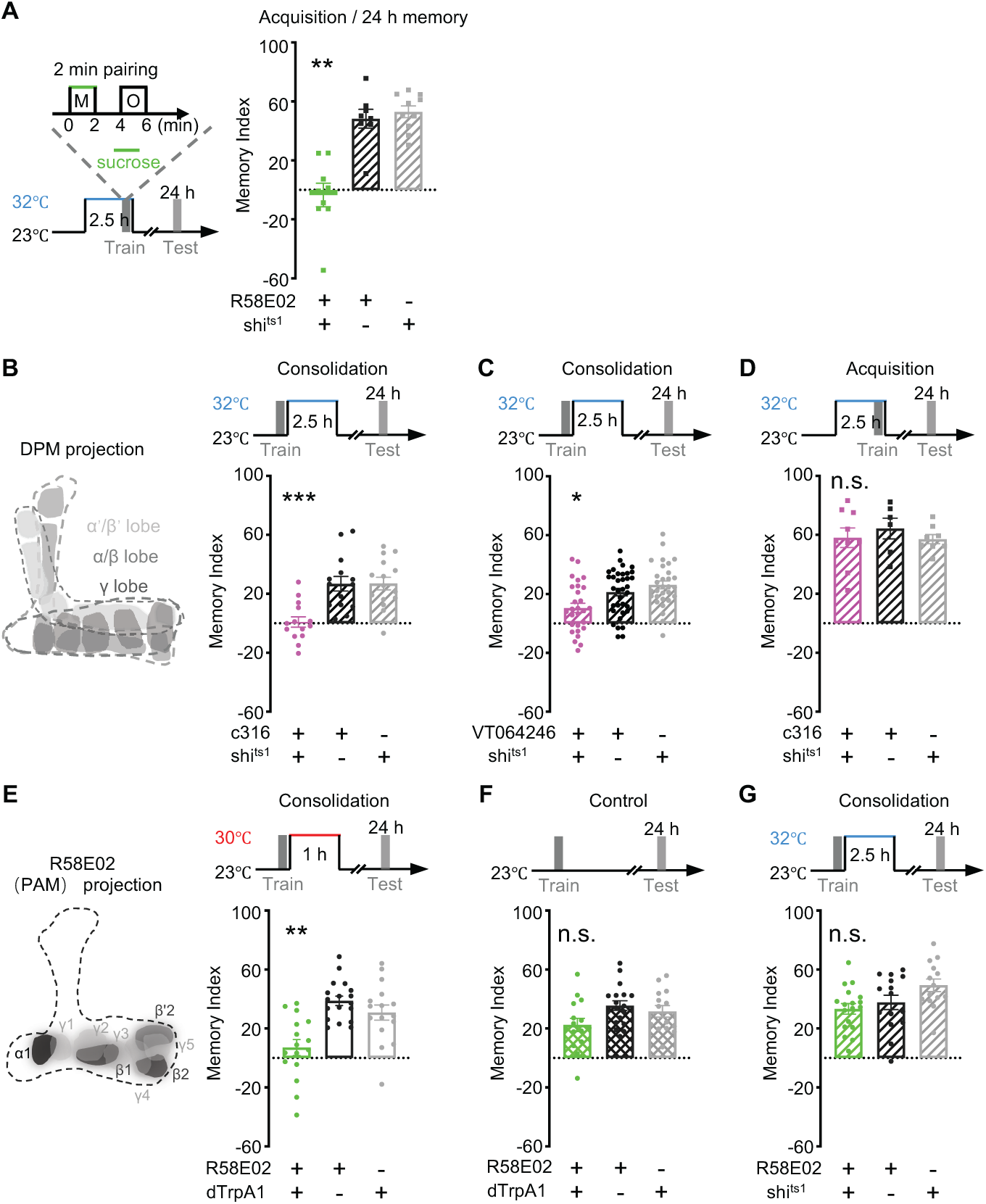
Activation of PAM neurons during consolidation impairs long-term memory, possibly via inhibiting DPM neurons. (A) The paradigm of a single session training with 2 min association of sucrose (Green) and an odor, and the paradigm of 24 h appetitive memory with blocking the output of neurons for 2.5 h before and during training at 32 ℃ (left). Inactivation of PAM neurons during training impaired 24 h memory (Right). n = 8-10. (B) Schematic of innervations of DPM neurons on all lobes of MB (left). Inactivation of DPM neurons for 2.5 h after training disrupts 24 h memory (right). (C) Inactivation of DPM neurons labeled by VT064246-GAL4 after training for 2.5 h impaired 24 h memory. n = 28-30. (D) Inactivation of DPM neurons during training did not impair 24 h memory. n = 6-9. (E) Schematic of innervations of R58E02-GAL4 labeling (PAM) neurons on the horizontal lobes of MB, including subdomains of γ1-5, β’2, β1-2 and α1 (left). Different gray scales were for each compartment’s visibility, not projection intensity. Activation of PAM neurons for 1 h after training disrupts 24 h memory (right). (F) Flies form 24 h appetitive memory without activation of R58E02-GAL4+ neurons. n = 14-18. (G) Inactivation of PAM neurons after training for 2.5 h did not impair 24 h memory. n = 14-15. Red line: 30 ℃; blue line: 32 ℃. n.s.: not significant; *p<0.05; **p<0.01; ***p<0.001; ****p<0.0001. Bar graph are represented as mean ± SEM with individual values. See also Table 1.

DPM neurons have been found to have an essential role in memory consolidation during a critical window for consolidation (Waddell, 2000; Yu et al., 2005; Keene et al., 2006; Cervantes-Sandoval and Davis, 2012). In accordance with the aforementioned findings, inactivation of DPM neurons by shi^ts1^ at restrictive temperature of 32 °C for 2.5 h after training resulted in a 24 h memory deficit (Figure 3B-C, see also Table 1), but there was no effect on 24 h memory when they were blocked before and during training (Figure 3D, see also Table 1).

Since PAM neurons can inhibit DPM neurons, we wondered whether PAM neurons participate in the consolidation process and used conditional activation/inactivation to probe their role. Activation of PAM neurons for 1 h after training resulted in an impairment of 24 h memory (Figure 3E, see also Table 1). Without activation, *R58E02-GAL4/UAS-dTrpA1* flies and their genetic controls formed normal 24 h memory (Figure 3F, see also Table 1). Blocking PAM neuronal activity for 2.5 h after training failed to disrupt 24 h memory (Figure 3G, see also Table 1), suggesting that either continued activity of PAM neurons becomes unnecessary in the consolidation window or that heterogeneity in the subsets of PAM neurons masks any phenotype.

### Increased or decreased activity of PAM neurons results in reduced and fragmented sleep

It has been suggested that one of the important functions of DPM in consolidation is its ability to regulate sleep (Haynes et al., 2015). To ask whether the impairments in LTM that occurred with activation of PAM neurons after training might be attributable to sleep disruption, we monitored sleep under starvation and normal feeding conditions for 30 h, a window sufficient for completion of LTM training and testing, with flies which had been entrained to a 12:12 light:dark cycle. Each panel in Figure 4 shows the schedule for the LTM paradigm we utilize to orient the sleep data to the relevant behavioral phase.

**Figure 4.**
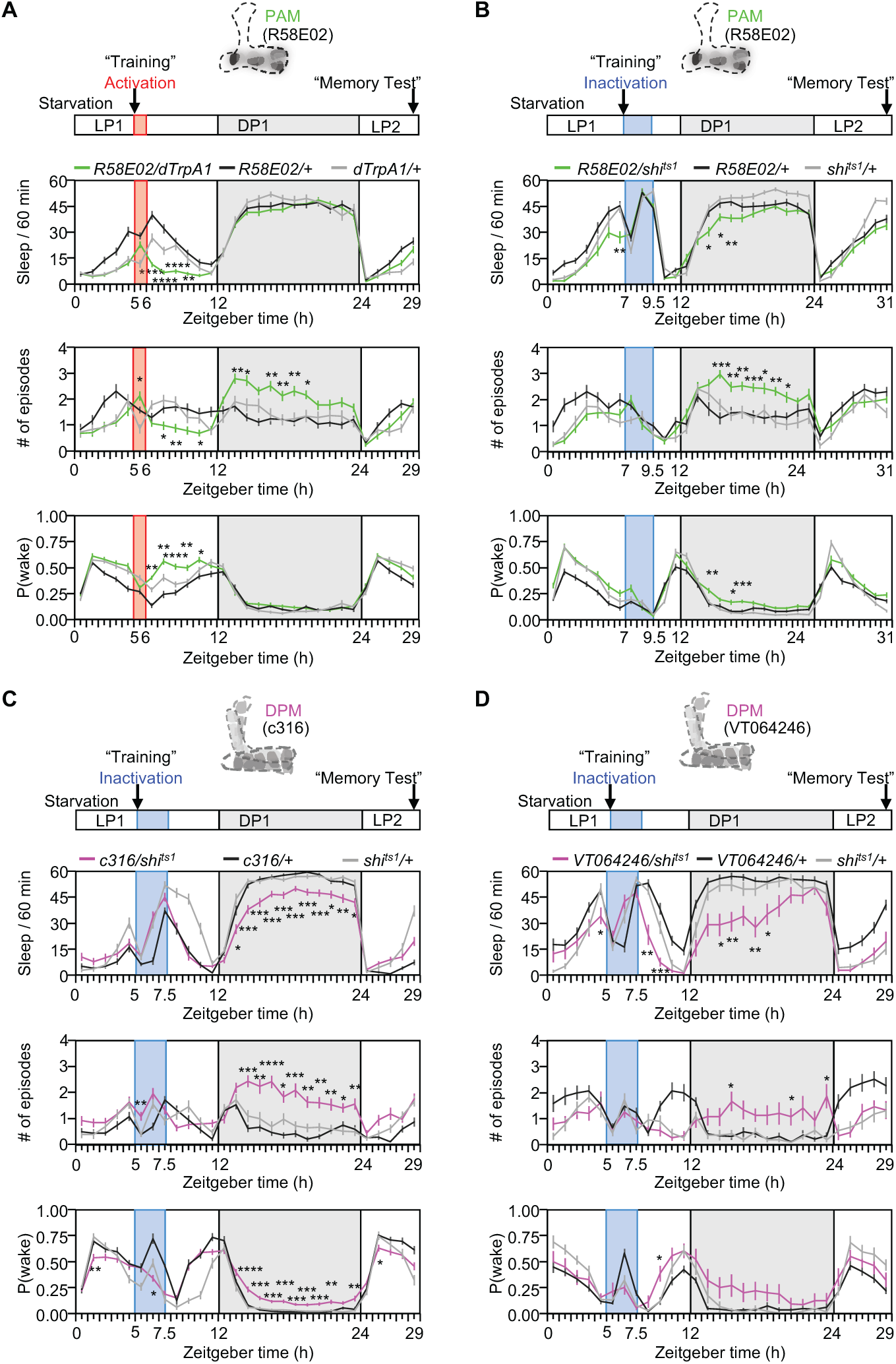
Increased or decreased activity of PAM/DPM neurons results reduced and fragmented sleep. (A) Timeline for sleep recording with activation of PAM neurons for 1 h before, during and after activation till supposed 24 h memory test. Total sleep (upper panel), number of sleep episodes (middle panel), and P(wake) (probability of transition from a sleep to an awake status) (lower panel) in 1 h bin for activation of PAM neurons. n = 40-43. (B) Inactivation of PAM neurons labeled by R58E02-GAL4 under starvation resulted in a significant decrease of sleep at early night, with increased number of sleep episodes and P(wake). n = 33-39. (C) Timeline for sleep recording with inactivation of DPM neurons labeled by c316-GAL4 for 2.5 h before, during and after activation till supposed 24 h memory test. Total sleep, number of sleep episodes, and P(wake) in 1 h bin for inactivation of DPM neurons. n = 33-43. (D) Inactivation of DPM neurons labeled by VT064246-GAL4 with starvation resulted in a decreased and fragmented sleep after activation at night, but almost no effect on P(wake). n = 11-37. Red line: 30 ℃; blue line: 32 ℃; LP1: light period on day 1; red box: activation of neurons for 1 h; DP1 (grey box): dark period on day 1; LP2: light period on day 2; ZT: Zeitgeber time. See also Table 4.

Using the Drosophila Activity Monitor (DAM2) System. We measured sleep amount, the number of sleep episodes and P(wake), a parameter which reflects sleep depth (Liu et al., 2019; Wiggin et al., 2020). Sleep data were analyzed in one-hour bins to assess dynamic changes during the course of starvation before the time when training would have occurred through the time when the 24 h memory tests would have been done (Figure 4, see also Table 4). Activation/inactivation periods were applied in the same time windows that reliably disrupted LTM 1 h for the PAM neurons and 2.5 h for the DPM neurons, i.e. immediately after when training would have occurred.

Activation of PAM neurons at the time which training occurred caused a reduction in sleep occurred a few hours after the activation of PAM neurons (Figure 4A, upper panel), possibly due to a reduced number of sleep episodes (Figure 4A, middle panel) and an increased possibility of waking (Figure 4A, lower panel). The biggest change, however, was structural-PAM activation significantly increased the number of sleep episodes, which reflects increased wakefulness during the dark period following activation (Figure 4A, middle panel). Although LTM impairment was similar in the memory experiments, the effect of inactivation of DPM neurons resulted in a slightly stronger, long-lasting sleep reduction and fragmentation that was delayed until the dark period after inactivation (Figure 4C-D). However, activation of DPM neurons failed to distrupt sleep and impair LTM (Supplemental Figure 1A-B). Unexpectedly, inactivation of PAM neurons led to a sleep change similar to that seen with inactivation of DPM neurons (Figure 4B), suggesting that perhaps a specific subsets of PAM neurons was important for linking memory consolidation and sleep via modulation of DPM. The weak effect of activation of the large R58E02-GAL4+ group of PAM neurons on sleep could be due to masking of the role of a PAM subgroup by inter-PAM interactions or DPM-independent effects on sleep.

Appetitive LTM expression requires that the animal be motivated by hunger (Liu et al., 2012) and starvation is known to suppresses sleep (MacFadyen, 1973; Keene et al., 2010; Thimgan et al. 2010). To determine whether the memory defect and sleep disruption are linked to a certain internal state, we also analyzed the sleep effects of manipulation of PAM and DPM neurons under a no starvation condition (Figure 5). Without starvation, activation of PAM neurons led to a significant decrease in total sleep during the activation period, and also to a significant reduction in the early night (Figure 5A, see also Table 4), suggesting a complex and context-dependent role for PAM neurons in regulation of sleep. Inactivation of PAM neurons (Figure 5B, see also Table 4), inactivation or activation DPM neurons (Figure 5C-D, see also Table 4) in the fed condition produced almost no change in sleep pattern compared to genetic controls before, during and after manipulation. Taken together, these observations again raise the possibility that there may be subtype-specific actions of PAM neurons on DPM neurons. They also demonstrate that their interactions are likely to be context-dependent.

**Figure 5.**
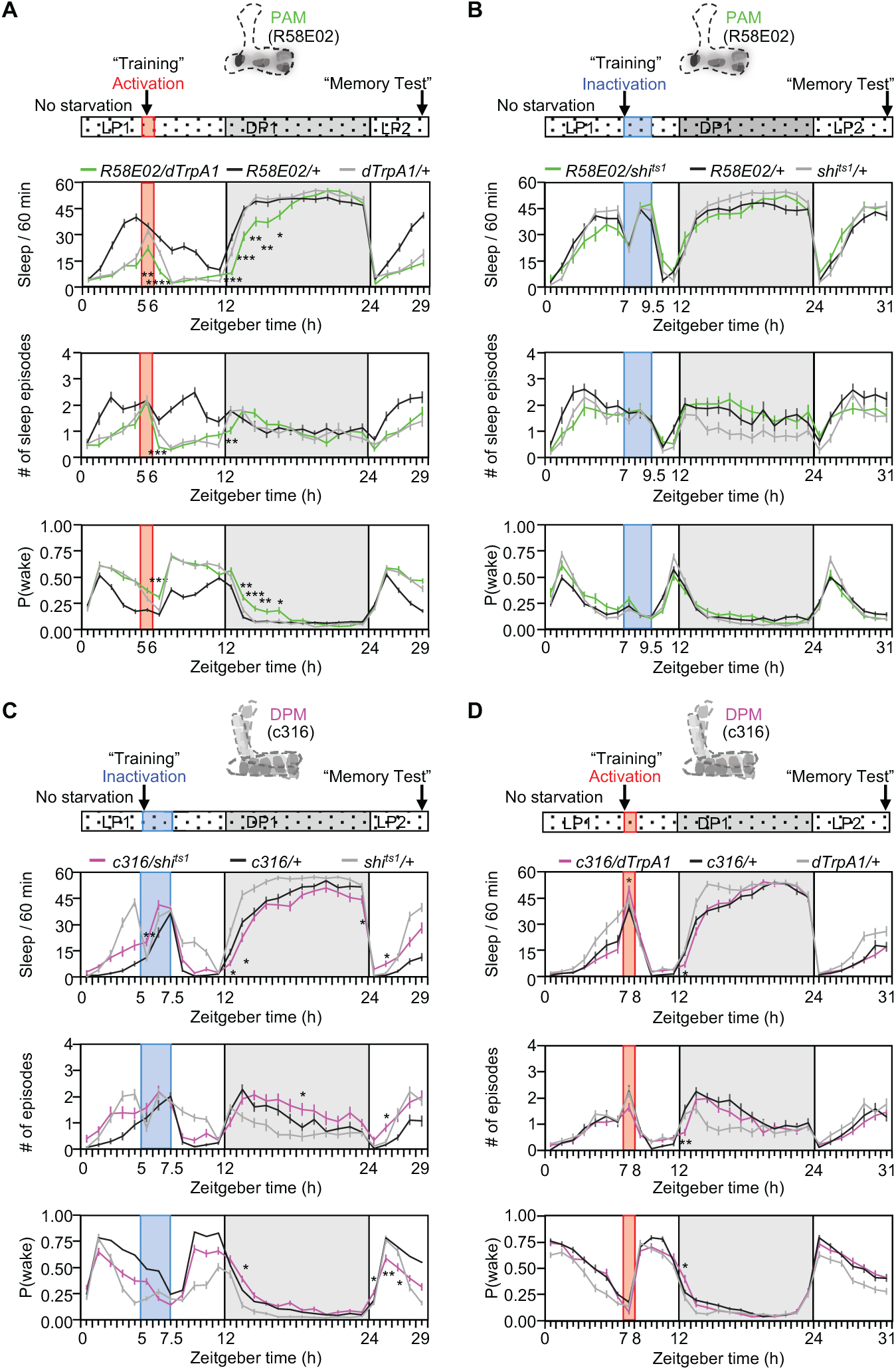
PAM-DPM interactions in sleep regulation are likely to be context-dependent. (A) Activation of PAM neurons without starvation resulted a reduction of sleep at early night, as well as a slight decrease in number of sleep episodes, but an increase in P(wake). n = 39-42. (B) Inactivation of PAM neurons had no effect on sleep without starvation. n = 22-27. (C) Inactivation of DPM neurons labeled by eyeless-GAL80;MB-GAL80;c316-GAL4 without starvation had barely no effects on sleep. n = 35-47. (D) Inactivation of DPM neurons labeled by VT064246-GAL4 had mild effect in sleep at early phase after inactivation without starvation. n = 27-36. Sleep data were presented in 1 h bin. White box: light period; red box: activation period; grey box, dark period; solid fill: starvation; patterned fill: no starvation. See also Table 4.

### The PAM-α1 subset of PAM neurons contributes to linking memory consolidation and sleep

The PAM neurons signaling reward for STM and LTM have been shown to be distinct groups that can be labeled separately by R48B04-GAL4 and R15A04-GAL4, except for overlapping expression in the neurons projecting to the γ5 subdomain (Yamagata et al., 2015). Terminals of R15A04-GAL4 labeling (LTM-PAM) neurons are localized in the β2, α1, β’1, γ5 and pedunculus MB subdomains (Figure 6A). Terminals of R48B04-GAL4 labeling (STM-PAM) neurons are localized in the β’2, β1, and γ1-5 MB subdomains (Figure 6C). To address which subsets of PAM neurons are involved in the process of consolidation, we employed the same single session of appetitive conditioning protocol, activated these neurons after training for 1 h, and examined their 24 h memory. We found that activation of LTM-PAM neurons impaired 24 h memory (Figure 6B, see also Table 1). In contrast, activation of STM-PAM neurons did not affect 24 h memory (Figure 6D, see also Table 1). Thus, LTM-PAM neurons participate in the process of memory consolidation.

**Figure 6.**
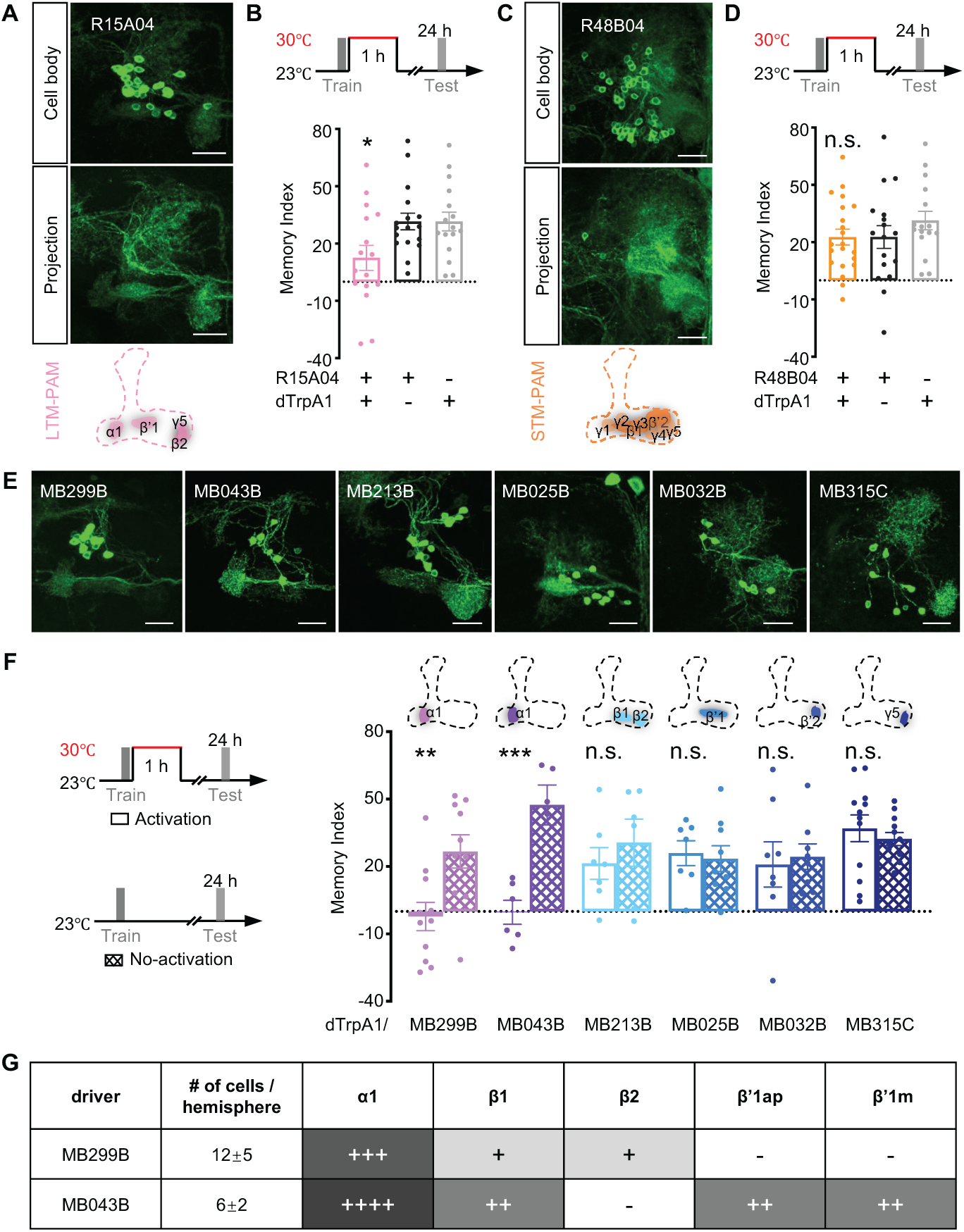
The PAM-α1 subset of PAM neurons contributes to memory consolidation. (A, C) The expression pattern of LTM-PAM and STM-PAM neurons labeled by R15A04-GAL4 and R48B04-GAL4, respectively. Upper panels, cell body; middle panels, projections on the MB; lower panels, schematic of the projections on the horizontal lobes. Scale bar: 20 μm. (B, D) The paradigm of 24 h memory by activation of LTM/STM-PAM neurons for 1 h after training. Activation of R15A04-GAL4+ neurons impairs 24 h memory (B). Activation of R48B04-GAL4+ neurons has no effect on 24 h memory (D). n = 15-21. (E) The expression patterns of specific subtypes of LTM-PAM neurons labeled by split-GAL4 lines: MB299B and MB043B (α1), MB213B (β1/β2), MB025B (β’1), MB032B (β’2), MB315C (γ5). Scale bar: 20 μm. (F) Left panel, activation and no-activation of subtypes of LTM-PAM neurons. Right panel, activation of MB299B and MB043B which project to α1 subdomain impairs LTM compared to the no-activation group. n = 5-12. (G) The differences in cell number/identity between the two drivers of MB299B and MB043B. “+” with greyscales indicated the levels of projections of a driver in MB subdomains; “-” indicated no projections. See also Tables 1 and 2.

To identify which subset(s) of LTM-PAM neurons are involved in the regulation of memory consolidation, we tested split-GAL4 drivers labeling more specific groups. MB213B-, MB025B-, MB032B- and MB315C-GAL4 send processes to β1/β2, β’1, β’2 and γ5 subdomains, respectively (Figure 6E). We trained flies expressing dTrpA1 under control of each of these drivers, with or without 1 h of activation after training, and tested 24 h memory. Only activation of MB299B-GAL4+ and MB043B-GAL4+ neurons impaired 24 h memory compared to their no-activation groups (Figure 6F, see also Table 2). MB299B-GAL4+ (which labels 12±5 PAM neurons/hemisphere) and MB043B-GAL4+ (which labels 6±2 PAM neurons/hemisphere) primarily express in the α1 subdomain (Figure 6G). These data indicate that elevated activity of PAM-α1 neurons during consolidation impedes LTM.

To further probe the requirement of PAM-α1 neurons captured in MB299B- and MB043B-GAL4s in memory consolidation, we either activated them or blocked their output for 1 h or 2.5 h after training, and tested 24 h memory of the experimental flies along with their genetic controls. Activation of both MB299B-GAL4 and MB043B-GAL4 labelled PAM-α1 neurons resulted in an impairment of 24 h memory (Figure 7A-B, see also Table 1). Blockade of MB299B-GAL4+, but not MB043B-GAL4+ neurons, also resulted in a 24 h memory defect (Figure 7C-D, see also Table 1), perhaps reflecting a difference in the number or identity of the PAM-α1 neurons in each line (Figure 6G). These results are consistent with previous studies employing MB299B-GAL4 (Ichinose et al., 2015; Chouhan et al., 2022). Since inactivation of the large group of R58E02+ PAM neurons (Figure 3H) also failed to impair 24 h LTM, these results support the idea that the presence of additional non-α1 PAM neurons might suppress PAM-α1 phenotypes. The connectome catalogs both PAM to PAM connections and recurrent PAM to MBON network loops that might underlie some of the differences between these GAL4 lines.

**Figure 7.**
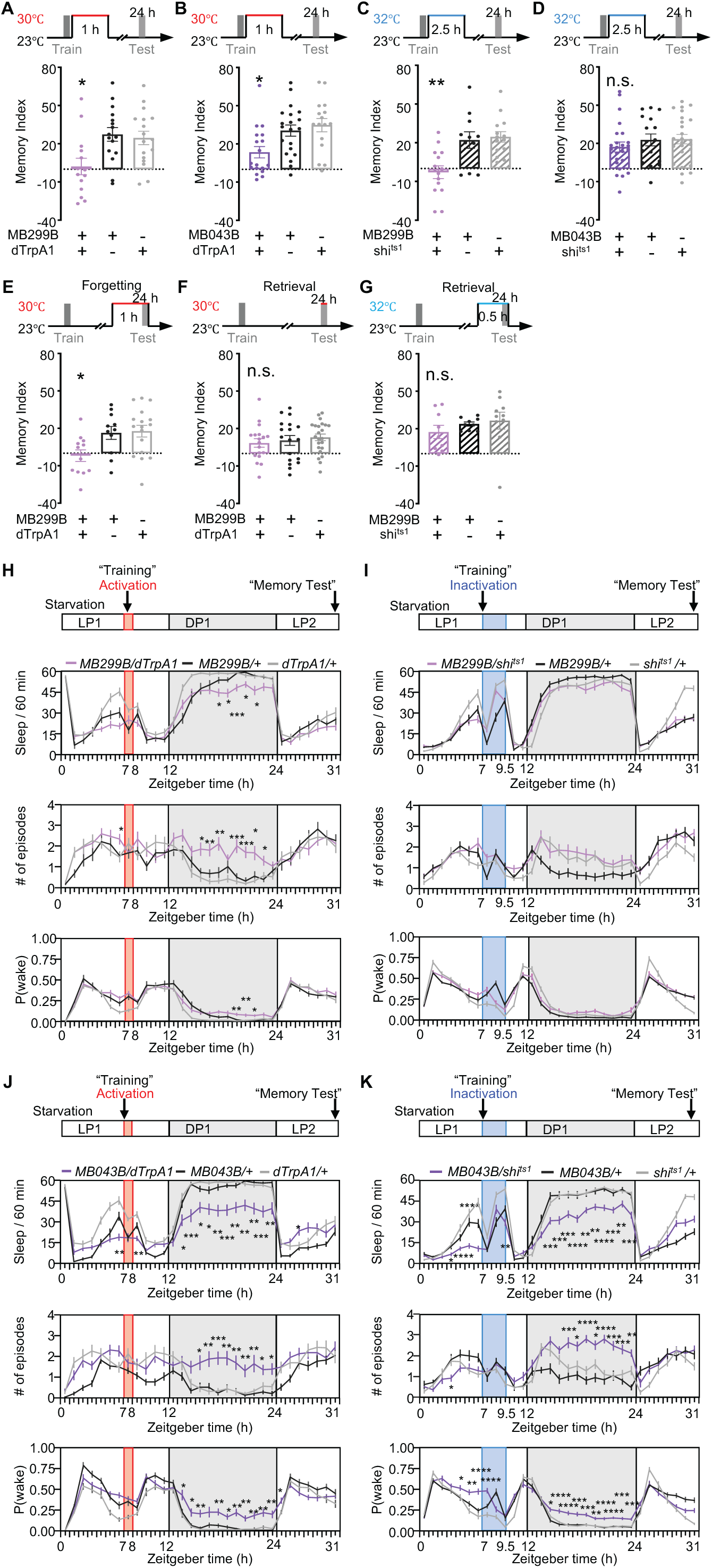
The PAM-α1 subset of PAM neurons contributes to linking memory consolidation and sleep. (A) Flies with activation of MB299B-GAL4+ neurons after training for 1 h exhibited impaired 24 h memory compared to their genetic controls. n = 14-17. (B) Activation of PAM-α1 neurons labeled by MB043B after training for 1 h impaired 24 h memory. n = 16-20. (C) Flies with blockade the output of MB299B-GAL4+ neurons after training for 2.5 h exhibited an impairment of 24 h memory compared to genetic controls. n = 11-14. (D) Inactivation of PAM-α1 neurons labeled by MB043B after training for 2.5 h did not affect 24 h memory. n = 16-25. (E) 1 h activation of PAM-α1 neurons labeled by MB299B prior to and during the test impaired 24 h memory. n = 11-17. (F) Activation of PAM-α1 neurons labeled by MB299B during the test did not affect 24 h memory. n = 18-23. (G) Inhibition of PAM-α1 neurons labeled by MB299B prior to and during the test for 30 min did not affect 24 h memory. n = 9-10. (H) Upper panel: the paradigm for sleep recording under starvation. Lower panel: total sleep, number of sleep episodes, and P(wake) in 1 h bin for activation of PAM-α1 neurons labeled by MB299B. n = 22-30. (I) Inactivation of PAM-α1 neurons labeled by MB299B under starvation had no effects on sleep. n = 33-39. (J) Activation of PAM-α1 neurons labeled by MB043B for 1 h reduced total sleep, increased the number of sleep episodes and P(wake) especially during the dark period. n = 23-30. (K) Inactivation of PAM-α1 neurons labeled by MB043B under starvation significantly reduced total sleep, and increased both the number of sleep episodes and P(wake), especially during the dark period. n = 30-39. See also Tables 1 and 4.

PAM-α1 neurons have been shown to be important for LTM formation, memory retention and updating memory (Huetteroth et al., 2015; Ichinose et al., 2015; Aso and Rubin, 2016; Yamagata et al., 2016). To clarify if their action is exclusive to consolidation or to forgetting or retrieval, we activated and/or blocked PAM-α1 neurons labeled by MB299B-GAL4 prior to and or during the 24 h memory test. Activation of PAM-α1 prior to retrieval impaired 24 h memory (Figure 7E, see also Table 1), but activation and inactivation of PAM-α1 neurons during retrieval did not affect 24 h memory (Figure 7F-G, see also Table 1). These data suggest that PAM-α1 neurons may interact with the active forgetting process during a specific window, but they are not required for retrieval. In aggregate, our findings indicate that the normal activity of PAM-α1 neurons after training is critical to maintain the processing of memory during consolidation.

To investigate whether 24 h memory defects were attributable to disrupted sleep secondary to changes in activity of PAM-α1 neurons, we recorded and analyzed sleep. Consistent with a context-regulated role in linking sleep and memory, activation of PAM-α1 neurons labeled by MB299B and MB043B under starvation conditions reduced sleep amount mainly during the night after training (DP1) (Figure 7H, 7J, upper panel, see also Table 4), reminiscent of DPM suppression. The night after activation of PAM-α1 neurons, flies exhibited a significantly increased number of sleep episodes and P(wake) (Figure 7H, 7J, middle and lower panels, see also Table 4). Inactivation of starved MB299B neurons had little effect on sleep, but inactivation of PAM-α1 cells labeled by MB043B produced a strong reduction and fragmentation of sleep under starvation (Figure 7I, 7K, see also Table 4), again suggesting effects of the differences in cell number/identity between the two drivers. In aggregate, these data demonstrate that both brief increases and brief decreases in activity of PAM-α1 neurons can reduce and fragment sleep. The 24 h memory defect resulting from this change of PAM-α1 activity during consolidation is possibly due to a delayed effect on nighttime sleep, and is completely dependent on internal starvation status since neither activation nor inactivation in fed flies had a major effect on sleep (Supplemental Figure 2A-D, see also Table 4).

### A specific PAM-α1-DPM inhibitory microcircuit binds memory consolidation and sleep via maintaining basal activity post-learning

Using a broadly-expressing driver that captures most if not all PAM neurons, we showed that PAM-DANs form inhibitory synapses onto the DPM neurons (Figure 1). To determine if the connections between the PAM-α1 subset and DPM neurons follow the same rules as the aggregate, we determined the connectivity of this subset. From the EM connectome database (Zheng et al., 2018; Scheffer et al., 2020) and analysis of the connectivity with NeuPrint (Plaza et al., 2022), we found that 14 PAM-α1 neurons of the right hemisphere form 11, 12, 13, 12, 16, 16, 12, 17, 21 and 17 synapses with the right DPM neuron, 5 out of 14 PAM-α1 neurons form a single synapse and one PAM-α1 neuron form 4 synapses with the left DPM neuron, based on the Hemibrain connectome dataset (Figure 8A). To determine the functional connection between PAM-α1 and DPM neurons, we expressed P2X_2_ ATP receptors in PAM-α1 neurons labeled by MB299B and GCaMP6f in the DPM neurons using VT064246-LexA (III), respectively, and confirmed an inhibition from PAM-α1 to DPM neurons by observing a significant decrease in GCaMP signal after application of ATP *in vitro* (Figure 8B, see also Table 3).

**Figure 8.**
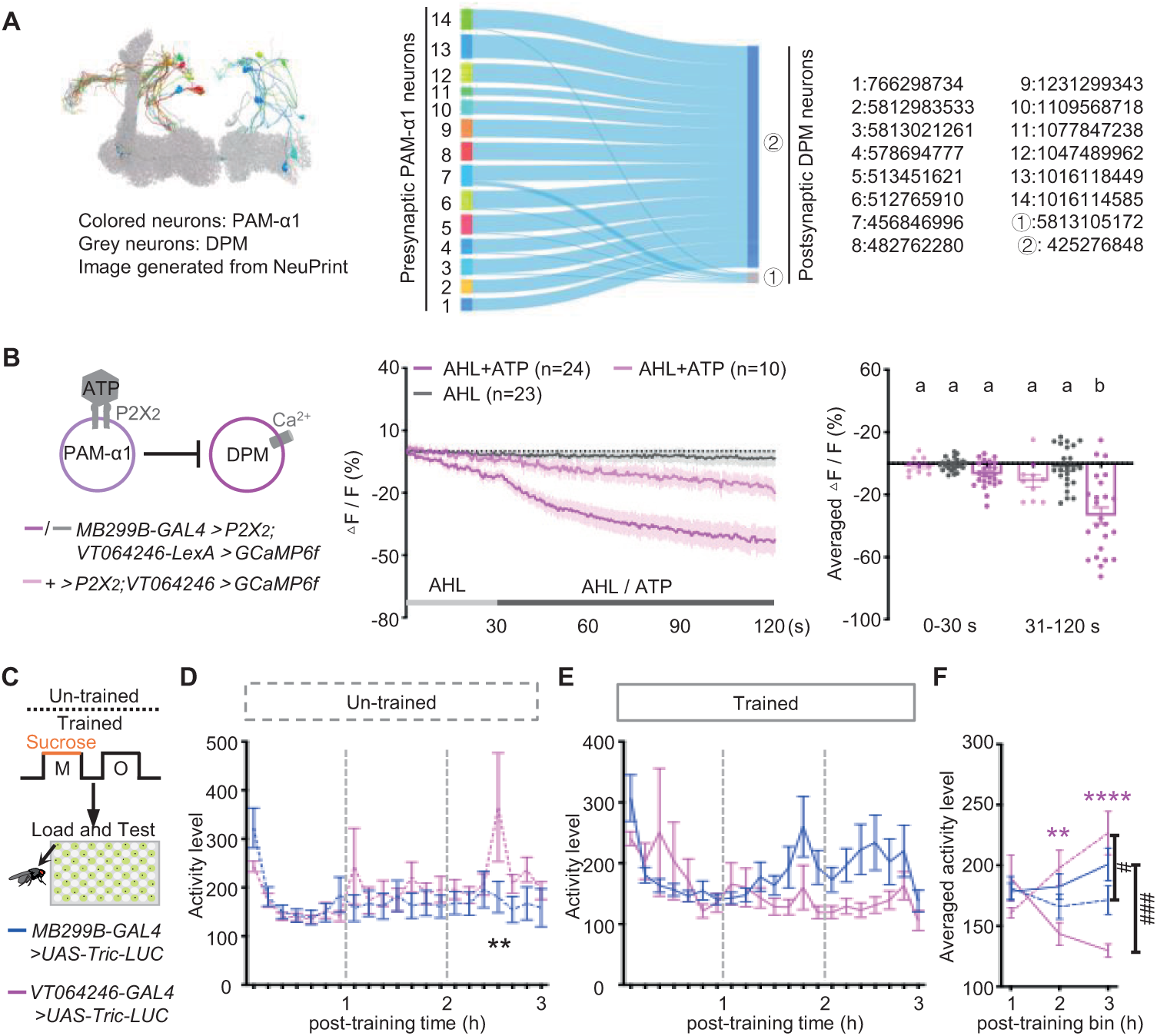
A specific PAM-α1-DPM inhibitory microcircuit maintains dynamic changes in activity post-learning. (A) Characterization of anatomical connections of PAM-α1-DPM microcircuit. Left panel: 3D view of expression patterns of PAM-α1 neurons (colored) and DPM neurons (grey) from NeuPrint. Right panel: synaptic connections between 14 PAM-α1 neurons and 2 DPM neurons with their identity number based on the Hemibrain connectome dataset. (B) Activation of P2X_2_ expressing PAM-α1 neurons labeled by MB299B by application of ATP reduced GCaMP levels in DPM neurons labeled by VT064246-LexA. Experimental genotype under the condition of AHL+ATP, n =24; experimental genotype under the condition of AHL, n = 23; control genotype under the condition of AHL+ATP, n = 10. (C-F) Paradigm for appetitive olfactory training and vehicle un-trained control, and representative neural activity profiles reflected by Tric-LUC expression in drivers labeling PAM-α1 and DPM neurons. Data were continuously recorded for three hours post-training in freely moving flies. (D-E) Dynamic activity changes of PAM-α1 (blue) and DPM (magenta) neurons under un-trained/trained conditions. Dotted lines represent un-trained groups, and solid lines represent trained groups. n = 20-24. (F) Quantification of the averaged changes in PAM-α1 and DPM neuronal activity at each hour post-training. *: significant difference between the DPM neuron groups under un-trained versus trained conditions; #: significant difference between PAM-α1 and DPM neuron groups. See also Table 3.

Given that memory consolidation and sleep are time-dependent processes, to characterize the dynamics of the PAM-DPM circuit over the course of the consolidation window following associative memory training, we performed 3-hour continuous neural activity recording in freely behaving flies. Specifically, we expressed the Tric-LUC reporter transgene, a calcium-responsive tool that harnesses the interaction between calmodulin and its cognate binding peptides to drive rapid luciferase transcription in a calcium-dependent manner (Gao et al., 2015; Guo et al., 2017), in PAM-α1 or DPM neurons. Flies were then subjected to either associative memory training or a no-training control condition, with real-time luciferase levels monitored throughout the recording window (Figure 8C). In the absence of training, both PAM-α1and DPM neurons had a decrease in luciferase signal in the first hour after transfer to monitor tubes, likely reflecting some non-specific acclimation effect (Figure 8D) since this decrease is also seen in trained animals (Figure 8E). Between 1 and 3 h, untrained animals showed similar baseline activity in PAM-α1 and DPM neurons (Figure 8D).

In contrast, associative memory training profoundly reshaped the activity profile of the PAM-DPM microcircuit. During the first hour, activity in PAM-α1 and DPM neurons was similar. During the second hour after training, the activity in the two neurons begins to diverge and becomes quite different as consolidation proceeds (Figure 8E). PAM-α1 neural activity is increased significantly over baseline and DPM neuron activity is robustly decreased (Figure 8E-F). These data suggest that the PAM-α1-DPM microcircuit exhibits delayed neural activity changes following training during the consolidation window, This delay may allow natural memory processes and sleep to be yoked. The fact that both increases and decreases of PAM-α1 activity break this synergy argues that the starvation-dependent resetting of PAM-α1 activity level is a critical feature of the state-dependent function of this circuit.

While it is clear that these two neuron types are connected, it was still possible that they function in sleep and LTM via parallel, independent pathways. If this was true, we would predict that the effects of inactivating both neuron groups at the same time should additively decrease sleep. To determine if this was the case, we simultaneously inactivated these neurons by expressing shi^ts1^ in both cell groups, found sleep reduction, fragmentation, and arousal threshold (P(wake)) statistically the same as when each of them was inactivated separately under starvation (Figure 9A-B, see also Table 4). This result supports a model of a PAM-DPM microcircuit which link sleep and memory consolidation.

**Figure 9.**
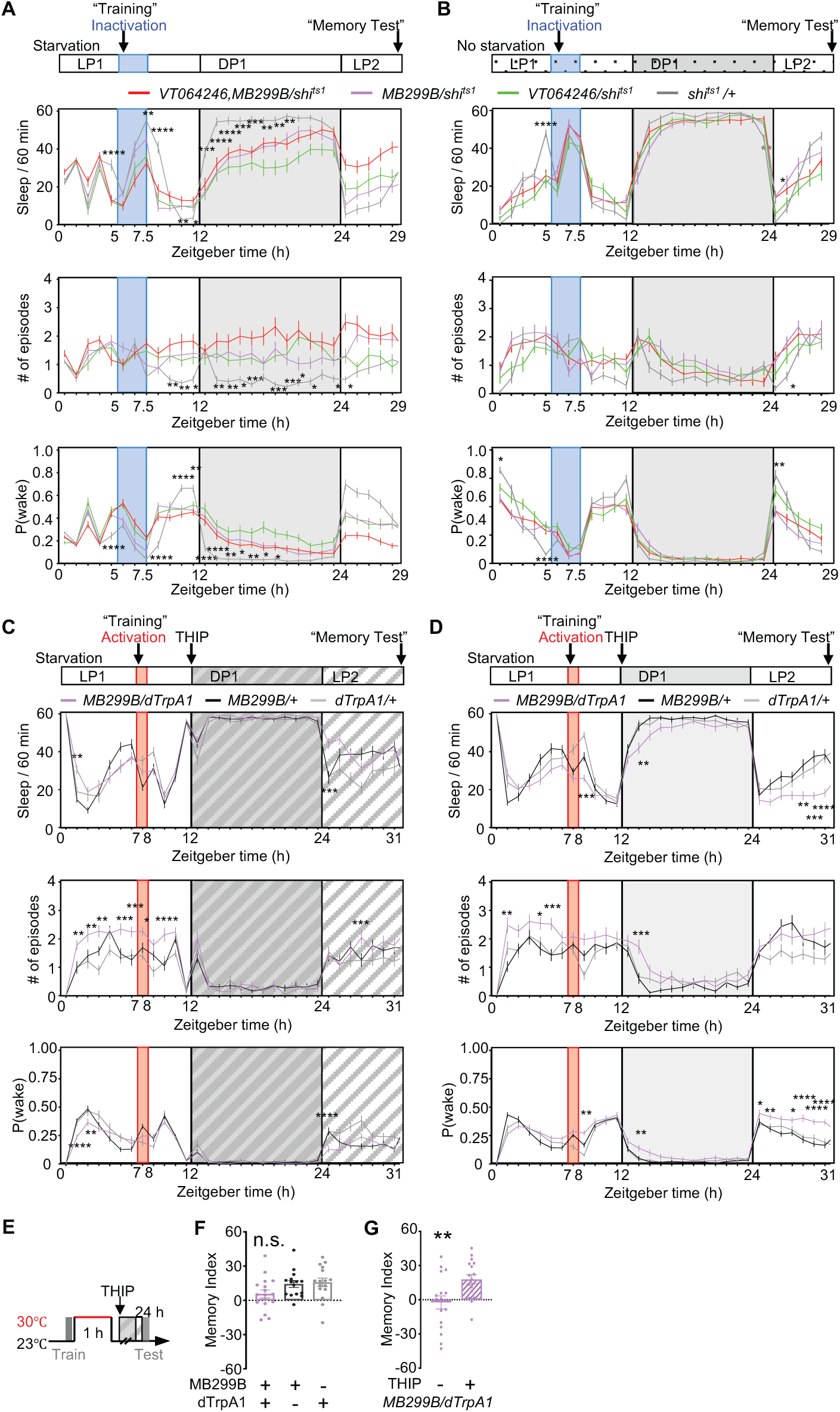
The PAM-α1-DPM inhibitory microcircuit binds memory consolidation and sleep in an internal-state dependent manner. (A) Simultaneous silence of PAM-α1-DPM microcircuit resulted in sleep reduction and fragmentation, increased possibility of waking. n = 36-40. (B) Simultaneous silence of PAM-α1-DPM microcircuit without starvation had little effect on sleep and the number of sleep episodes and P (wake). n = 29-31. (C-D) Sleep profiles of before, during and after activation of PAM-α1 neurons till supposed 24 h memory test with/without 0.1 mM gaboxadol (THIP) treatment. Application of THIP rescued sleep reduction induced by activation of PAM-α1 neurons. n = 38-42 (C), n = 37-44 (D). (E) The paradigm of 24 h memory test with activation of PAM-α1 neurons for 1 h after training. (F-G) THIP rescued 24 h memory deficit induced by 1 h activation of PAM-α1 neurons during consolidation compared to genetic controls (F, n = 15-17) or compared to no THIP group (G, n = 17). See also Tables 1, 2 and 4.

Lastly, to cement the link between sleep disruption by manipulation of this circuit and memory impairment, we employed a pharmacological method. Rescue of the sleep disturbance produced by activating PAM-α1 for an hour with feeding of gaboxadol (also called THIP), a GABA agonist that promotes sleep (Dissel et al., 2015) was sufficient to also rescue memory consolidation, as reflected by memory levels comparable to the genetic controls and significantly better than the no-THIP group (Figure 9C-G, see also Table 1, 2 and 4). Taken together, these results support a mechanistic link between PAM-α1-DPM microcircuit activity in the post-learning time window and both memory consolidation and sleep.

### Dop1R1 expressed in DPM neurons mediates sleep-memory integration

To define the dopamine receptor (DAR) signaling mechanisms underlying DPM neuron activity and its regulatory roles in sleep and memory, we first characterized the DAR expression profile of DPM neurons. Using a double-labeling assay, we detected robust expression of Dop1R1 and Dop1R2 in DPM neurons, whereas no detectable colocalization was observed for DopEcR and Dop2R (Figure 10A). Dop1R1 and Dop1R2 have been reported to stimulate an increase in intracellular cAMP in the presence of DA (Yamamoto and Seto, 2014). To clarify how these DARs mediate the inhibition of DPM neurons, we knocked down Dop1R1 or Dop1R2 in VT046004-GAL4+ neurons and imaged EPAC, an indicator of cAMP levels in DPMs, after application of dopamine (Figure 10B). Dop1R1 RNAi and Dop1R2 RNAi both efficiently reduced the relative mRNA levels of their target gene (Supplemental Figure 3A-B). Knockdown of Dop1R1 receptors in DPM neurons significantly suppressed the dopamine-induced increase in cAMP (visualized as a decrease in EPAC FRET), whereas knockdown of Dop1R2 trended to further increasing cAMP levels (Figure 10B, see also Table 3). These data confirm that Gαs-coupled Dop1R1 is the primary receptor mediating DA-dependent cAMP elevation in DPM neurons. To further identify the DARs responsible for transducing the DA-induced inhibitory effect on DPM neural activity, we quantified GCaMP signals in DPM neurons with targeted knockdown of individual DARs. Knockdown of either Dop1R1 or Dop1R2 failed to abolish the DA-induced Ca^2+^ decrease (Supplemental Figure 3C-E). Because the experiments were done in the presence of TTX to eliminate action potential-dependent release of substances from other cells in the circuit, this suggests that either both receptors cooperate to modulate DPM neural activity, or that there is another (undetected) DAR expressed in DPMs. It is also possible that since RNAi knockdown is never complete, residual Dop1R1/R2 is sufficient to mostly support the calcium-reducing effects of DA, but not the efficient coupling to adenylate cyclase if the downstream pathways are differentially sensitive.

**Figure 10.**
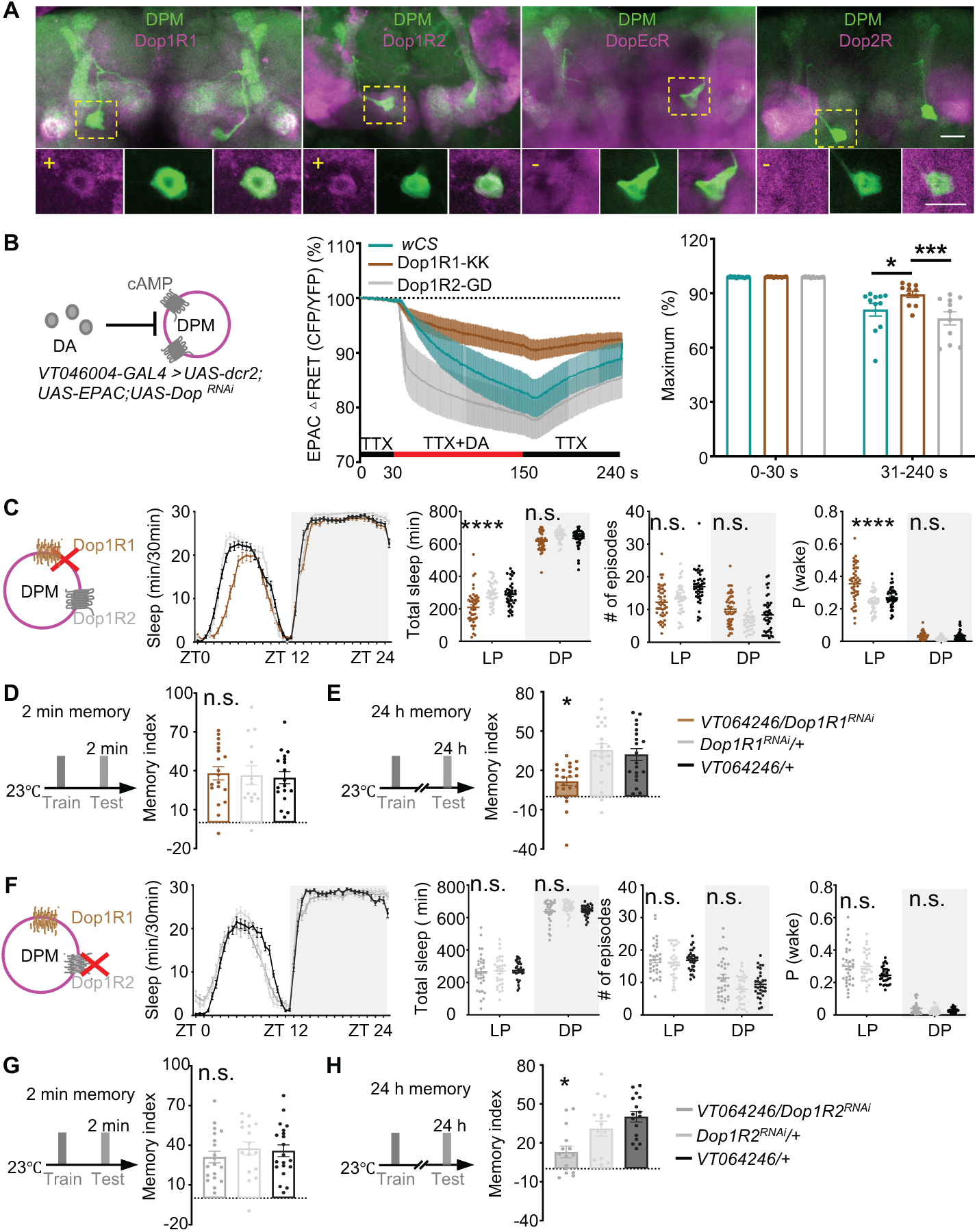
Dop1R1 expressed in DPM neurons mediates sleep-memory integration. (A) Colocalization of DPM neurons with distinct dopamine receptor subtypes. DPM neurons labeled by VT064246-LexA were visualized by GFP (green), and dopamine receptors labeled by Dop1R1-GAL4, Dop1R2-GAL4, DopEcR-GAL4 and Dop2R-GAL4 were visualized by RFP (magenta). The yellow dashed box outlines the cell body of one DPM neuron, with the upper and lower panels on the right showing the split channels for DPM neurons and the corresponding dopamine receptor subtype (white arrow), respectively. Scale bar: 20 μm. (B) Averaged cAMP traces (middle panel) and quantification (right panel) of DPM neurons in response to DA with the presence of 2.0 mM TTX with or without Dop1R1 or Dop1R2 knockdown. n = 11 for all groups. (C) Knock down Dop1R1 on DPM neurons decreased the total amount of daytime sleep and increased P(wake). n = 40-44. (D-E) Knock down Dop1R1 on DPM neurons had no effect on 2min, but impaired 24 memory. n = 14-19 (D), n = 20-22 (E). (F) Knock down Dop1R2 on DPM neurons, had no effect on sleep. n = 31-32. (G-H) Knock down Dop1R2 on DPM neurons had no effect on 2min, but impaired 24 memory. n = 17-20 (G), n = 15-16 (H). See also Tables 1 and 3.

Finally, to dissect the specific contributions of DARs in DPM neurons to sleep and/or memory regulation, we performed sleep monitoring and memory assays in animals with DPM-specific knockdown of distinct DARs (Figure 10C-H). Knockdown of Dop1R1 in DPM neurons resulted in statistically significant sleep reduction, decreased arousal threshold, and impaired 24 h LTM memory with intact learning (Figure 10C-E). In contrast, knockdown Dop1R2 in DPM neuron selectively impaired 24 h LTM memory with no effect on sleep and learning (Figure 10F-H). Collectively, these findings demonstrate that coupling sleep and LTM requires Dop1R1 in DPM neurons through the modulation of both cAMP signaling and neuronal activity, while Dop1R2 specifically mediates LTM regulation, likely through modulating DPM neural activity alone.

### MBON-α1 neurons participate in regulation of both memory consolidation and sleep

PAM-α1 has previously been demonstrated to drive appetitive LTM formation and consolidation via a recurrent loop with MBON-α1 (Ichinose et al., 2015). To investigate whether MBON-α1 also participates in sleep regulation, we activated or inactivated MBON-α1 neurons under both starvation and non-starvation conditions (Figure 11A-D). Our results revealed that inhibition of MBON-α1 under both starvation and non-starvation conditions resulted in a significant reduction in sleep and a reduced arousal threshold (Figure 11B, D), suggesting that MBON-α1 participates in regulating sleep in a state-independent manner. However, no significant changes were observed upon activation of MBON-α1 neurons (Figure 11A, C). Moreover, consistent with previous finding, inhibition of MBON-α1 during the memory consolidation phase significantly impaired 24 h LTM (Figure 11F). These results indicate that MBON-α1mediated sleep is necessary for effective memory consolidation. Notably, while activation of MBON-α1 during consolidation phase similarly impaired LTM (Figure 11E), this manipulation did not alter the sleep profile, suggesting a dissociation between MBON-α1’s mechanistic roles in sleep regulation and LTM processing. Taken together, these findings reveal a dedicated hierarchical, modular regulatory network that mediates sleep-LTM coupling via an activity-dependent mechanism. Within this network, activation of PAM-α1 acts as an upstream modulator to inhibit the activity of DPM, a downstream integrative hub that coordinates the execution of sleep and memory processes, which is modulated by the recurrent positive feedback loop of PAM-α1-MBON-α1.

**Figure 11.**
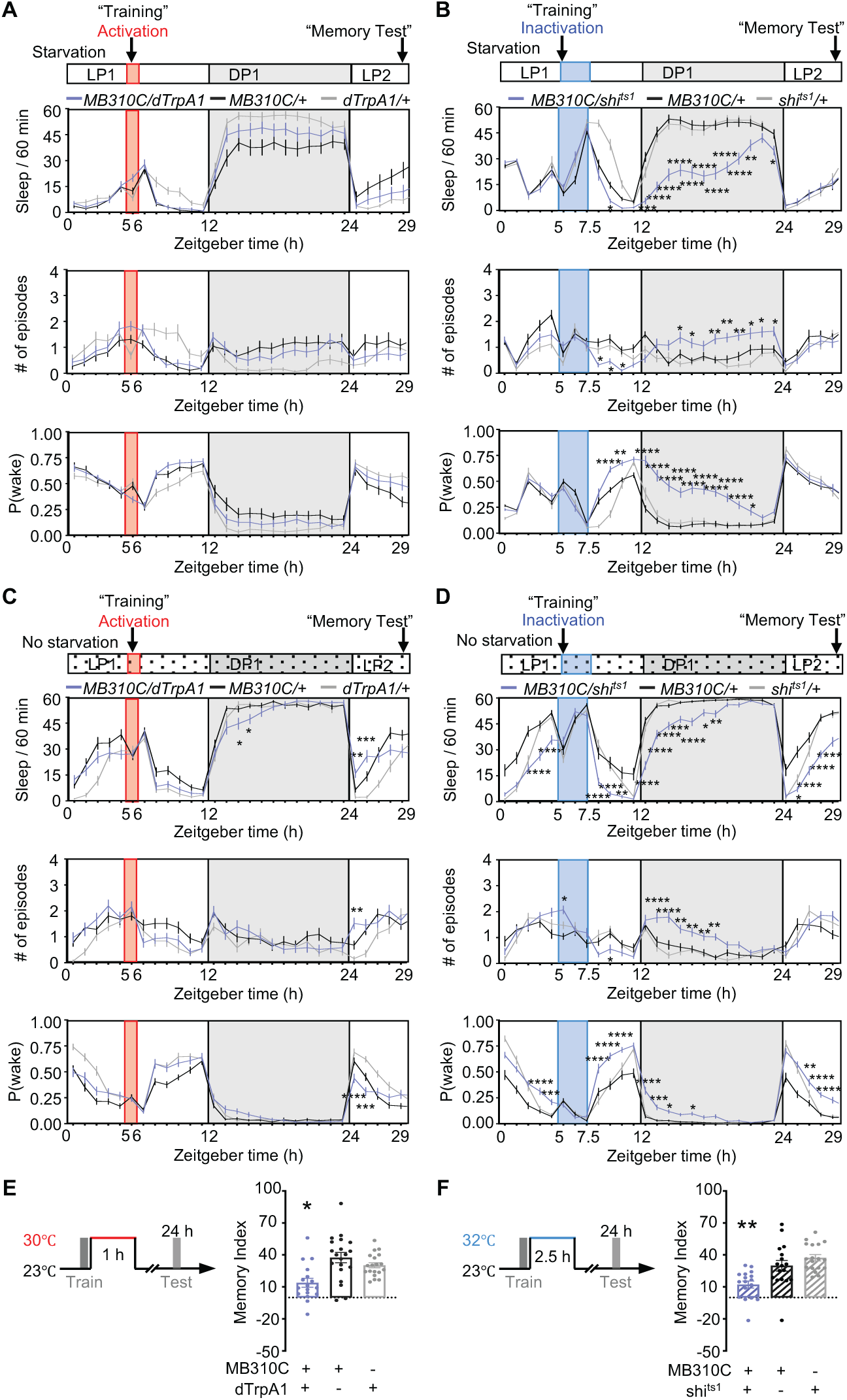
MBON-α1 neurons regulate sleep irrespective of starvation state, and their activity disruption during memory consolidation impairs LTM. (A, C) Activation of MBON-α1 neurons labeled by MB310C under starvation (A, n = 26-32) or no starvation (C, n = 31 for all groups) condition had no effects on sleep. (B, D) Inactivation of MBON-α1 neurons for 2.5 h significantly reduced total sleep, increased the number of sleep episodes and P(wake) especially during the dark period under starvation (B, n = 37-44) and no starvation (D, n = 33-43) condition. (E) Flies with activation of MBON-α1 neurons after training for 1 h exhibited impaired 24 h memory compared to their genetic controls. n = 17-20. (F) Flies with blockade the output of MBON-α1 neurons after training for 2.5 h exhibited an impairment of 24 h memory compared to their genetic controls. n = 17-18. See also Tables 1 and 4.

## Discussion

PAM-DANs have been known to be important for formation of short-term and long-term memories, and DPM neurons for memory consolidation and sleep. In the present study as summarized in Figure 12, we find that these two types of MB extrinsic neurons are connected by inhibitory synapses and play an essential role in bridging LTM stabilization via dynamic sleep change. We demonstrate that a brief disruption of this circuit, in particular of a subset of PAM neurons (PAM-α1) during the critical memory consolidation window, results in an impairment of LTM. Worth noticing, sleep loss and fragmentation occur hours after the brief disruption of this circuit when the flies are starved. Thus, maintenance of basal activity of PAM-α1-DPM microcircuit during memory consolidation has a profound effect on LTM, through influencing sleep when the animal is experiencing a certain internal state. Distinct signaling cascades may be recruited via different dopamine receptors in DPM neurons to integrate or segregate sleep and memory processes. Additional parallel circuits further contribute to linking and specifying the regulatory logic of these processes.

**Figure 12.**
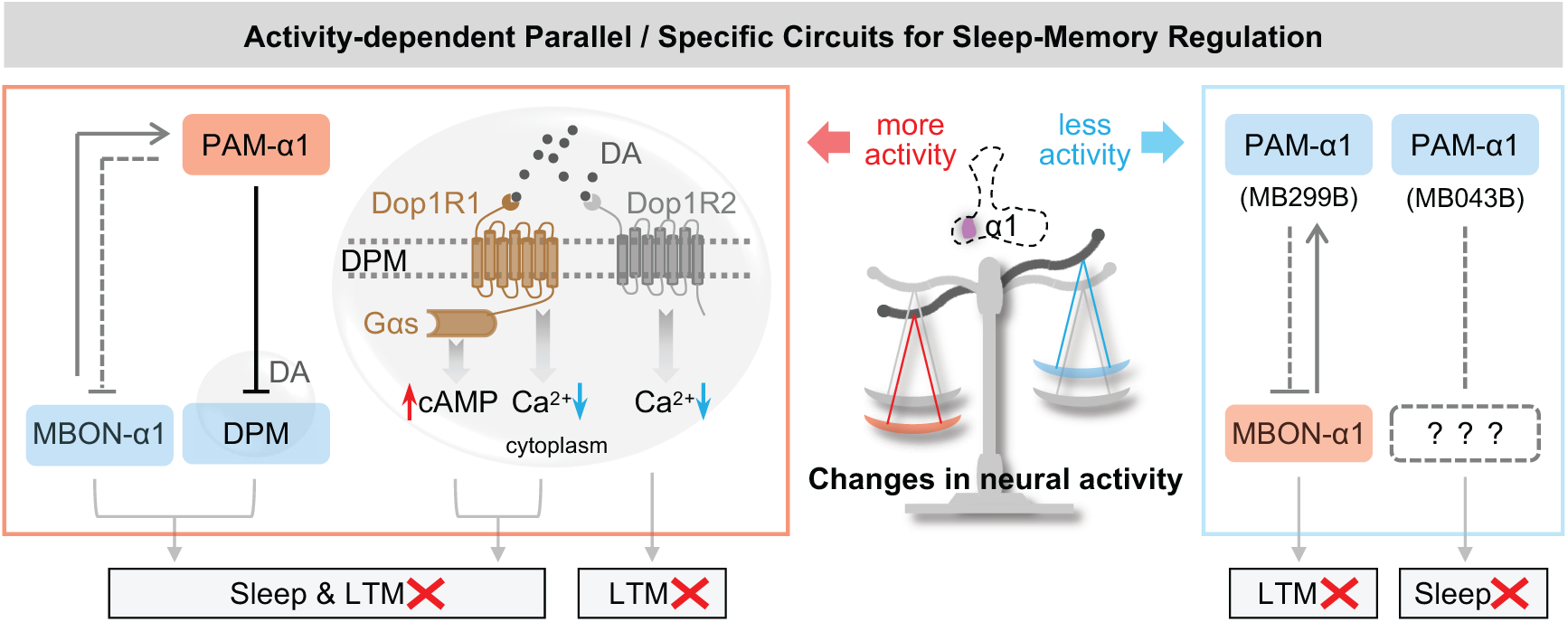
Schematic of activity-dependent parallel and specific regulatory circuits for sleep-memory regulation. Increased activity of PAM-α1 neurons strongly inhibits DPM, leading to disrupted sleep and impaired LTM. MBON-α1 likely acts as a parallel circuit relative to PAM-α1-DPM pathway, interfering with the coupling between sleep and memory. DA signaling through Dop1R1 receptors triggers an increase in cAMP and a decrease in Ca²⁺, disrupting both sleep and LTM. By contrast, signaling via Dop1R2 receptors only reduces Ca²⁺ levels, specifically impairing LTM. However, inactivated PAM-α1 may engage different downstream pathways to regulate sleep and memory independently. Inactivation of PAM-α1 neurons labeled by MB299B and MB043B leads to LTM deficits and sleep disturbance, respectively. Combined with the lack of sleep phenotype upon MBON-α1 activation, reduced PAM-α1 activity and elevated MBON-α1 activity participate in LTM regulation. Nevertheless, which downstream targets of MB043B-labeled PAM-α1 mediate sleep regulation remains an open question.

### Mushroom body DANs are not just a valence signal

DANs can have multiple roles at different times in the memory process. PAM-α1 is one of many cell types of DANs that required for memory formation (Yamagata et al., 2015; Yamagata et al., 2021). We find that not only suppressed activity but also elevated activity of PAM-α1 neurons right after training can disrupt memory consolidation, supporting a role in consolidation (Ichinose et al., 2015). PAM-α1 neurons form a recurrent loop with MBON-α1 neurons (Aso et al., 2014b; Ichinose et al., 2015), and the recurrent DAN/MBON networks are likely to be important in many aspects of memory processing. Several subsets of DANs have been shown to be involved in forgetting, PPL1-γ1pedc (MP1) and PPL1-γ2α’1 (MV1) for permanent forgetting (Berry et al., 2012), and PPL1-α2α’2 for transient forgetting (Sabandal et al., 2021). In addition, Aso and Rubin (2016) demonstrated that writing and updating memories recruit different rules across distinct MB compartments. Memory consolidation is a dynamic time window from hours after learning but may continue for weeks (Roesler and McGaugh, 2010), and many studies have suggested that sleep deprivation is important during memory consolidation but not retrieval (Krishnan et al., 2016; Manassero et al., 2022). Our results do not rule out a role for PAM-α1 in active forgetting processes, but it is clearly not required for retrieval (Figure 7E-G), adding a new layer of complexity of multiple functions within a single type of neurons.

### Sleep interacts with mushroom body DANs in multiple ways

Sleep is thought to be indispensable to memory consolidation process, converting initially unstable memory into stable long-term memory during sleep (Diekelmann and Born, 2010; Klinzing et al., 2019; Girardeau and Lopes-Dos-Santos, 2021; Yan et al., 2025). Besides the amount of sleep, accumulating evidence has suggested that sleep structure can influence memory processing (Rolls et al., 2011; Lipinska and Thomas, 2019; Liu et al., 2019; Kjaerby et al., 2022). In *Drosophila*, sleep structure can be evaluated by the number of sleep episodes as well as P(wake). The number of sleep episodes reflects the frequency of awakenings, and P(wake) is the probability of transitioning from a sleep to an awake state, reflecting the threshold for arousal (Liu et al., 2019; Wiggin et al., 2020). In the present study, we found that a brief disruption of neural activity of a specific microcircuit resulted in reduced and fragmented sleep, with a consequent impairment of LTM, providing a new circuit basis for the linkage between these two processes.

DPM neurons process memory consolidation via promoting sleep, likely via inhibition of wake-promoting MB-α’β’ neurons (Donlea, 2011; Haynes et al., 2015), and this circuit also seems particularly important for consolidation (Krashes et al., 2007). DPM neurons have been recently endowed with new functions, such as a gatekeeper for intrinsic coincidence time windows, and a bridge for mediating multisensory stimulus binding (Okray et al., 2023; Zeng et al., 2023), further revealing a richness in the compartmentalized specificity and plasticity of microcircuits. The PAM-α1-DPM microcircuit in the present work unlocks an upstream modulation of MB circuits for linking memory and sleep.

There are “forgetting” DANs (such as PPL1-γ2α’1) whose activity has been shown to be suppressed by sleep (Berry et al., 2015). This PPL1-γ2α’1 subset, and other MB DANs (including PAM-α1) have been identified as wake promoting neurons (Donlea, 2011; Sitaraman et al., 2015; Driscoll et al., 2021). Maybe this is one of the ways the recurrent network processes sleep. Perhaps the activation of PAM-α1 decreases sleep and that allows PPL1-γ2α’1 to cause forgetting. This is a model that could be tested in the future. In spite of the fact that MB circuits have been elegantly investigated, the leap from the microcircuit level to the systems scale remains daunting, and how these circuits enable consolidation during sleep still needs further investigation.

### Internal state dictates circuit configuration for consolidation

Internal state, like starvation, gates many processes through molecular mediators and alternative neural circuits (Keene et al., 2010; McDonald, 2010; Lin et al., 2019; Yurgel et al., 2019). DANs contribute robustly to coordination of state-dependent behaviors, such as facilitating starvation-dependent sugar memory, and mediating starvation-driven food seeking behavior (Krashes et al., 2009; Tsao et al., 2018; Senapati et al., 2019). Driven by context, distinct subsets of DANs exhibit suppressed or enhanced neuronal responses facilitating integration of motivational signals and modulating output behaviors (Liu et al., 2017; Jovanoski et al., 2023). Our present study characterizes another subset of DANs that respond to starvation with an increase in neural activity to provide a context-dependent regulation of behavior.

It is well established that starvation suppresses sleep (MacFadyen, 1973; Thimgan et al., 2010; Melnattur and Shaw, 2019; Keene et al., 2010; He et al., 2020; Oh and Suh, 2023), and our results are consistent with these previous findings: all genotypes exhibited less sleep under starvation than under fed conditions (i.e. Figures 4A, B, 5A, B and 11). With normal appetitive memory training, starvation-induced sleep loss does not necessarily impair memory processing (Thimgan et al., 2010; Chouhan et al., 2021). PAM-α1 neuronal activity is higher in starved, trained flies than in fed or untrained flies (data not shown), suggesting that these neurons act as a critical node for integrating internal motivational and arousal states, as well as conveying positive valance for the normal appetitive memory process, independently of starvation-induced sleep loss. While DPM neurons are less sensitive to starvation, they still exhibit training-induced elevated activity (data not shown), indicating coherent responsiveness to upstream signaling. In the present study, we found that under fed conditions, sleep remained intact even when large changes in neural activity occurred within the PAM(-α1)-DPM circuit; in contrast, under starvation conditions, significant sleep reduction and fragmentation were observed. These observations indicate that starvation may trigger a transition from a physiologically normal brain state to an unstable, abnormally active state, which consequently elicits negative behavioral outputs.

Sleep serves memory consolidation (Brodt et al., 2023; Chandra et al., 2023). Paradoxically, flies need to be starved to express appetitive LTM. But it can be interpretated in a way that the animal’s state during consolidation can recruit different circuitry at both the KC level and the DAN level (Chouhan et al., 2021; Chouhan and Sehgal, 2022). Our results provides yet another example of this. Suppressed but not increased activity of MBON-α1 negatively impacts on sleep no matter the animal is starved or not. However, disrupted activity of MBON-α1 during memory consolidation phase impairs appetitive LTM. The activity of the microcircuit we have identified is likely to be key to understanding how sleep need is taken out of the loop during starvation.

### Activity changes trigger distinct circuits controlling sleep and memory consolidation

An additional interesting feature of the role of PAM-α1 neurons in consolidation of appetitive LTM is that it is very sensitive to the level of activity within this neuron group (Figure 12). Both increases and decreases in activity can block LTM and fragment sleep. The inhibitory nature of the connectivity between PAM-α1 and DPM provides a plausible explanation for the effects of overactivation of PAM-α1: strongly blocking DPM activity would suppress both sleep and consolidation (Waddell, 2000; Keene et al., 2004; Yu et al., 2005; Krashes and Waddell, 2008; Haynes et al., 2015). Furthermore, MBON-α1, provides an additional component which serves as a parallel pathway to suppress sleep and impair LTM modulated by this PAM-α1-MBON-α1 recurrent feedback loop. PAM-α1 may forms an inhibitory pathway on MBON-α1, supported by the same sleep and memory outcomes upon activation of PAM-α1 and inactivation of MBON-α1. How inactivation of PAM-α1 produces a similar ‘DPM-inhibition-like’ state is more difficult to explain with current data. However, according to the observations of specified modulation on memory or sleep, PAM-α1 relieves its inhibitory control over MBON-α1, leading to MBON-α1 activation which has no effect on sleep; MBON-α1 then functions as a signal amplifier that further exacerbates the reduced activity of PAM-α1, specifically resulting in LTM impairment. Whether this is a direct effect of decreasing the level of stimulation of dopamine receptors that occurs under starvation conditions when PAM-α1 activity increases or if it is an indirect effect mediated by an interneuron will require further investigation. In another case, inactivation of PAM-α1, possibly non-PAM-α1 neurons labeled by MB043B, specifically contributes to the regulation of sleep. In addition to the α1 subdomain, MB043B also projects to several other compartments, but which downstream targets are recruited to mediate sleep regulation remains an open question. Sleep and memory are highly intertwined within this circuit, where distinct neuronal populations exhibit specialized yet interdependent functional roles, with overlapping and divergent regulatory contributions to sleep and LTM. The inherent complexity of this regulatory network thus merits further dedicated investigation in future studies.

## Materials and methods

### Animals

All flies were reared on standard cornmeal food at 25 ℃ and 60% relative humidity on a 12 h/12 h light-dark cycle. Flies expressing dTrpA1 and shi^ts1^ used in temperature-shift experiments were raised at 23 ℃. Flies for memory and sleep experiments allowed to freely mate. 5-10 days old and 5-7 days old adult flies were used at the start of memory and sleep experiments, respectively. The following stocks were obtained from the Bloomington Drosophila stock center (BDSC, Indiana University): *w;;R58E02-GAL4* (70226), *w;;R15A04-GAL4* (48671), *w;;R48B04-GAL4* (50347), *MB025B-GAL4* (68299), *MB213B-GAL4* (68273), *MB032B-GAL4* (68251), *MB315C-GAL4* (68316), *MB299B-GAL4* (68310), *MB043B-GAL4* (68304), *elav-GAL4* (458), *UAS-mCD8::GFP; Pin/Cyo* (5136), *w;UAS-dTrpA1* (26263), *w;;UAS-shibire^ts1^* (44222), *w;R58E02-LexA* (52740), *eyeless-GAL80;MB-GAL80/Cyo;c316-GAL4/TM6B* (30380), *UAS-CD8::RFP,LexAop-CD8::GFP*(302229), *LexAop-nSyb:spGFP1-10,UAS-CD4:spGFP11;MKRS/TM6B* (64315), *UAS-EPAC* (78802), *UAS-dcr2* (24650), *LexAop-P2X_2_* (76030), *UAS-GCaMP6f* (42747), *UAS-CD4-spGFP1-10* (93016)*, LexAop-CD4-spGFP11* (93019), *UAS-Dop1R2^RNAi^* (51423) (Ni et al., 2011), *UAS-P2X_2_* (91223) and *LexAop-GCaMP6f* (44277), *UAS-CRTC::GFP/CyO;UAS-mCD8::mCherry-T2A-nls::LacZ in VK00005/TM2* (99656). *VT064246-GAL4/TM2* (v204311), *UAS-Dop1R1^RNAi^* (v107058) (Scaplen et al., 2020), *UAS-Dop1R2^RNAi^* (v3392) (Dietzl et al., 2007) were ordered from Vienna Drosophila Resource Center (VDRC, Vienna, Austria). *VT046004-GAL4* was one of Vienna Tile GAL4 lines, currently available at Korea *Drosophila* Resource Center (12772). The following lines have been previously described: *trans-Tango,QUAS-FLP;LexAop-FRT-RFP-FRT-GFP* (Sun et al., 2022). *w*;;Dop1R1-GAL4 and w*;;Dop1R2-GAL4* was kindly provided by Prof. J. W. Kim, and *Dop2R-KI-GAL4* was kindly provided by Prof. Yufeng Pan. For behavioral experiments, *w^CS^,UAS-dTrpA1* or *w;;UAS*-*shi^ts1^* virgin female flies were crossed to male flies of GAL4 lines. Virgin female *w^CS^* flies were crossed with GAL4 or UAS parental lines as genetic controls.

The *VT064246-LexA* transgenic lines were based on VT064246-GAL4. The 2296 bp promotor fragment was amplified from genomic DNA of wild-type flies using the same primers as used for VT064246-GAL4 (forward primer: 5’-AATACGAGAGCG CTCACTAC-3’ and reverse primer: 5’-TCTTATGGACGGCGAGGAG-3’). This fragment was then cloned into enhancer-LexA vector (Fungene Biotech, http://www.fungene.tech) using NotI and AscI (Thermo Fisher Scientific) restriction sites. The sequence was confirmed by sequencing using primers as follow: VT064246-3F: 5’-AATACGAGAGCGCTCACTAC-3’ and reverse lexA-5R: 5’-TCTTATGGACGGCGAGGAG-3’. The plasmids were inserted into attP40 (25C6 of chromosome 2) site and attP2 (68A4 of chromosome 3) site using attp PhiC31 mediated recombination (Fungene Biotech).

### Memory assays

F1 generation of mixed genders in a group were trained and tested together. All training was carried out under the condition of dim red light and all tests were performed in darkness.

4-methylcyclohexanol (MCH) and 3-octanol (OCT) diluted in mineral oil to 10% were used as conditioned odors. Flies were trained with/without starvation. Flies were starved for 30-46 h when each genotype exhibits a mortality rate of 10%-20% when flies started to be tested.

For all 24 h sucrose-odor memory, a single training session of sucrose paired with an odor for 2 min was employed (Figure 3A). Blockade of the output of neurons by shi^ts1^ during the training phase, flies were placed at 32 °C 2.5 h prior to training. Manipulation of the activity of target neurons during memory consolidation phase included: 1) activation for 1 h by dTrpA1 at 30 ℃; and 2) blockade for 2.5 h by shi^ts1^ at 32 °C.

A performance Index (PI) was calculated by the difference between flies chose the odor paired with an unconditioned stimulus (US: high temperature or sucrose) and flies chose the other odor without US, divided by the total number of flies. A learning index (LI) was then calculated as the averaged two PIs from two reciprocal groups multiplied by 100. Half of LIs were from the first odor paired with US, and the other half of LIs from the second odor paired with US to minimize the bias of the conditioning order.

### Sleep assays

Regular sleep tubes contained food of 2% agar with 5% sucrose, and the starvation sleep tubes contained only 2% agar. A female fly was anesthetized with CO_2_, and then loaded into a 65 mm × 5 mm glass tube which was then placed into a *Drosophila* Activity Monitoring (DAM2, Trikinetics, Waltham) board. DAM boards were then loaded into an incubator with a 12 h:12 h light-dark cycle at 60% relative humidity, and connected to DAM system. After 2 days of entrainment, flies were recorded for another 2 days for sleep analysis. Starvation time of each genotype was consistent as in the memory tests. Flies were transferred to starvation sleep tubes between Zeitgeber Time (ZT) 23 and ZT2. The environment temperature was increased to 30 ℃ for 1 h to activate the neurons expressing dTrpA1, or to 32 ℃ for 2.5 h to inactivate the neurons expressing shi^ts1^ at ZT5/7, and then shifted the temperature back to 23 ℃ for another 24 h. Sleep parameters including total sleep, number of sleep episodes, and P(wake) were analyzed in 1 h bin. Sleep files acquired from DAM system were processed with DAMFileScan113X (TriKinetics Inc, Waltham, MA), and SCAMP 2019v2 using MATLAB_R2021b (MathWorks, Natick, MA). The probability of transition from a sleep to an awake state P(wake) was analyzed as previously described (Wiggin et al., 2020). The MATLAB scripts for analysis is accessed in GitHub at https://github.com/Griffith-Lab/fLy_Sleep_Probability.

For sleep with an application of gaboxadol hydrochloride (known as THIP, Cat# 85118-33-8, Sigma-Aldrich), flies were activated for 1 h at ZT7 in 2% agar starvation tubes, and then transferred to sleep tubes containing THIP (0.1mg/ml) in 2% agar at ZT12 for 20 h.

### Immunohistochemistry

Both male and female flies aged 5-7 days were used for immunostaining. Fly brains were dissected in Schneider’s Drosophila Medium (1×) (Cat# 21720-024, Gibco). Dissected brains were fixed in 4% paraformaldehyde (PFA, Cat# 50-00-0, GHTECH) for 20 min at room temperature, and then washed in 1× phosphate buffered saline (PBS) containing 0.5% Triton X-100 (PBS-T) at 4 ℃ for 10 min for 3 times. After that, the brains were incubated in primary antibodies with 5% normal goat serum (Cat# 0005-000-121, Jackson) at 4 ℃ overnight. Primary antibodies used for visualization of GFP and RFP (including restricted *trans*-Tango) were: chicken anti-GFP (1:200, Cat# 13970, Abcam) and rabbit anti-DsRed (1:200, Cat# 632496, Clontech); primary antibodies used to detect GRASP signals was mouse monoclonal anti-GFP (1:200, Cat# 11814460001, Roche Applied Science). Samples were then washed in PBS-T at 4 ℃ for 10 min for 3 times, incubated in secondary antibodies at 4 ℃ overnight. Secondary antibodies (1:200) included: Alexa Fluor 488 (goat anti-chicken, Cat# A11039, Invitrogen), Alexa 568 (goat anti-rabbit, Cat# A11011, Invitrogen), Alexa Fluor 633 (goat anti-mouse, Cat# A-21052, Invitrogen). Samples were washed in PBS-T at 4 ℃ for 10 min for 3 times, and mounted with Vectashield (Cat# H-1000, VECTOR). Images were acquired with a Zeiss LSM 900 confocal microscope with 40× oil objective. Confocal Z-stacks were acquired in 2 μm, and analyzed using a free and open source FIJI (https://fiji.sc).

### *In vivo* Tric-Luciferase assays

5-7 days flies (*MB299B-split-GALl4>Tric-LUC* and *VT064246-GAL4>Tric-LUC*) were collected and pre-fed for 2 days on food containing 5% sucrose, 2% agar and 20 mM D-luciferin potassium salt (Cat# 115144-35-9, Gold Biotechnology). Flies were then transferred to starvation tubes containing 2% agar food supplemented with D-luciferin potassium salt and starved for 22 hours Subsequently, flies underwent either 2 min sucrose-odor pairing training or remained untrained., They were then placed into white 96-well Microfluor 2 plates (Costar) with starvation medium. Luciferase activity from individual flies was recorded using a Microbeta 2 microplate counter (PerkinElmer), with luminescence signals collected continuously for 3 hours.

### *In vitro* calcium imaging

Functional imaging experiments of neurons were performed on both male and female flies aged 5-7 days. Adult hemolymph-like saline (AHL) composed with (Concentration, mM): NaCl 108; KCl 5; NaH_2_PO4 1; MgCl_2_.6H_2_O 8.2; CaCl_2_.2H_2_O 2; NaHCO_3_ 4; trehalose 5; sucrose 10; and HEPES 5. Adenosine 5′-triphosphate magnesium salt (ATP, Cat# A100885, Aladdin) was dissolved in AHL immediately prior to the experiment to the final concentration of 2.5 mM. Tetrodotoxin (TTX, Cat# 554412, Sigma-Aldrich) and dopamine (DA, Cat# 73483, Sigma-Aldrich) were made freshly to final concentrations of 1 μM and 1 mM, respectively.

Flies of genotypes *w^1118^;R58E02-LexA/+;VT064246-GAL4/LexAop-P2X_2_, UAS-GCaMP6f, w^1118^;UAS-dcr2/+;VT046004-GAL4/UAS-EPAC, w^1118^;UAS-dcr2 /UAS-Dop1R1^RNAi^;VT046004-GAL4/UAS-EPAC,w^1118^;UAS-dcr2/UAS-Dop1R2^RNAi^;VT046004-GAL4/UAS-EPAC*and *w^1118^;UAS-P2X_2_,LexAop-GCaMP6f/R58E02-p65.AD;VT064246-LexA/R37E10-GAL4.DBD* were employed. Flies were anesthetized on ice, and brains were dissected in AHL (PH = 7.5) at room temperature. Dissected brains were then pinned to a layer of Sylgard silicone (Dow Corning, Midland, MI) in a perfusion chamber containing AHL. Drugs were applied though a gravity-fed ValveLink perfusion system that allowed a switch from one channel to the other (Automate Scientific, Berkeley, CA).

For functional connectivity experiments, after 30 s of baseline recording with perfusion of AHL, ATP was delivered for 90 s. As a control condition, AHL was perfused for 30 s and switched to a second channel of AHL for another 90 s. Brains expressing the GCaMP6f were imaged using an Mshot MF43-N fluorescence microscope (Mshot, Guangdong, China) under an Olympus × 40 (0.80W, LUMPlanFL N) water-immersion objective, and recordings were captured using a camera of Hamamatsu ORCA-Flash4.0 LT+. Images were captured using hCimage Live software. Regions of interests (ROIs) were selected in the MB of PAM/PAM-α1 and cell bodies of DPM neurons. ROIs were analyzed using FIJI. The percent change in fluorescence over time was calculated using △F / F = (F_n_-F_0_) / F_0_ ×100% as we previously described (Liu et al., 2019). F_n_ is the fluorescence at time point n, and F_0_ is the fluorescence at time 0. F is calculated as the fluorescence difference between the ROI and the background. Averaged fluorescence change values were quantified for periods of 0-30 s and 31-120 s, respectively. The sample size for DPM neural activity analyses was defined by the number of recorded cell bodies.

For EPAC experiments, brains were exposed to fluorescent light for 5 min to reduce the photobleaching rates between the CFP and YPF. Also, TTX was applied for 5 min prior to the recording to block voltage-dependent sodium channels and throughout the entire recording period. After 30 s of baseline recording, DA was delivered for 120 s and switched back to TTX again for another 30 s. Brains expressing the FRET sensor EPACcamps1 were imaged using an Olympus BX51WI fluorescence microscope (Olympus, Center Valley, PA) under an Olympus × 40 (0.8W, LUMPlanFI) water-immersion objective, and recordings were captured using a charge-coupled device camera (Hamamatsu ORCAC472-80-12AG). Images were captured using μManager acquisition software (Edelstein et al., 2010). ROIs analysis was the same as previously described using custom software developed in MATLAB (Liu et al., 2019). Identical ROIs were selected from both the CFP and YFP channels, and calculated the ratio of normalized CFP/YFP to the first 20 frames of baseline recording for each time point. The maximum change values were determined for each group during drug perfusion period (31-150 s) and TTX washout period (151-240 s) for the following quantification.

### *In vivo* calcium imaging

All-trans-retinal (ATR) powder (Cat# R2500, Sigma) was dissolved in 100% alcohol to prepare a 100 mM stock solution. For ATR food used in the optogenetic experiments, 240 μl of stock solution was added to 30 ml of 5% sucrose, 1% agar medium to yield final concentration of approximately 800 μM ATR. 3-5 day-old flies (*VT064246-LexA/LexAop-GCaMP6f;R58E02-GAL4/UAS-Cschrimson*and *VT064246-LexA/LexAop-GCaMP6f;+/UAS-Cschrimson,*) were transferred to ATR food and maintained for 3 days prior to optogenetic imaging.

Flies were anesthetized on ice, and their heads were fixed in a custom-made imaging chamber using UV-curable glue. A small window was carefully dissected in the anterior head cuticle under extracellular saline solution AHL. For optogenetic activation, a 630 nm LED was focused onto the window at an intensity of 1-2 mW. Images were acquired at 1024 × 1024 pixels with a frame rate of 2 Hz. ROIs were analyzed using Fiji. Signal quantification was performed using the same method as for in vitro calcium imaging.

### Quantitative PCR

To valid the efficiency of the dopamine receptor knockdown, UAS-RNAi flies were crossed with elav-GAL4 flies. GAL4 and UAS homozygous parental lines were used as genetic controls. For quantitative PCR (qPCR), total RNA was extracted from 40 fly heads per sample using the RNeasy Mini Kit (Cat# 74104, QIAGEN), and cDNA was synthesized using the ReverTra Ace qPCR RT Kit (Cat# FSQ-201, TOYOBO). Three biological replicates were performed for each genotype, with two technical replicates per biological sample.

qPCRs were conducted using the SYBR Green Master Mix (Cat# QPK-201, TOYOBO). Raw fluorescence data were processed using dynamic tube and slope correction via QuantStudio^TM^ Real-Time PCR software. Relative mRNA expression levels were calculated using the comparative CT method, with rp49 as the endogenous reference gene. qPCR primers (0.1 mM each, 5’-3’) were used as follows: forward primer for Dop1R1: GCCGCTGTCACTTGTGTGTCAATTGTAG; reverse primer for Dop1R1: ACACCGGCAAAGGTCATCACCAGC; forward primer for Dop1R2: GGCCACCAACTCTCTCATCACCAGC; reverse primer for Dop1R2: AGATTCAGTATGGAGGCGGTGCTG; and forward primer for rp49: GCTAAGCTGTCGCACAAA; reverse primer for rp49, TCCGGTGGGCAGCATGTG.

### Experimental design and statistical analysis

All the raw data were analyzed using the Prism 9.0 software (GraphPad). For most memory experiments, datasets with normal distribution, a one-way ANOVA followed by Sidak’s multiple comparisons test was applied. For those did not have a normal distribution, nonparametric statistics, Kruskal-Wallis, followed by Dunn’s multiple comparisons test was applied. *In vitro/vivo* functional imaging experiments (Figure 2A, B, Supplemental Figure 3C-E and Figure 8B) and Tric-LUC experiments (Figure 8F), two-way ANOVA followed by Tukey’s and Sidak’s multiple comparisons test were applied. Furthermore, in the experiments to compare the activation group to the no-activation group (Figure 6F), an unpaired t test or nonparametric test was applied based on data distribution. For sleep experiments, a two-way ANOVA followed by Dunn’s multiple comparisons test was applied to determine statistical significance between the experimental groups and the control groups for each 1 h bin. All data were presented as mean ± standard error (SEM). A statistically significant difference was set to p < 0.05. Detailed statistical analysis of each experiment was summarized in Tables 1-4: Table 1 for one-way ANOVA analysis, Table 2 for unpaired t-test analysis, Table 4 for two-way ANOVA analysis for sleep, and Table 3 for the rest two-way ANOVA analysis.

## Data availability statement

All data are contained within the manuscript. Statistical analysis details, including sample size, statistical tests and p values have been provided in Tables 1-4. This work generated fly stocks, which we will distribute on request. Raw data will be distributed on request.

## Acknowledgments

This work is supported by National Natural Science Foundation of China (32371063, 82341248, 32071009 to C.L., 82171430 to F.L.), Guangdong Basic and Applied Basic Research Foundation (2024A151501150 to C.L., 2025A1515011127 to Z.M.), CAS Key Laboratory of Brain Connectome and Manipulation (2019DP173024) and National Institute of Health grant R01MH67284 (to L.C.G.), and Shenzhen Science and Technology Program (KQTD20221101093608028). We thank Kristine Paik for help with the behavioral experiments at the beginning of this study, Dr. Martha L. Reed and all lab members for their valuable discussion.

## Additional information

## Author Contributions

C. L. designed and supervised research; L.Y., L.W., T.D.W., X.S., W.Y., H.L., and C.L. performed research; L.Y., L.L., Z.L., Y.L., Z.M., F.G., L.C.G. and C.L. analyzed and interpreted data; L.Y., Z.M., Y.L., F.L., L.C.G. and C.L. wrote the manuscript and all authors reviewed the manuscript.

## Supplemental Figure Legends

**Supplemental Figure 1 related to Figure 4.**
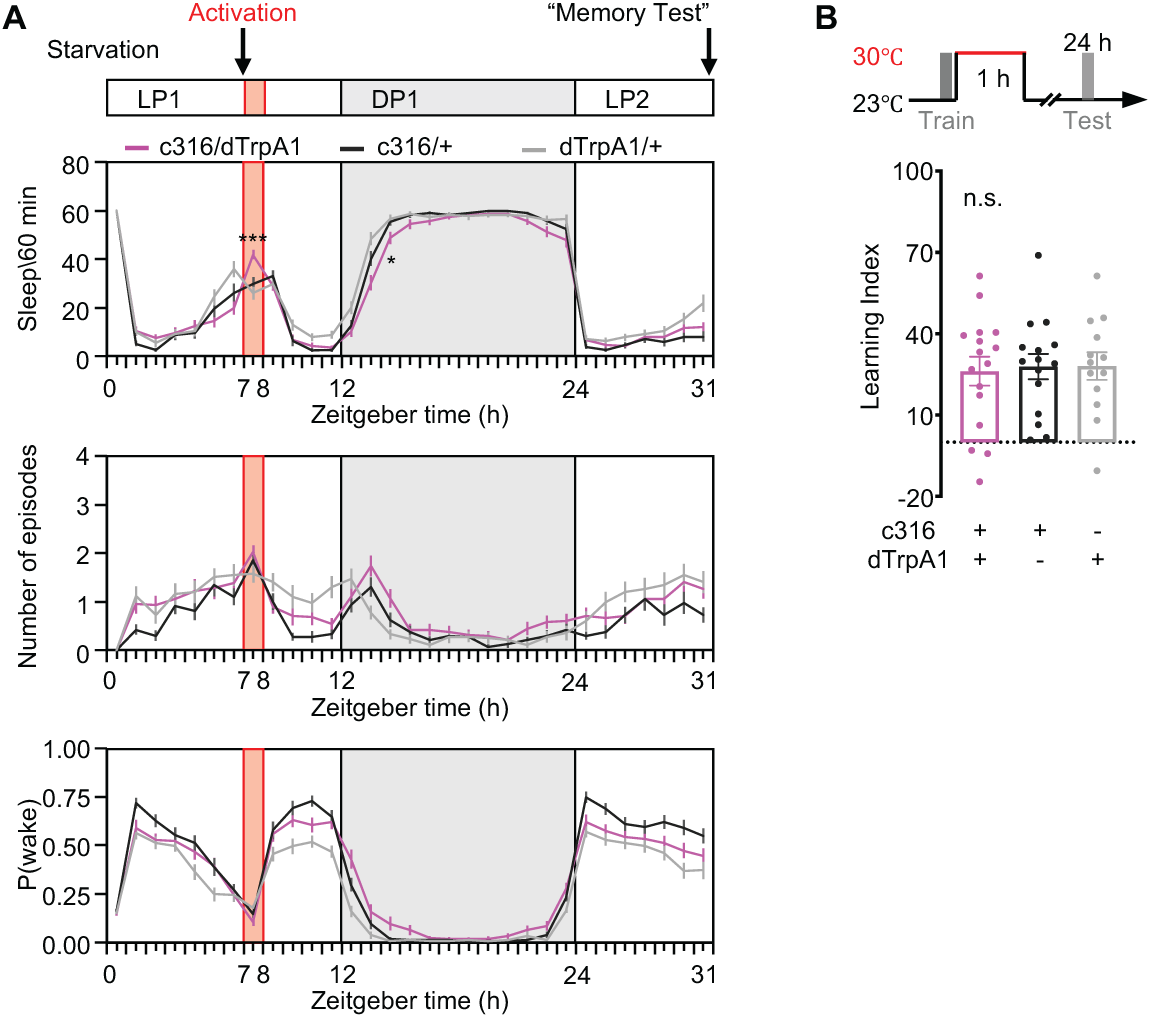
Activation of DPM neurons increases sleep but has no effect on memory consolidation. (A) Activation of DPM neurons labeled by eyeless-GAL80;MB-GAL80;c316-GAL4 under starvation increased total sleep during the activation period, with little to no effect afterwards. n = 35-42. (B) Activation of DPM neurons for 1 h immediately following training did not affect 24 h memory. n = 13-16. See also Tables 1 and 4.

**Supplemental Figure 2 related to Figure 7.**
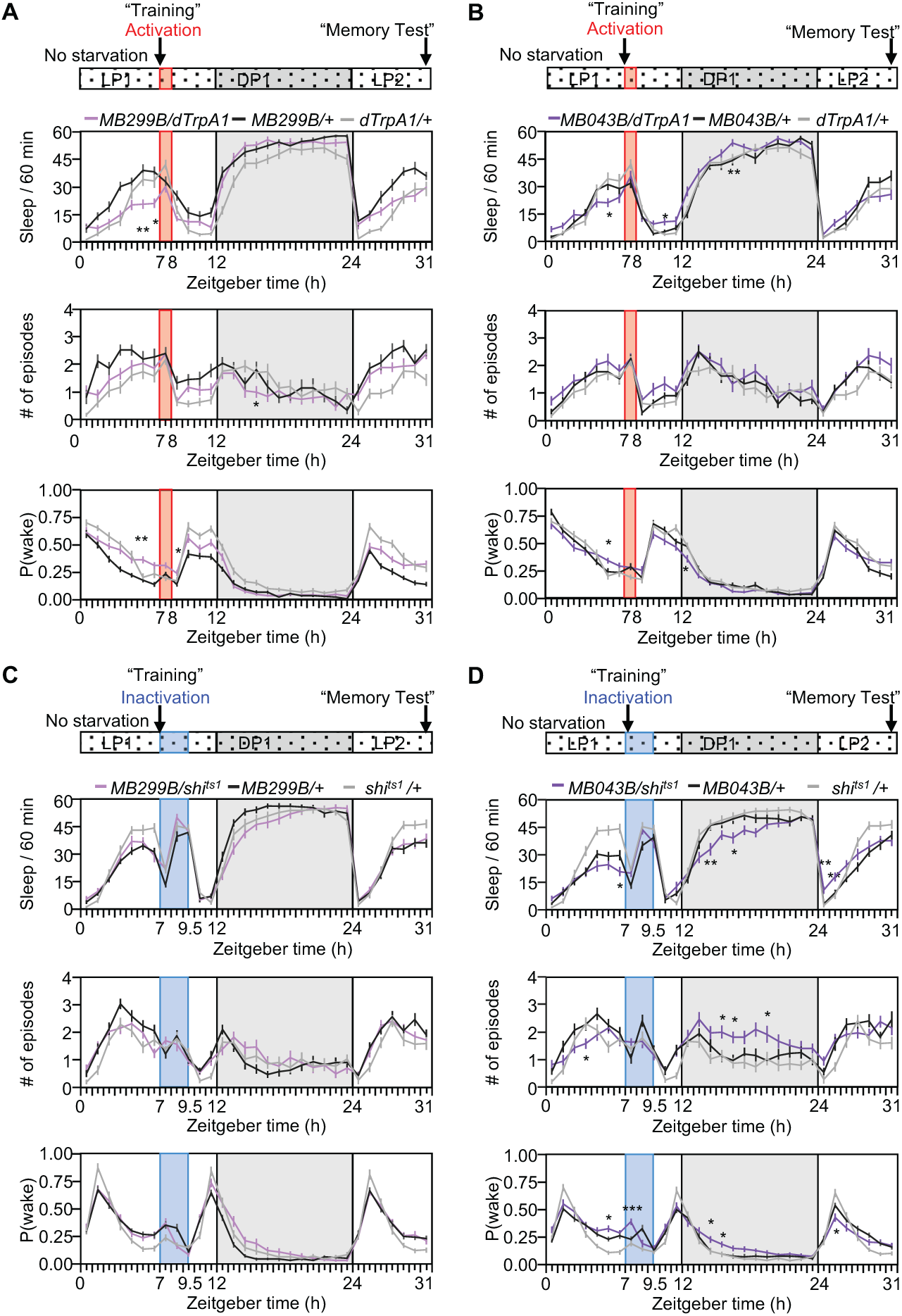
The PAM-α1 subset linking memory consolidation and sleep depends on internal starvation status. (A-B) Sleep profiles of total sleep, the number of sleep episodes, and P(wake) with 1 h activation of MB299B (A) or MB043B (B) before, during and after activation. Sleep was not affected without starvation. n = 28-31. (C) Inactivation of MB299B for 2.5 h without starvation had no effects on sleep. n = 30-37. (D) Inactivation of MB043B for 2.5 h without starvation resulted a significant reduction in total sleep, a mild increase of number of sleep episodes during the dark period, and no effect on P(wake). n = 30-39. See also Table 4.

**Supplemental Figure 3 related to Figure 10.**
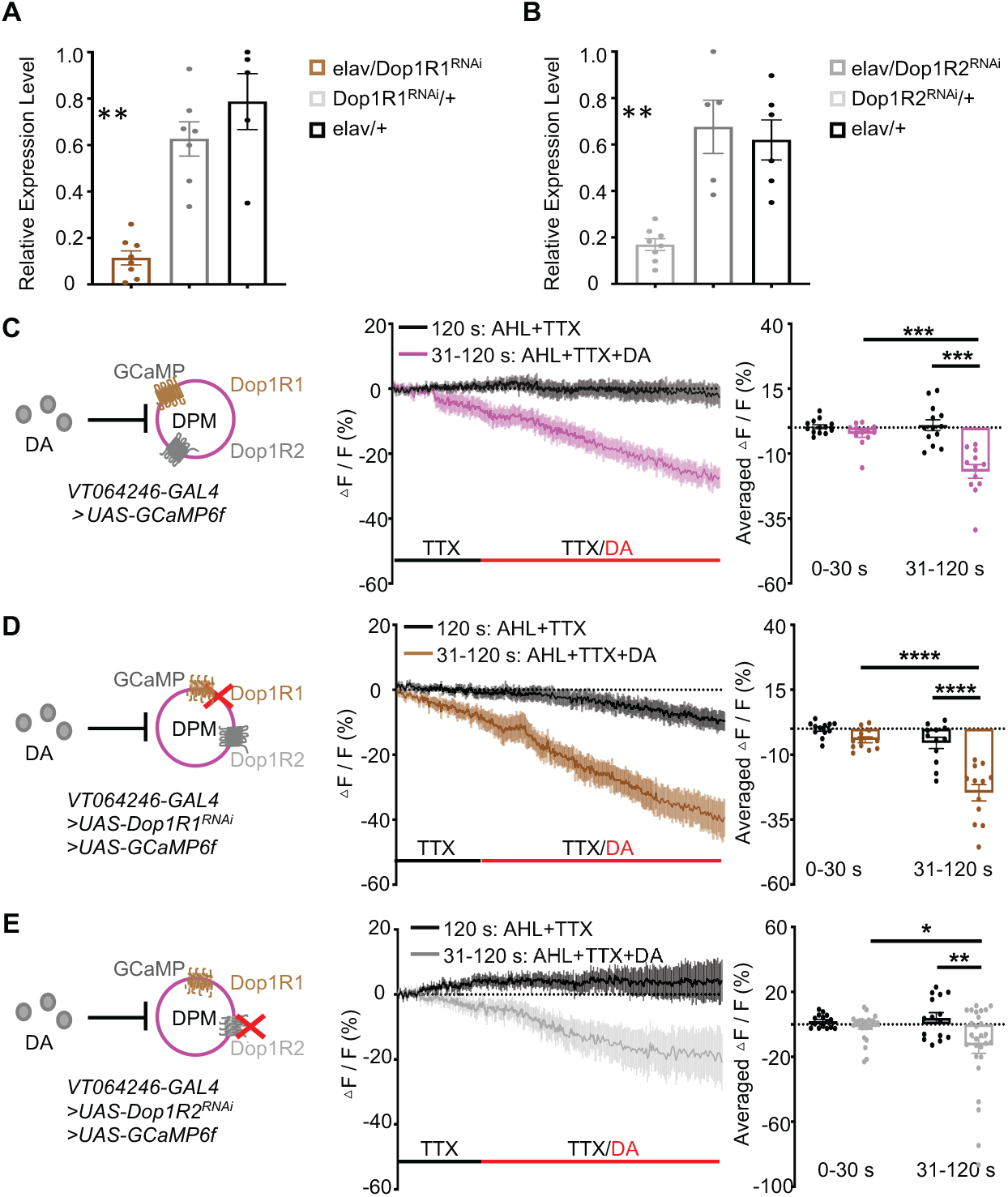
Verification of Dop1R1 and Dop1R2 knockdown efficiency and their effects on neuronal activity. (A-B) Relative mRNA levels of Dop1R1 (A) and Dop1R2 (B) in adult fly heads. Knockdown of either Dop1R1 or Dop1R2 in elav-GAL4 labeling cells resulted in a significant reduction in transcript levels compared to controls. Data are presented as mean ± SEM (n = 5-8). **p < 0.01. (C-E) Averaged GCaMP traces (upper panels) and quantification (lower panels) of DPM neurons in response to DA with the presence of 2.0 mM TTX with or without Dop1R1 or Dop1R2 knocked-down. (C, n = 12-13; D, n = 12 for all groups; E, n = 15-27). See also Tables 1 and 3.

