## Supplemental Table 1 for "Brief disruption of activity in a subset of dopaminergic neurons during consolidation impairs long-term memory by fragmenting sleep"

**Table 1. One way ANOVA for memory and neural activity tests (Related to Figs. 3, 6, 7, 9-11, S1, S3)**

| | ANOVA | DFn, DFd | F | p | post-hoc test | Comparisions | n1 | n2 | Mean Diff. | Adjusted p Value | Summary |
|---|---|---|---|---|---|---|---|---|---|---|---|
| Fig. 3A | Kruskal-Wallis | 3, 27 | 16.75 | 0.0002 | Dunn's | R58E02/shi$^{ts1}$ vs R58E02/+ | 9 | 8 | -51.69 | 0.0033 | ** |
| | | | | | | R58E02/shi$^{ts1}$ vs shi$^{ts1}$/+ | 9 | 10 | -56.39 | 0.0003 | *** |
| Fig. 3B | One way ANOVA | 2, 40 | 12.04 | <0.0001 | Sidak's | c316/shi$^{ts1}$ vs c316/+ | 14 | 14 | -25.95 | 0.0003 | *** |
| | | | | | | c316/shi$^{ts1}$ vs shi$^{ts1}$/+ | 14 | 15 | -26.15 | 0.0002 | *** |
| Fig. 3C | One way ANOVA | 2, 90 | 7.534 | 0.0009 | Sidak's | VT064246/shi$^{ts1}$ vs VT064246/+ | 28 | 35 | -10.81 | 0.0158 | * |
| | | | | | | VT064246/shi$^{ts1}$ vs shi$^{ts1}$/+ | 28 | 30 | -15.67 | 0.0005 | *** |
| Fig. 3D | One way ANOVA | 2, 20 | 0.384 | 0.6861 | Sidak's | c316/shi$^{ts1}$ vs c316/+ | 9 | 6 | -6.252 | 0.7470 | ns |
| | | | | | | c316/shi$^{ts1}$ vs shi$^{ts1}$/+ | 9 | 8 | 0.8992 | 0.9928 | ns |
| Fig. 3E | One way ANOVA | 2, 47 | 12.90 | <0.0001 | Sidak's | R58E02/dTrpA1 vs R58E02/+ | 16 | 18 | -31.73 | <0.0001 | **** |
| | | | | | | R58E02/dTrpA1 vs dTrpA1/+ | 16 | 16 | -23.71 | 0.0016 | ** |
| Fig. 3F | One way ANOVA | 2, 44 | 3.187 | 0.0510 | Sidak's | R58E02/dTrpA1 vs R58E02/+ | 15 | 18 | -13.21 | 0.0331 | * |
| | | | | | | R58E02/dTrpA1 vs dTrpA1/+ | 15 | 14 | -9.268 | 0.2037 | ns |
| Fig. 3G | One way ANOVA | 2, 44 | 3.827 | 0.0293 | Sidak's | R58E02/shi$^{ts1}$ vs R58E02/+ | 18 | 15 | -4.409 | 0.6971 | ns |
| | | | | | | R58E02/shi$^{ts1}$ vs shi$^{ts1}$/+ | 18 | 14 | -16.03 | 0.0185 | ns |
| Fig. 6B | One way ANOVA | 2, 46 | 4.201 | 0.0211 | Sidak's | R15A04/dTrpA1 vs R15A04/+ | 16 | 17 | -18.97 | 0.0386 | * |
| | | | | | | R15A04/dTrpA1 vs dTrpA1/+ | 16 | 16 | -19 | 0.0417 | * |
| Fig. 6D | Kruskal-Wallis | 3, 54 | 2.163 | 0.3391 | Dunn's | R48B04/dTrpA1 vs R48B04/+ | 21 | 17 | -0.444 | >0.9999 | ns |
| | | | | | | R48B04/dTrpA1 vs dTrpA1/+ | 21 | 16 | -7.082 | 0.3498 | ns |
| Fig. 7A | One way ANOVA | 2, 44 | 5.597 | 0.0068 | Sidak's | MB299B/dTrpA1 vs MB299B/+ | 14 | 16 | -25.14 | 0.0073 | ** |
| | | | | | | MB299B/dTrpA1 vs dTrpA1/+ | 14 | 17 | -22.45 | 0.0159 | * |
| Fig. 7B | One way ANOVA | 3, 54 | 11.72 | 0.0029 | Sidak's | MB043B/dTrpA1 vs MB043B/+ | 18 | 20 | -13.69 | 0.0148 | * |
| | | | | | | MB043B/dTrpA1 vs dTrpA1/+ | 18 | 16 | -17.2 | 0.0029 | ** |
| Fig. 7C | One way ANOVA | 2, 36 | 9.9375 | 0.0005 | Sidak's | MB299Bshi$^{ts1}$ vs MB299B/+ | 14 | 11 | -24.98 | 0.0034 | ** |
| | | | | | | MB299B/shi$^{ts1}$ vs shi$^{ts1}$/+ | 14 | 14 | -27.41 | 0.0007 | *** |
| Fig. 7D | Kruskal-Wallis | 3, 65 | 1.966 | 0.3742 | Dunn's | MB043B/shi$^{ts1}$ vs MB043B/+ | 24 | 16 | -5.969 | 0.6560 | ns |

| | | | | | | | | | | | |
|---|---|---|---|---|---|---|---|---|---|---|---|
| | | | | | | MB043B/shi$^{ts1}$ vs shi$^{ts1}$/+ | 24 | 25 | -7.23 | 0.3617 | ns |
| Fig. 7E | One way ANOVA | 2, 37 | 4.849 | 0.0135 | Sidak's | MB299B/dTrpA1 vs MB299B/+ | 12 | 11 | -18.26 | 0.0378 | * |
| | | | | | | MB299B/dTrpA1 vs dTrpA1/+ | 12 | 17 | -19.7 | 0.0116 | * |
| Fig. 7F | One way ANOVA | 2, 56 | 0.5049 | 0.6063 | Sidak's | MB299B/dTrpA1 vs MB299B/+ | 18 | 18 | -2.053 | 0.8939 | ns |
| | | | | | | MB299B/dTrpA1 vs dTrpA1/+ | 18 | 23 | -4.574 | 0.5417 | ns |
| Fig. 7G | Kruskal-Wallis | 3, 28 | 2.770 | 0.2503 | Dunn's | MB299Bshi$^{ts1}$ vs MB299B/+ | 9 | 9 | -2.0000 | >0.9999 | ns |
| | | | | | | MB299B/shi$^{ts1}$ vs shi$^{ts1}$/+ | 9 | 10 | -6.133 | 0.2092 | ns |
| Fig. 9F | One way ANOVA | 2, 45 | 2.631 | 0.0831 | Sidak's | MB299B/dTrpA1 vs MB299B/+ | 17 | 15 | -8.711 | 0.1548 | ns |
| | | | | | | MB299B/dTrpA1 vs dTrpA1/+ | 17 | 16 | -10.14 | 0.0783 | ns |
| Fig. 10D | One way ANOVA | 2, 48 | 0.1022 | 0.9030 | Sidak's | VT064246/Dop1R1$^{RNAi}$ vs Dop1R1$^{RNAi}$/+ | 19 | 14 | 1.385 | 0.9814 | ns |
| | | | | | | VT064246/Dop1R1$^{RNAi}$ vs VT064246/+ | 19 | 18 | 3.379 | 0.8805 | ns |
| Fig. 10E | One way ANOVA | 3, 64 | 14.31 | 0.0008 | Dunn's | VT064246/Dop1R1$^{RNAi}$ vs Dop1R1$^{RNAi}$/+ | 22 | 22 | -20.14 | 0.0007 | *** |
| | | | | | | VT064246/Dop1R1$^{RNAi}$ vs VT064246/+ | 22 | 20 | -16.10 | 0.0102 | * |
| Fig. 10G | One way ANOVA | 2, 53 | 0.5260 | 0.5940 | Sidak's | VT064246/Dop1R2$^{RNAi}$ vs Dop1R2$^{RNAi}$/+ | 19 | 17 | -6.487 | 0.5529 | ns |
| | | | | | | VT064246/Dop1R2$^{RNAi}$ vs VT064246/+ | 19 | 20 | -4.730 | 0.7080 | ns |
| Fig. 10H | One way ANOVA | 3, 43 | 8.302 | 0.0009 | Sidak's | VT064246/Dop1R2$^{RNAi}$ vs Dop1R2$^{RNAi}$/+ | 15 | 15 | -18.04 | 0.0240 | * |
| | | | | | | VT064246/Dop1R2$^{RNAi}$ vs VT064246/+ | 15 | 16 | -27.24 | 0.0005 | *** |
| Fig. 11E | One way ANOVA | 2, 53 | 9.050 | 0.0004 | Dunn's | MB310C/dTrpA1 vs MB310C/+ | 17 | 20 | -23.64 | 0.0002 | **** |
| | | | | | | MB310C/dTrpA1 vs dTrpA1/+ | 17 | 19 | -16.37 | 0.0113 | * |
| Fig. 11F | One way ANOVA | 2, 49 | 11.31 | <0.0001 | Sidak's | MB310C/shi$^{ts1}$ vs MB310C/+ | 17 | 17 | -17.67 | 0.0045 | ** |
| | | | | | | MB310C/shi$^{ts1}$ vs shi$^{ts1}$/+ | 17 | 18 | -25.12 | <0.0001 | **** |
| Fig. S1B | One way ANOVA | 2, 41 | 0.04409 | 0.9569 | Sidak's | c316/dTrpA1 vs c316/+ | 16 | 15 | -1.700 | 0.9633 | ns |
| | | | | | | c316/dTrpA1 vs dTrpA1/+ | 16 | 13 | -1.906 | 0.9574 | ns |
| Fig. S3A | One way ANOVA | 3, 21 | 15.37 | <0.0001 | Dunn's | elav/Dop1R1$^{RNAi}$ vs Dop1R1$^{RNAi}$/+ | 9 | 7 | -9.286 | 0.0060 | ** |
| | | | | | | elav/Dop1R1$^{RNAi}$ vs elav/+ | 9 | 5 | -12.20 | 0.0008 | *** |
| Fig. S3B | One way ANOVA | 3, 19 | 13.39 | <0.0001 | Dunn's | elav/Dop1R2$^{RNAi}$ vs Dop1R2$^{RNAi}$/+ | 8 | 5 | -10.30 | 0.0026 | ** |
| | | | | | | elav/Dop1R2$^{RNAi}$ vs elav/+ | 8 | 6 | -8.833 | 0.0073 | ** |
