## Supplemental Table 2 for "Brief disruption of activity in a subset of dopaminergic neurons during consolidation impairs long-term memory by fragmenting sleep"

**Table 2. Unpaired t test for 24 h memory (Related to Figs. 6 and 9)**

| | Comparisions | Test | df | t | p |
|---|---|---|---|---|---|
| Fig. 6F | MB299B/dTrpA1: Activation vs No-activation | Unpaired t test | 19 | 2.9940 | 0.0075 |
| | MB043B/dTrpA1: Activation vs No-activation | Unpaired t test | 9 | 4.7900 | 0.0010 |
| | MB213B/dTrpA1: Activation vs No-activation | Unpaired t test | 11 | 0.7516 | 0.4680 |
| | MB025B/dTrpA1: Activation vs No-activation | Unpaired t test | 14 | 0.3025 | 0.7667 |
| | MB032B/dTrpA1: Activation vs No-activation | Unpaired t test | 14 | 0.3011 | 0.7677 |
| | MB315C/dTrpA1: Activation vs No-activation | Unpaired t test | 22 | 0.7166 | 0.4812 |
| Fig. 9G | MB299B/dTrpA1: THIP- vs THIP+ | Unpaired t test | 32 | 2.8970 | 0.0067 |
