## Supplemental Table 3 for "Brief disruption of activity in a subset of dopaminergic neurons during consolidation impairs long-term memory by fragmenting sleep"

**Table 3. Two-way ANOVA for functional imaging and memory tests (Related to Figs. 2, 8, 10 and Fig. S3)**

| | DFn, DFd | Group | | Time | | Interaction | | Post-hoc Test | Comparisions | | Adjusted p Value | Summar y |
|---|---|---|---|---|---|---|---|---|---|---|---|---|
| | | F | p | F | p | F | p | | | | | |
| Fig. 2A | 2, 48 | 16.20 | ＜ 0.0001 | 48.65 | ＜ 0.0001 | 20.27 | ＜0.0001 | Tukey's | 0-30 s | R58E02 > P2X2;VT064246 > GCaMP6f: AHL vs AHL+ATP | 0.7409 | ns |
| | | | | | | | | | | AHL vs AHL+ATP: R58E02＞P2X2;VT064246＞GCaMP6f vs +/P2X2;VT064246＞GCaMP6f | 0.9786 | ns |
| | | | | | | | | | | AHL+ATP: R58E02＞P2X2;VT064246＞GCaMP6f vs +/P2X2;VT064246＞GCaMP6f | 0.6585 | ns |
| | | | | | | | | | 31-120 s | R58E02＞P2X2;VT064246＞GCaMP6f: AHL vs AHL+ATP | ＜0.0001 | **** |
| | | | | | | | | | | AHL vs AHL+ATP: R58E02＞P2X2;VT064246＞GCaMP6f vs +/P2X2;VT064246＞GCaMP6f | 0.5258 | ns |
| | | | | | | | | | | AHL+ATP: R58E02＞P2X2;VT064246＞GCaMP6f vs +/P2X2;VT064246＞GCaMP6f | ＜0.0001 | **** |
| | 2, 96 | 21.60 | ＜ 0.0001 | 32.44 | ＜ 0.0001 | 13.52 | ＜0.0001 | Sidak's | AHL | 0-30 s vs 31-120 s | 0.1909 | ns |
| | | | | | | | | | AHL+ATP (P2X2;VT064246＞GCaMP6f) | 0-30 s vs 31-120 s | 0.8484 | ns |

| | | | | | | | | | | | | |
|---|---|---|---|---|---|---|---|---|---|---|---|---|
| | | | | | | | | | AHL+ATP (R58E02＞P2X2;VT064246＞GCaMP6f) | 0-30 s vs 31-120 s | ＜0.0001 | **** |
| Fig. 2B | 2, 84 | 13.50 | ＜ 0.0001 | 24.54 | ＜ 0.0001 | 7.868 | 0.0007 | Sidak's | 0-30s | VT064246＞GCaMP6f; R58E02＞CsChrimson vs VT064246＞GCaMP6f;+/CsChrimson | 0.9999 | ns |
| | | | | | | | | | 31-60 s | | 0.0076 | **** |
| | | | | | | | | | 61-120 s | | ＜0.0001 | **** |
| | 1, 84 | 24.54 | ＜ 0.0001 | 13.50 | ＜ 0.0001 | 7.868 | 0.0007 | Tukey's | VT064246＞GCaMP6f;R58E02＞CsChrimson | 0-30 s vs 31-60 s | 0.0012 | ** |
| | | | | | | | | | | 0-30 s vs 61-120 s | ＜0.0001 | **** |
| | | | | | | | | | | 31-60 s vs 61-120 s | 0.0094 | ** |
| Fig. 8B | 2, 54 | 15.46 | 0.0001 | 29.23 | ＜ 0.0001 | 13.87 | ＜0.0001 | Tukey's | 0-30 s | MB299B＞P2X2;VT064246＞GCaMP6f: AHL vs AHL+ATP | 0.2995 | ns |
| | | | | | | | | | | AHL vs AHL+ATP: MB299B＞P2X2;VT064246＞GCaMP6f vs +/P2X2;VT064246＞GCaMP6f | 0.9880 | ns |
| | | | | | | | | | | AHL+ATP: MB299B＞P2X2;VT064246＞GCaMP6f vs +/P2X2;VT064246＞GCaMP6f | 0.5742 | ns |
| | | | | | | | | | 31-120 s | MB299B＞P2X2;VT064246＞GCaMP6f: AHL vs AHL+ATP | ＜0.0001 | **** |
| | | | | | | | | | | AHL vs AHL+ATP: MB299B＞P2X2;VT064246＞GCaMP6f vs +/P2X2;VT064246＞GCaMP6f | 0.2327 | ns |

| | | | | | | | | | | | | |
|---|---|---|---|---|---|---|---|---|---|---|---|---|
| | | | | | | | | | | AHL+ATP: R58E02＞P2X2;VT064246＞GCaMP6f vs +/P2X2;VT064246＞GCaMP6f | 0.0002 | *** |
| | 2, 108 | 20.82 | ＜ 0.0001 | 19.10 | ＜ 0.0001 | 9.065 | 0.0002 | Sidak's | AHL | 0-30 s vs 31-120 s | 0.9694 | ns |
| | | | | | | | | | AHL+ATP (+/P2X2;VT064246＞GCaMP6f) | 0-30 s vs 31-120 s | 0.3381 | ns |
| | | | | | | | | | AHL+ATP (R58E02＞P2X2;VT064246＞GCaMP6f) | 0-30 s vs 31-120 s | ＜0.0001 | **** |
| Fig. 8D | 20, 7238 | 2.731 | ＜ 0.0001 | 5.461 | 0.0195 | 1.376 | 0.1216 | Sidak's | Post-training time-1 | | 0.7542 | ns |
| | | | | | | | | | Post-training time-2 | | >0.9999 | ns |
| | | | | | | | | | Post-training time-3 | | >0.9999 | ns |
| | | | | | | | | | Post-training time-4 | | >0.9999 | ns |
| | | | | | | | | | Post-training time-5 | | >0.9999 | ns |
| | | | | | | | | | Post-training time-6 | | >0.9999 | ns |
| | | | | | | | | | Post-training time-7 | Un-trained: PAM-α1 vs DPM | 0.9995 | ns |
| | | | | | | | | | Post-training time-8 | | 0.6254 | ns |
| | | | | | | | | | Post-training time-9 | | >0.9999 | ns |
| | | | | | | | | | Post-training time-10 | | >0.9999 | ns |
| | | | | | | | | | Post-training time-11 | | >0.9999 | ns |
| | | | | | | | | | Post-training time-12 | | 0.9805 | ns |
| | | | | | | | | | Post-training time-13 | | >0.9999 | ns |

| | | | | | | | | | | | | |
|---|---|---|---|---|---|---|---|---|---|---|---|---|
| | | | | | | | | | Post-training time-14 | | >0.9999 | ns |
| | | | | | | | | | Post-training time-15 | | >0.9999 | ns |
| | | | | | | | | | Post-training time-16 | | >0.9999 | ns |
| | | | | | | | | | Post-training time-17 | | >0.9999 | ns |
| | | | | | | | | | Post-training time-18 | | 0.0012 | ** |
| | | | | | | | | | Post-training time-19 | | 0.9974 | ns |
| | | | | | | | | | Post-training time-20 | | 0.9875 | ns |
| | | | | | | | | | Post-training time-21 | | >0.9999 | ns |
| Fig. 8E | 20, 5561 | 2.230 | 0.0013 | 10.87 | 0.0010 | 1.456 | 0.0859 | Sidak's | Post-training time-1 | | 0.9275 | ns |
| | | | | | | | | | Post-training time-2 | | >0.9999 | ns |
| | | | | | | | | | Post-training time-3 | | 0.5864 | ns |
| | | | | | | | | | Post-training time-4 | | 0.9986 | ns |
| | | | | | | | | | Post-training time-5 | | >0.9999 | ns |
| | | | | | | | | | Post-training time-6 | | >0.9999 | ns |
| | | | | | | | | | Post-training time-7 | Trained: PAM-α1 vs DPM | >0.9999 | ns |
| | | | | | | | | | Post-training time-8 | | >0.9999 | ns |
| | | | | | | | | | Post-training time-9 | | >0.9999 | ns |
| | | | | | | | | | Post-training time-10 | | >0.9999 | ns |
| | | | | | | | | | Post-training time-11 | | >0.9999 | ns |
| | | | | | | | | | Post-training time-12 | | 0.9387 | ns |
| | | | | | | | | | Post-training time-13 | | 0.3313 | ns |
| | | | | | | | | | Post-training time-14 | | 0.8392 | ns |

| | | | | | | | | | | | | |
|---|---|---|---|---|---|---|---|---|---|---|---|---|
| | | | | | | | | | Post-training time-15 | | 0.9929 | ns |
| | | | | | | | | | Post-training time-16 | | 0.9202 | ns |
| | | | | | | | | | Post-training time-17 | | 0.3396 | ns |
| | | | | | | | | | Post-training time-18 | | 0.4471 | ns |
| | | | | | | | | | Post-training time-19 | | 0.983 | ns |
| | | | | | | | | | Post-training time-20 | | 0.9989 | ns |
| | | | | | | | | | Post-training time-21 | | >0.9999 | ns |
| Fig. 8F | 3, 1929 | 6.973 | 0.0001 | 0.6061 | 0.5456 | 5.277 | ＜0.0001 | Tukey's | post-training time 1 h | MB299B-No T vs. MB299B-T | 0.9985 | ns |
| | | | | | | | | | | MB299B-No T vs. VT064246-No T | 0.6689 | ns |
| | | | | | | | | | | MB299B-T vs. VT064246-T | 0.9167 | ns |
| | | | | | | | | | | VT064246-No T vs. VT064246-T | 0.3023 | ns |
| | | | | | | | | | post-training time 2 h | MB299B-No T vs. MB299B-T | 0.8153 | ns |
| | | | | | | | | | | MB299B-No T vs. VT064246-No T | 0.2980 | ns |
| | | | | | | | | | | MB299B-T vs. VT064246-T | 0.1046 | ns |
| | | | | | | | | | | VT064246-No T vs. VT064246-T | 0.0055 | ** |
| | | | | | | | | | post-training time 3 h | MB299B-No T vs. MB299B-T | 0.4037 | ns |
| | | | | | | | | | | MB299B-No T vs. VT064246-No T | 0.0132 | * |
| | | | | | | | | | | MB299B-T vs. VT064246-T | 0.0002 | *** |
| | | | | | | | | | | VT064246-No T vs. VT064246-T | ＜0.0001 | **** |
| Fig. 10B | 2, 30 | 4.785 | 0.0157 | 90.16 | ＜ 0.0001 | 4.799 | 0.0155 | Dunnett's | 0-30 s | wCS vs Dop1R1-KK | 0.9998 | ns |
| | | | | | | | | | | wCS vs Dop1R2-GD | 0.9999 | ns |
| | | | | | | | | | | Dop1R1-KK vs Dop1R2-GD | 0.9994 | ns |
| | | | | | | | | | 31-240 s | wCS vs Dop1R1-KK | 0.0209 | * |

| | | | | | | | | | | | | |
|---|---|---|---|---|---|---|---|---|---|---|---|---|
| | | | | | | | | | | wCS vs Dop1R2-GD | 0.2665 | ns |
| | | | | | | | | | | Dop1R1-KK vs Dop1R2-GD | 0.0002 | *** |
| Fig. 10C | 2, 122 | 13.99 | < 0.0001 | 2645 | < 0.0001 | 6.955 | 0.0014 | Sidak's | LP (Total sleep) | VT064246/Dop1R1$^{RNAi}$ vs Dop1R1$^{RNAi}$/+ | <0.0001 | **** |
| | | | | | | | | | | VT064246/Dop1R1$^{RNAi}$ vs VT064246/+ | <0.0001 | **** |
| | | | | | | | | | DP (Total sleep) | VT064246/Dop1R1$^{RNAi}$ vs Dop1R1$^{RNAi}$/+ | 0.0321 | * |
| | | | | | | | | | | VT064246/Dop1R1$^{RNAi}$ vs VT064246/+ | 0.3955 | ns |
| | 2, 122 | 8.348 | 0.0004 | 66.19 | < 0.0001 | 7.535 | 0.0008 | Dunnett's | LP (# of episodes) | VT064246/Dop1R1$^{RNAi}$ vs Dop1R1$^{RNAi}$/+ | 0.5045 | ns |
| | | | | | | | | | | VT064246/Dop1R1$^{RNAi}$ vs VT064246/+ | <0.0001 | **** |
| | | | | | | | | | DP (# of episodes) | VT064246/Dop1R1$^{RNAi}$ vs Dop1R1$^{RNAi}$/+ | 0.0060 | ** |
| | | | | | | | | | | VT064246/Dop1R1$^{RNAi}$ vs VT064246/+ | 0.1710 | ns |
| | 2, 122 | 23.59 | < 0.0001 | 1290 | < 0.0001 | 18.66 | <0.0001 | Dunnett's | LP (Pwake) | VT064246/Dop1R1$^{RNAi}$ vs Dop1R1$^{RNAi}$/+ | <0.0001 | **** |
| | | | | | | | | | | VT064246/Dop1R1$^{RNAi}$ vs VT064246/+ | <0.0001 | **** |
| | | | | | | | | | DP (Pwake) | VT064246/Dop1R1$^{RNAi}$ vs Dop1R1$^{RNAi}$/+ | 0.3129 | ns |
| | | | | | | | | | | VT064246/Dop1R1$^{RNAi}$ vs VT064246/+ | 0.9947 | ns |

| | | | | | | | | | | | | |
|---|---|---|---|---|---|---|---|---|---|---|---|---|
| Fig. 10F | 2, 91 | 0.5364 | 0.5867 | 2159 | <0.0001 | 0.7245 | 0.4922 | Sidak's | LP (Total sleep) | VT064246/Dop1R2$^{RNAi}$ vs Dop1R2$^{RNAi}$/+ | 0.9935 | ns |
| | | | | | | | | | | VT064246/Dop1R2$^{RNAi}$ vs VT064246/+ | 0.9733 | ns |
| | | | | | | | | | DP (Total sleep) | VT064246/Dop1R2$^{RNAi}$ vs Dop1R2$^{RNAi}$/+ | 0.5317 | ns |
| | | | | | | | | | | VT064246/Dop1R2$^{RNAi}$ vs VT064246/+ | 0.9971 | ns |
| | 2, 91 | 4.098 | 0.0198 | 82.19 | <0.0001 | 0.9814 | 0.3787 | Sidak's | LP (# of episodes) | VT064246/Dop1R2$^{RNAi}$ vs Dop1R2$^{RNAi}$/+ | 0.9196 | ns |
| | | | | | | | | | | VT064246/Dop1R2$^{RNAi}$ vs VT064246/+ | >0.9999 | ns |
| | | | | | | | | | DP (# of episodes) | VT064246/Dop1R2$^{RNAi}$ vs Dop1R2$^{RNAi}$/+ | 0.0193 | * |
| | | | | | | | | | | VT064246/Dop1R2$^{RNAi}$ vs VT064246/+ | 0.3626 | ns |
| | 2, 91 | 3.425 | 0.0368 | 849.6 | <0.0001 | 3.982 | 0.0220 | Dunnett's | LP (Pwake) | VT064246/Dop1R2$^{RNAi}$ vs Dop1R2$^{RNAi}$/+ | 0.9928 | ns |
| | | | | | | | | | | VT064246/Dop1R2$^{RNAi}$ vs VT064246/+ | 0.0024 | ** |
| | | | | | | | | | DP (Pwake) | VT064246/Dop1R2$^{RNAi}$ vs Dop1R2$^{RNAi}$/+ | 0.6227 | ns |
| | | | | | | | | | | VT064246/Dop1R2$^{RNAi}$ vs VT064246/+ | 0.7357 | ns |
| Fig. S3C | 1, 23 | 21,77 | 0.0001 | 24.94 | <0.0001 | 29.45 | <0.0001 | Sidak's | 0-30 s | AHL+TTX vs AHL+TTX+DA | 0.5129 | ns |

| | | | | | | | | | | | | |
|---|---|---|---|---|---|---|---|---|---|---|---|---|
| | | | | | | | | | 31-120 s | AHL+TTX vs AHL+TTX+DA | ＜0.0001 | **** |
| | 1, 24 | 13.02 | 0.0014 | 33.45 | ＜ 0.0001 | 17.92 | 0.0003 | Sidak's | AHL+TTX | 0-30 s vs 31-120 s | 0.9655 | ns |
| | | | | | | | | | AHL+TTX+DA | 0-30 s vs 31-120 s | ＜0.0001 | **** |
| Fig. S3D | 1, 22 | 22.04 | 0.0001 | 77.04 | ＜ 0.0001 | 26.46 | ＜0.0001 | Sidak's | 0-30 s | AHL+TTX vs AHL+TTX+DA | 0.2944 | ns |
| | | | | | | | | | 31-120 s | AHL+TTX vs AHL+TTX+DA | ＜0.0001 | **** |
| | 1, 22 | 26.74 | ＜ 0.0001 | 51.19 | ＜ 0.0001 | 11.18 | 0.0029 | Sidak's | AHL+TTX | 0-30 s vs 31-120 s | 0.1380 | ns |
| | | | | | | | | | AHL+TTX+DA | 0-30 s vs 31-120 s | ＜0.0001 | **** |
| Fig. S3E | 1, 40 | 5.345 | 0.0260 | 2.876 | ＜ 0.0001 | 5.899 | 0.0197 | Sidak's | 0-30 s | AHL+TTX vs AHL+TTX+DA | 0.7618 | ns |
| | | | | | | | | | 31-120 s | AHL+TTX vs AHL+TTX+DA | 0.0034 | ** |
| | 1, 28 | 7.551 | 0.0104 | 2.582 | 0.1142 | 2.434 | 0.1300 | Sidak's | AHL+TTX | 0-30 s vs 31-120 s | 0.8620 | ns |
| | | | | | | | | | AHL+TTX+DA | 0-30 s vs 31-120 s | 0.0308 | * |
