## Supplemental Table 4 for "Brief disruption of activity in a subset of dopaminergic neurons during consolidation impairs long-term memory by fragmenting sleep"

**Table 4. Two-way ANOVA for sleep analysis (Related to Figs. 4, 5, 7, 9, 11, S1, S2)**

| | Data | DFn, DFd | Group | | Time | | Interaction | |
|---|---|---|---|---|---|---|---|---|
| | | | F | p | F | p | F | p |
| Fig. 4A | Total sleep | 2, 119 | 5.9350 | 0.0035 | 233.4 | <0.0001 | 6.746 | <0.0001 |
| | Sleep episodes | 2, 119 | 4.0780 | 0.0194 | 10.38 | <0.0001 | 4.768 | <0.0001 |
| | P(wake) | 2, 119 | 7.2560 | 0.0011 | 187.1 | <0.0001 | 5.960 | <0.0001 |

Dunnett's multiple comparisons

| | R58E02/dTrpA1 vs R58E02/+ | | | R58E02/dTrpA1 vs dTrpA1/+ | | | Summary | | |
|---|---|---|---|---|---|---|---|---|---|
| | Total sleep | Sleep episodes | P(wake) | Total sleep | Sleep episodes | P(wake) | Total sleep | Sleep episodes | P(wake) |
| ZT0-1 | 0.9756 | 0.8063 | 0.9645 | 0.9806 | 0.9865 | 0.0277 | ns | ns | ns |
| ZT1-2 | 0.2459 | 0.1827 | 0.1497 | 0.9896 | 0.8710 | 0.5054 | ns | ns | ns |
| ZT2-3 | 0.0007 | 0.0031 | 0.0008 | 0.9049 | 0.9681 | 0.8836 | ns | ns | ns |
| ZT3-4 | 0.0004 | 0.0006 | 0.0001 | 0.9980 | 0.8952 | 0.6851 | ns | ns | ns |
| ZT4-5 | <0.0001 | 0.2933 | <0.0001 | 0.5611 | 0.3347 | 0.3473 | ns | ns | ns |
| ZT5-6 | 0.1148 | 0.0319 | 0.3845 | 0.0009 | <0.0001 | 0.0111 | ns | * | ns |
| ZT6-7 | <0.0001 | 0.4502 | <0.0001 | <0.0001 | 0.0048 | 0.0052 | **** | ns | ** |
| ZT7-8 | <0.0001 | 0.0182 | <0.0001 | <0.0001 | 0.0010 | 0.0017 | **** | * | ** |
| ZT8-9 | <0.0001 | 0.0018 | <0.0001 | <0.0001 | 0.0016 | <0.0001 | **** | ** | **** |
| ZT9-10 | <0.0001 | 0.0036 | 0.0002 | 0.0026 | 0.1240 | 0.0026 | ** | ns | ** |
| ZT10-11 | 0.0038 | 0.0090 | <0.0001 | 0.0677 | 0.0406 | 0.0133 | ns | * | * |
| ZT11-12 | 0.028 | 0.0108 | 0.0377 | 0.9890 | 0.9447 | 0.5754 | ns | ns | ns |
| ZT12-13 | 0.9747 | 0.1524 | 0.7279 | 0.9914 | 0.0032 | 0.7997 | ns | ns | ns |
| ZT13-14 | 0.9163 | 0.0007 | 0.7183 | 0.8430 | 0.0080 | 0.9813 | ns | ** | ns |
| ZT14-15 | 0.6078 | <0.0001 | 0.8074 | 0.0909 | 0.0100 | 0.2094 | ns | * | ns |
| ZT15-16 | 0.3992 | <0.0001 | 0.2807 | 0.0159 | 0.1387 | 0.0447 | ns | ns | ns |
| ZT16-17 | 0.7681 | <0.0001 | 0.9273 | 0.0006 | 0.0019 | 0.0011 | ns | ** | ns |
| ZT17-18 | 0.2873 | 0.0029 | 0.2764 | 0.0279 | 0.0041 | 0.0005 | ns | ** | ns |
| ZT18-19 | >0.9999 | <0.0001 | 0.1640 | 0.5784 | 0.0019 | 0.1675 | ns | ** | ns |
| ZT19-20 | 0.9898 | 0.0060 | 0.8583 | 0.4328 | 0.0124 | 0.2425 | ns | * | ns |
| ZT20-21 | 0.8351 | 0.0123 | 0.9942 | 0.8891 | 0.1945 | 0.8824 | ns | ns | ns |
| ZT21-22 | 0.4996 | 0.0129 | 0.9399 | 0.7758 | 0.2699 | 0.7315 | ns | ns | ns |
| ZT22-23 | 0.7468 | 0.0010 | 0.9476 | 0.3676 | 0.1273 | 0.2538 | ns | ns | ns |
| ZT23-24 | 0.5168 | 0.0728 | 0.2801 | 0.3308 | 0.0447 | 0.9372 | ns | ns | ns |
| ZT24-25 | 0.3769 | 0.6626 | 0.9080 | 0.1401 | 0.4485 | 0.2559 | ns | ns | ns |
| ZT25-26 | 0.1607 | 0.1756 | 0.2289 | 0.9114 | 0.5869 | 0.8935 | ns | ns | ns |
| ZT26-27 | 0.2468 | 0.0393 | 0.0542 | 0.6703 | 0.6408 | 0.8712 | ns | ns | ns |
| ZT27-28 | 0.6703 | 0.1063 | 0.0501 | 0.0053 | 0.1016 | 0.6792 | ns | ns | ns |
| ZT28-29 | 0.1917 | 0.7103 | 0.0849 | 0.0345 | 0.7491 | 0.0855 | ns | ns | ns |

| | Data | DFn, DFd | Group | | Time | | Interaction | |
|---|---|---|---|---|---|---|---|---|
| | | | F | p | F | p | F | p |
| Fig. 4B | Total sleep | 2, 104 | 9.135 | 0.0002 | 228.7 | <0.0001 | 5.274 | <0.0001 |
| | Sleep episodes | 2, 104 | 11.32 | <0.0001 | 21.39 | <0.0001 | 4.321 | <0.0001 |
| | P(wake) | 2, 104 | 8.225 | 0.0005 | 178.2 | <0.0001 | 5.579 | <0.0001 |

Dunnett's multiple comparisons

| | R58E024/$shi^{ts1}$ vs R58E02/+ | | | R58E02/$shi^{ts1}$ vs $shi^{ts1}$/+ | | | Summary | | |
|---|---|---|---|---|---|---|---|---|---|
| | Total sleep | Sleep episodes | P(wake) | Total sleep | Sleep episodes | P(wake) | Total sleep | Sleep episodes | P(wake) |
| ZT0-1 | 0.0181 | 0.0094 | - | 0.8051 | 0.9767 | - | ns | ns | - |
| ZT1-2 | <0.0001 | <0.0001 | <0.0001 | 0.3105 | 0.6616 | 0.9997 | ns | ns | ns |
| ZT2-3 | 0.0079 | 0.0022 | 0.0014 | 0.9927 | 0.6353 | 0.8757 | ns | ns | ns |
| ZT3-4 | 0.0250 | 0.0310 | 0.0027 | 0.8589 | 0.5669 | 0.9968 | ns | ns | ns |
| ZT4-5 | 0.0084 | 0.1151 | 0.0043 | 0.2469 | 0.6219 | 0.8930 | ns | ns | ns |
| ZT5-6 | 0.0163 | 0.7762 | 0.0018 | 0.2830 | 0.6890 | 0.1746 | ns | ns | ns |
| ZT6-7 | 0.0001 | 0.1505 | 0.0027 | 0.0010 | 0.4891 | 0.1983 | ** | ns | ns |
| ZT7-8 | 0.6738 | 0.4714 | 0.0147 | 0.0012 | 0.0017 | 0.0526 | ns | ns | ns |
| ZT8-9 | 0.9676 | 0.6733 | 0.8487 | 0.3096 | 0.4022 | 0.1828 | ns | ns | ns |
| ZT9-10 | 0.8522 | 0.2422 | 0.3108 | 0.0011 | 0.9952 | 0.6962 | ns | ns | ns |
| ZT10-11 | 0.9966 | 0.8254 | 0.0683 | 0.0039 | 0.8336 | <0.0001 | ns | ns | ns |
| ZT11-12 | 0.0268 | 0.2804 | 0.0678 | 0.9538 | 0.2620 | 0.8916 | ns | ns | ns |
| ZT12-13 | 0.2144 | 0.3250 | 0.8537 | 0.0083 | 0.0513 | 0.1085 | ns | ns | ns |
| ZT13-14 | 0.8022 | 0.5470 | 0.9997 | 0.7276 | 0.9974 | 0.9802 | ns | ns | ns |
| ZT14-15 | 0.0108 | 0.0149 | 0.0022 | 0.0005 | 0.5129 | 0.0040 | * | ns | ** |
| ZT15-16 | 0.0470 | <0.0001 | 0.2140 | 0.0015 | 0.0002 | 0.0010 | * | *** | ns |
| ZT16-17 | 0.0093 | 0.0029 | 0.0174 | 0.0003 | 0.0004 | 0.0009 | ** | ** | * |
| ZT17-18 | 0.1503 | 0.0012 | 0.0009 | 0.0008 | 0.0031 | <0.0001 | ns | ** | *** |
| ZT18-19 | 0.3069 | 0.0002 | 0.2621 | 0.0013 | <0.0001 | 0.0012 | ns | *** | ns |
| ZT19-20 | 0.7608 | 0.0005 | 0.4389 | <0.0001 | 0.0182 | 0.0005 | ns | * | ns |

| | | | | | | | | | |
|---|---|---|---|---|---|---|---|---|---|
| ZT20-21 | 0.9067 | 0.0016 | 0.1881 | 0.0002 | <0.0001 | 0.0001 | ns | ** | ns |
| ZT21-22 | 0.1312 | 0.0174 | 0.3207 | 0.0004 | 0.0008 | 0.0008 | ns | * | ns |
| ZT22-23 | 0.7829 | 0.4124 | 0.3113 | 0.0014 | 0.0055 | 0.0031 | ns | ns | ns |
| ZT23-24 | 0.9600 | 0.1246 | 0.6941 | 0.0003 | 0.0081 | 0.0009 | ns | ns | ns |
| ZT24-25 | 0.5007 | 0.5953 | 0.4108 | 0.0642 | 0.0241 | 0.0114 | ns | ns | ns |
| ZT25-26 | 0.1666 | 0.3557 | 0.6301 | 0.3275 | 0.2272 | 0.0052 | ns | ns | ns |
| ZT26-27 | 0.1584 | 0.2923 | 0.0050 | 0.7385 | 0.9887 | 0.9384 | ns | ns | ns |
| ZT27-28 | 0.5233 | 0.5943 | 0.5007 | 0.5311 | 0.6781 | 0.6761 | ns | ns | ns |
| ZT28-29 | 0.7872 | 0.1363 | 0.7345 | 0.0305 | 0.4297 | 0.2494 | ns | ns | ns |
| ZT29-30 | 0.4707 | 0.4440 | 0.9097 | <0.0001 | 0.9572 | 0.0419 | ns | ns | ns |
| ZT30-31 | 0.4530 | 0.4523 | 0.2749 | 0.0001 | 0.0805 | <0.0001 | ns | ns | ns |

| | | | Group | | Time | | Interaction | |
|---|---|---|---|---|---|---|---|---|
| | Data | DFn, DFd | F | p | F | p | F | p |
| Fig. 4C | Total sleep | 2, 113 | 21.95 | <0.0001 | 346.6 | <0.0001 | 16.94 | <0.0001 |
| | Sleep episodes | 2, 113 | 38.34 | <0.0001 | 13.33 | <0.0001 | 5.525 | <0.0001 |
| | P(wake) | 2, 112 | 7.799 | 0.0007 | 247.6 | <0.0001 | 11.82 | <0.0001 |

Dunnett's multiple comparisons

| | c316/$shi^{ts1}$ vs c316/+ | | | c316/$shi^{ts1}$ vs $shi^{ts1}$/+ | | | Summary | | |
|---|---|---|---|---|---|---|---|---|---|
| | Total sleep | Sleep episodes | P(wake) | Total sleep | Sleep episodes | P(wake) | Total sleep | Sleep episodes | P(wake) |
| ZT0-1 | 0.1048 | 0.1433 | 0.8009 | 0.0053 | 0.0490 | 0.1680 | ns | ns | ns |
| ZT1-2 | 0.1867 | 0.1312 | 0.0051 | 0.1948 | 0.1097 | 0.0001 | ns | ns | ** |
| ZT2-3 | 0.1621 | 0.1417 | 0.2292 | 0.2487 | 0.8214 | 0.0791 | ns | ns | ns |
| ZT3-4 | 0.1985 | 0.1068 | 0.2110 | 0.6183 | 0.9873 | 0.7170 | ns | ns | ns |
| ZT4-5 | 0.7876 | 0.0814 | 0.9244 | 0.0131 | 0.9763 | 0.0451 | ns | ns | ns |
| ZT5-6 | 0.0374 | 0.0008 | 0.9977 | 0.9417 | 0.0011 | 0.0011 | ns | ** | ns |
| ZT6-7 | <0.0001 | <0.0001 | <0.0001 | 0.5373 | 0.2511 | 0.0134 | ns | ns | * |
| ZT7-8 | 0.0244 | 0.1735 | <0.0001 | 0.0637 | 0.1970 | 0.5041 | ns | ns | ns |
| ZT8-9 | 0.9933 | 0.0032 | 0.5954 | <0.0001 | 0.0703 | 0.0461 | ns | ns | ns |
| ZT9-10 | 0.5792 | 0.7133 | 0.9481 | <0.0001 | 0.0308 | <0.0001 | ns | ns | ns |
| ZT10-11 | 0.9478 | 0.5248 | 0.8991 | <0.0001 | 0.9233 | <0.0001 | ns | ns | ns |
| ZT11-12 | 0.0028 | 0.0011 | 0.0042 | 0.7002 | 0.1342 | 0.1334 | ns | ns | ns |
| ZT12-13 | 0.1545 | 0.3935 | 0.1160 | 0.0986 | 0.2958 | 0.8508 | ns | ns | ns |
| ZT13-14 | 0.0174 | 0.0229 | 0.1881 | 0.0002 | 0.2108 | 0.0232 | * | ns | ns |
| ZT14-15 | 0.0002 | <0.0001 | <0.0001 | <0.0001 | 0.0002 | <0.0001 | *** | *** | **** |
| ZT15-16 | <0.0001 | <0.0001 | <0.0001 | 0.0003 | 0.0031 | 0.0004 | *** | ** | *** |
| ZT16-17 | 0.0002 | <0.0001 | <0.0001 | 0.0001 | <0.0001 | 0.0001 | *** | **** | *** |
| ZT17-18 | <0.0001 | <0.0001 | <0.0001 | 0.0008 | 0.0169 | 0.0001 | *** | * | *** |
| ZT18-19 | <0.0001 | <0.0001 | <0.0001 | 0.0003 | 0.0001 | 0.0007 | *** | *** | *** |
| ZT19-20 | <0.0001 | <0.0001 | <0.0001 | 0.0007 | 0.0011 | 0.0009 | *** | ** | *** |
| ZT20-21 | <0.0001 | <0.0001 | <0.0001 | 0.0001 | 0.0016 | 0.0003 | *** | ** | *** |
| ZT21-22 | 0.0137 | 0.0010 | 0.0005 | 0.0055 | 0.0032 | 0.0011 | * | ** | ** |
| ZT22-23 | 0.0045 | 0.0118 | 0.0711 | <0.0001 | 0.0008 | <0.0001 | ** | * | ns |
| ZT23-24 | 0.0127 | 0.0011 | 0.0025 | 0.0002 | 0.0005 | 0.0003 | * | ** | ** |
| ZT24-25 | 0.8349 | 0.3781 | 0.1102 | 0.3149 | 0.3449 | 0.0019 | ns | ns | ns |
| ZT25-26 | 0.0147 | 0.0113 | 0.0099 | 0.5632 | 0.0657 | 0.0407 | ns | ns | * |
| ZT26-27 | <0.0001 | 0.0001 | 0.0386 | 0.6998 | 0.0850 | 0.3568 | ns | ns | ns |
| ZT27-28 | 0.0095 | 0.0964 | 0.0063 | 0.2883 | 0.8494 | 0.9278 | ns | ns | ns |
| ZT28-29 | 0.0003 | 0.0028 | 0.0061 | 0.0001 | 0.9295 | 0.0663 | *** | ns | ns |

| | | | Group | | Time | | Interaction | |
|---|---|---|---|---|---|---|---|---|
| | Data | DFn, DFd | F | p | F | p | F | p |
| Fig. 4D | Total sleep | 2, 63 | 12.78 | <0.0001 | 104.1 | <0.0001 | 6.416 | <0.0001 |
| | Sleep episodes | 2, 63 | 11.13 | <0.0001 | 8.574 | <0.0001 | 5.329 | <0.0001 |
| | P(wake) | 2, 56 | 3.284 | 0.0448 | 68.15 | <0.0001 | 5.795 | <0.0001 |

Dunnett's multiple comparisons

| | VT064246/$shi^{ts1}$ vs VT064246/+ | | | VT064246/$shi^{ts1}$vs $shi^{ts1}$/+ | | | Summary | | |
|---|---|---|---|---|---|---|---|---|---|
| | Total sleep | Sleep episodes | P(wake) | Total sleep | Sleep episodes | P(wake) | Total sleep | Sleep episodes | P(wake) |
| ZT0-1 | 0.6014 | 0.1033 | 0.8663 | 0.1096 | 0.1622 | 0.2061 | ns | ns | ns |
| ZT1-2 | 0.8298 | 0.0350 | 0.8296 | 0.1747 | 0.9427 | 0.1743 | ns | ns | ns |
| ZT2-3 | 0.5753 | 0.1187 | 0.9827 | 0.4390 | 0.8938 | 0.1038 | ns | ns | ns |
| ZT3-4 | 0.0919 | 0.0350 | 0.4160 | >0.9999 | 0.1342 | 0.4869 | ns | ns | ns |
| ZT4-5 | 0.0232 | 0.9933 | 0.6256 | 0.0151 | 0.7952 | 0.9207 | * | ns | ns |
| ZT5-6 | 0.8659 | 0.9558 | 0.3036 | 0.2953 | 0.6880 | 0.5350 | ns | ns | ns |
| ZT6-7 | <0.0001 | 0.6634 | 0.0008 | 0.2530 | 0.2875 | 0.5392 | ns | ns | ns |
| ZT7-8 | 0.4548 | 0.4910 | 0.0004 | 0.0632 | 0.2992 | 0.6562 | ns | ns | ns |
| ZT8-9 | 0.0003 | 0.7733 | 0.2305 | 0.0014 | 0.4441 | 0.2841 | ** | ns | ns |
| ZT9-10 | <0.0001 | 0.0052 | 0.0122 | 0.0002 | 0.3562 | 0.0101 | *** | ns | * |

| | | | | | | | | | |
|---|---|---|---|---|---|---|---|---|---|
| ZT10-11 | <0.0001 | <0.0001 | 0.0258 | 0.2955 | 0.5178 | 0.4076 | ns | ns | ns |
| ZT11-12 | <0.0001 | 0.0002 | 0.0683 | 0.5733 | 0.9356 | 0.9960 | ns | ns | ns |
| ZT12-13 | 0.0002 | 0.0662 | 0.1071 | 0.2021 | 0.8383 | 0.6134 | ns | ns | ns |
| ZT13-14 | 0.0058 | 0.1391 | 0.0659 | 0.0515 | 0.1148 | 0.4398 | ns | ns | ns |
| ZT14-15 | 0.0020 | 0.1087 | 0.1218 | 0.0104 | 0.1525 | 0.2061 | * | ns | ns |
| ZT15-16 | 0.0008 | 0.0056 | 0.0835 | 0.0071 | 0.0106 | 0.0883 | ** | * | ns |
| ZT16-17 | 0.0035 | 0.0813 | 0.1588 | 0.0617 | 0.0130 | 0.1196 | ns | ns | ns |
| ZT17-18 | 0.0012 | 0.0392 | 0.1032 | 0.0093 | 0.0781 | 0.1483 | ** | ns | ns |
| ZT18-19 | 0.0215 | 0.0525 | 0.2687 | 0.0325 | 0.0466 | 0.0915 | * | ns | ns |
| ZT19-20 | 0.0395 | 0.1104 | 0.2266 | 0.1014 | 0.0598 | 0.1230 | ns | ns | ns |
| ZT20-21 | 0.1632 | 0.0208 | 0.4819 | 0.1634 | 0.0194 | 0.2061 | ns | * | ns |
| ZT21-22 | 0.1451 | 0.1005 | 0.7523 | 0.6687 | 0.0731 | 0.4734 | ns | ns | ns |
| ZT22-23 | 0.6272 | 0.1125 | 0.4157 | 0.9994 | 0.4994 | 0.4309 | ns | ns | ns |
| ZT23-24 | 0.0636 | 0.0111 | 0.5989 | 0.4181 | 0.0054 | 0.2601 | ns | * | ns |
| ZT24-25 | 0.0007 | 0.0004 | 0.2471 | 0.7169 | 0.6568 | 0.2810 | ns | ns | ns |
| ZT25-26 | 0.0005 | 0.0002 | 0.3645 | 0.5520 | 0.6667 | 0.2427 | ns | ns | ns |
| ZT26-27 | 0.0031 | 0.0326 | 0.3271 | 0.9418 | 0.5021 | 0.5079 | ns | ns | ns |
| ZT27-28 | 0.0045 | 0.0179 | 0.3211 | 0.4364 | 0.6784 | 0.5074 | ns | ns | ns |
| ZT28-29 | 0.0044 | 0.0946 | 0.3225 | 0.6624 | 0.9991 | 0.2044 | ns | ns | ns |

| | | | Group | | Time | | Interaction | |
|---|---|---|---|---|---|---|---|---|
| | Data | DFn, DFd | F | p | F | p | F | p |
| Fig. 5A | Total sleep | 2, 120 | 32.18 | ＜0.0001 | 343.8 | ＜0.0001 | 12.58 | ＜0.0001 |
| | Sleep episodes | 2, 120 | 27.39 | ＜0.0001 | 16.31 | ＜0.0001 | 5.367 | ＜0.0001 |
| | P(wake) | 2, 120 | 39.01 | ＜0.0001 | 264.0 | ＜0.0001 | 11.71 | ＜0.0001 |

Dunnett's multiple comparisons

| | R58E02/dTrpA1 vs R58E02/+ | | | R58E02/dTrpA1 vs dTrpA1/+ | | | Summary | | |
|---|---|---|---|---|---|---|---|---|---|
| | Total sleep | Sleep episodes | P(wake) | Total sleep | Sleep episodes | P(wake) | Total sleep | Sleep episodes | P(wake) |
| ZT0-1 | 0.9086 | 0.8154 | 0.5343 | 0.9342 | 0.9490 | 0.6511 | ns | ns | ns |
| ZT1-2 | 0.0383 | 0.0006 | 0.0511 | 0.8805 | 0.2848 | 0.9609 | ns | ns | ns |
| ZT2-3 | <0.0001 | <0.0001 | <0.0001 | 0.9143 | 0.9886 | 0.9971 | ns | ns | ns |
| ZT3-4 | <0.0001 | 0.0097 | <0.0001 | 0.9898 | 0.7700 | 0.6867 | ns | ns | ns |
| ZT4-5 | <0.0001 | 0.0165 | <0.0001 | 0.3169 | 0.5500 | 0.9712 | ns | ns | ns |
| ZT5-6 | <0.0001 | 0.9966 | <0.0001 | 0.0024 | 0.9747 | 0.0659 | ** | ns | ns |
| ZT6-7 | <0.0001 | <0.0001 | <0.0001 | <0.0001 | 0.0010 | 0.0004 | **** | *** | *** |
| ZT7-8 | <0.0001 | <0.0001 | <0.0001 | ＞0.9999 | 0.8920 | 0.9271 | ns | ns | ns |
| ZT8-9 | <0.0001 | <0.0001 | <0.0001 | 0.4656 | 0.7701 | 0.9989 | ns | ns | ns |
| ZT9-10 | <0.0001 | <0.0001 | <0.0001 | 0.9999 | 0.9275 | 0.7883 | ns | ns | ns |
| ZT10-11 | 0.0072 | 0.0086 | <0.0001 | 0.4619 | 0.5395 | 0.7640 | ns | ns | ns |
| ZT11-12 | 0.2348 | 0.0630 | 0.6876 | 0.0365 | 0.0433 | 0.1069 | ns | ns | ns |
| ZT12-13 | <0.0001 | 0.0010 | 0.0003 | 0.0001 | 0.0050 | 0.3617 | *** | ** | ns |
| ZT13-14 | <0.0001 | 0.4330 | <0.0001 | 0.0005 | 0.9703 | 0.0067 | *** | ns | ** |
| ZT14-15 | 0.0019 | 0.1080 | 0.0003 | 0.0004 | 0.0718 | 0.0001 | ** | ns | *** |
| ZT15-16 | 0.0058 | 0.2221 | 0.0081 | 0.0014 | 0.7349 | 0.0084 | ** | ns | ** |
| ZT16-17 | 0.0151 | 0.7221 | 0.0038 | 0.0155 | 0.2442 | 0.0349 | * | ns | * |
| ZT17-18 | 0.5671 | 0.9007 | 0.5788 | 0.4124 | 0.9989 | 0.3067 | ns | ns | ns |
| ZT18-19 | 0.9169 | 0.6717 | 0.4843 | 0.6928 | 0.8824 | 0.8054 | ns | ns | ns |
| ZT19-20 | 0.5228 | 0.7593 | 0.9883 | 0.3200 | 0.8915 | 0.0783 | ns | ns | ns |
| ZT20-21 | 0.2034 | 0.0700 | 0.6126 | 0.9719 | 0.3042 | 0.8387 | ns | ns | ns |
| ZT21-22 | 0.0373 | 0.9963 | 0.0364 | 0.4777 | 0.7408 | 0.2915 | ns | ns | ns |
| ZT22-23 | 0.2897 | 0.9932 | 0.0325 | 0.9445 | 0.6610 | 0.3055 | ns | ns | ns |
| ZT23-24 | 0.8547 | 0.6399 | 0.8373 | 0.6993 | 0.5527 | 0.5578 | ns | ns | ns |
| ZT24-25 | 0.0827 | 0.1046 | 0.7116 | 0.1276 | 0.4522 | 0.6013 | ns | ns | ns |
| ZT25-26 | 0.0188 | 0.0101 | 0.2142 | 0.8966 | 0.8637 | 0.9667 | ns | ns | ns |
| ZT26-27 | <0.0001 | 0.0001 | <0.0001 | 0.9692 | 0.9879 | 0.7914 | ns | ns | ns |
| ZT27-28 | <0.0001 | 0.0006 | <0.0001 | 0.7390 | 0.7811 | 0.9991 | ns | ns | ns |
| ZT28-29 | <0.0001 | 0.1204 | <0.0001 | 0.2086 | 0.4445 | 0.1619 | ns | ns | ns |

| | | | Group | | Time | | Interaction | |
|---|---|---|---|---|---|---|---|---|
| | Data | DFn, DFd | F | p | F | p | F | p |
| Fig. 5B | Total sleep | 2, 85 | 0.8538 | 0.4294 | 145.1 | ＜0.0001 | 2.800 | ＜0.0001 |
| | Sleep episodes | 2, 85 | 10.55 | ＜0.0001 | 16.12 | ＜0.0001 | 1.593 | ＜0.0001 |
| | P(wake) | 2, 85 | 0.1322 | 0.8764 | 122.3 | ＜0.0001 | 3.910 | ＜0.0001 |

Dunnett's multiple comparisons

| | R58E02/$shi^{ts1}$ vs R58E02/+ | | | R58E02/$shi^{ts1}$ vs $shi^{ts1}$/+ | | | Summary | | |
|---|---|---|---|---|---|---|---|---|---|
| | Total sleep | Sleep episodes | P(wake) | Total sleep | Sleep episodes | P(wake) | Total sleep | Sleep episodes | P(wake) |
| ZT0-1 | 0.8005 | 0.2826 | 0.0435 | 0.4019 | 0.8339 | 0.0771 | ns | ns | ns |
| ZT1-2 | 0.9382 | 0.1464 | 0.0793 | 0.1645 | 0.4917 | 0.1169 | ns | ns | ns |

| | | | | | | | | | |
|---|---|---|---|---|---|---|---|---|---|
| ZT2-3 | 0.1552 | 0.0021 | 0.3853 | 0.7564 | 0.1270 | 0.7221 | ns | ns | ns |
| ZT3-4 | 0.3085 | 0.1483 | 0.1580 | 0.3739 | 0.5286 | 0.5679 | ns | ns | ns |
| ZT4-5 | 0.0856 | 0.4209 | 0.1641 | 0.0247 | 0.7966 | 0.0538 | ns | ns | ns |
| ZT5-6 | 0.7124 | 0.6537 | 0.2077 | 0.2381 | 0.7568 | 0.0337 | ns | ns | ns |
| ZT6-7 | 0.2565 | 0.4993 | 0.8119 | 0.0179 | 0.9113 | 0.1739 | ns | ns | ns |
| ZT7-8 | 0.9895 | 0.9558 | 0.2751 | 0.7794 | 0.1494 | 0.0590 | ns | ns | ns |
| ZT8-9 | 0.9689 | 0.9665 | 0.9934 | 0.9968 | 0.9880 | 0.9565 | ns | ns | ns |
| ZT9-10 | 0.0042 | >0.9999 | 0.609 | 0.4358 | 0.8578 | 0.7616 | ns | ns | ns |
| ZT10-11 | 0.0331 | 0.5824 | 0.0017 | 0.3377 | 0.1917 | 0.5663 | ns | ns | ns |
| ZT11-12 | 0.9911 | 0.0647 | 0.5694 | 0.3057 | 0.7984 | 0.0155 | ns | ns | ns |
| ZT12-13 | 0.2987 | 0.9709 | 0.7210 | 0.2382 | 0.6114 | 0.1945 | ns | ns | ns |
| ZT13-14 | 0.5461 | 0.8163 | 0.6356 | 0.3262 | 0.2385 | 0.9144 | ns | ns | ns |
| ZT14-15 | 0.4584 | 0.8238 | 0.3040 | 0.2050 | 0.0029 | 0.4159 | ns | ns | ns |
| ZT15-16 | 0.6002 | 0.9381 | 0.7440 | 0.1588 | 0.0311 | 0.3794 | ns | ns | ns |
| ZT16-17 | 0.8609 | 0.4353 | 0.3110 | 0.0467 | 0.0002 | 0.0432 | ns | ns | ns |
| ZT17-18 | 0.3560 | 0.2718 | 0.3548 | 0.0095 | 0.0114 | 0.0272 | ns | ns | ns |
| ZT18-19 | 0.9652 | 0.3089 | 0.2594 | 0.0750 | 0.0064 | 0.0130 | ns | ns | ns |
| ZT19-20 | 0.6333 | 0.8799 | 0.6772 | 0.4020 | 0.1600 | 0.1175 | ns | ns | ns |
| ZT20-21 | 0.5231 | 0.8895 | 0.5613 | 0.3095 | 0.1607 | 0.9982 | ns | ns | ns |
| ZT21-22 | 0.0171 | 0.9988 | 0.1173 | 0.4731 | 0.0465 | 0.3027 | ns | ns | ns |
| ZT22-23 | 0.1985 | 0.6118 | 0.0365 | 0.2963 | 0.1331 | 0.8238 | ns | ns | ns |
| ZT23-24 | 0.9784 | 0.3963 | 0.8772 | 0.4226 | 0.0608 | 0.3506 | ns | ns | ns |
| ZT24-25 | 0.3422 | 0.7435 | 0.8284 | 0.1850 | 0.1291 | 0.5364 | ns | ns | ns |
| ZT25-26 | 0.7121 | 0.9965 | 0.8877 | 0.0971 | 0.0168 | 0.0378 | ns | ns | ns |
| ZT26-27 | 0.1224 | 0.4326 | 0.6606 | 0.1271 | 0.9026 | 0.2283 | ns | ns | ns |
| ZT27-28 | 0.6677 | 0.0765 | 0.6609 | 0.9972 | 0.1588 | 0.9780 | ns | ns | ns |
| ZT28-29 | 0.1900 | 0.2959 | 0.2459 | 0.9980 | 0.9821 | 0.7576 | ns | ns | ns |
| ZT29-30 | 0.8075 | 0.2680 | 0.3006 | 0.9481 | 0.5323 | 0.9075 | ns | ns | ns |
| ZT30-31 | 0.2392 | 0.2211 | 0.2012 | 0.9997 | 0.9988 | 0.9419 | ns | ns | ns |

| | Data | DFn, DFd | Group F | Group p | Time F | Time p | Interaction F | Interaction p |
|---|---|---|---|---|---|---|---|---|
| Fig. 5C | Total sleep | 2, 109 | 40.01 | <0.0001 | 305.9 | <0.0001 | 12.03 | <0.0001 |
| | Sleep episodes | 2, 109 | 17.56 | <0.0001 | 27.15 | <0.0001 | 5.9780 | <0.0001 |
| | P(wake) | 2, 109 | 32.74 | <0.0001 | 252.9 | <0.0001 | 14.91 | <0.0001 |

Dunnett's multiple comparisons

| | c316/shi$^{ts1}$ vs c316/+ | | | c316/shi$^{ts1}$ vs shi$^{ts1}$/+ | | | Summary | | |
|---|---|---|---|---|---|---|---|---|---|
| | Total sleep | Sleep episodes | P(wake) | Total sleep | Sleep episodes | P(wake) | Total sleep | Sleep episodes | P(wake) |
| ZT0-1 | 0.2363 | 0.0786 | 0.1531 | 0.0893 | 0.0645 | 0.0042 | ns | ns | ns |
| ZT1-2 | 0.0073 | 0.0160 | 0.0121 | 0.6278 | 0.4555 | 0.0622 | ns | ns | ns |
| ZT2-3 | 0.0030 | 0.0012 | 0.0001 | 0.8066 | 0.8262 | 0.6792 | ns | ns | ns |
| ZT3-4 | 0.0025 | 0.0049 | <0.0001 | 0.0003 | 0.0889 | 0.0111 | ** | ns | * |
| ZT4-5 | 0.0068 | 0.2228 | <0.0001 | <0.0001 | 0.0416 | <0.0001 | ** | ns | **** |
| ZT5-6 | 0.0059 | 0.2774 | 0.0203 | 0.0015 | 0.0047 | 0.0003 | ** | ns | * |
| ZT6-7 | 0.0004 | 0.1944 | <0.0001 | 0.1674 | 0.9849 | 0.1763 | ns | ns | ns |
| ZT7-8 | 0.6153 | 0.5574 | 0.0191 | 0.9404 | 0.9968 | 0.1813 | ns | ns | ns |
| ZT8-9 | 0.0221 | 0.4296 | 0.4767 | 0.0083 | 0.0001 | 0.2909 | * | ns | ns |
| ZT9-10 | 0.1397 | 0.1204 | 0.0040 | <0.0001 | <0.0001 | <0.0001 | ns | ns | ** |
| ZT10-11 | 0.0312 | 0.0256 | 0.0018 | 0.0004 | 0.0200 | <0.0001 | * | * | ** |
| ZT11-12 | 0.6914 | 0.7137 | 0.0019 | 0.9579 | 0.9744 | 0.0140 | ns | ns | * |
| ZT12-13 | 0.0187 | 0.1381 | 0.4596 | <0.0001 | 0.0398 | 0.0417 | * | ns | ns |
| ZT13-14 | 0.0414 | 0.3252 | 0.0279 | <0.0001 | 0.0304 | <0.0001 | * | ns | * |
| ZT14-15 | 0.7500 | 0.1517 | 0.2397 | <0.0001 | 0.0004 | 0.0010 | ns | ns | ns |
| ZT15-16 | 0.7791 | 0.7682 | 0.9804 | 0.0004 | 0.0156 | 0.0017 | ns | ns | ns |
| ZT16-17 | 0.4514 | 0.3751 | 0.6402 | <0.0001 | <0.0001 | 0.0002 | ns | ns | ns |
| ZT17-18 | 0.0528 | 0.1956 | 0.1104 | <0.0001 | 0.0012 | <0.0001 | ns | ns | ns |
| ZT18-19 | 0.1888 | 0.0408 | 0.5773 | 0.0016 | 0.0019 | 0.0029 | ns | * | ns |
| ZT19-20 | 0.5402 | 0.5251 | 0.2536 | 0.0100 | 0.0078 | 0.0151 | ns | ns | ns |
| ZT20-21 | 0.2873 | 0.1806 | 0.9350 | 0.1306 | 0.1345 | 0.0418 | ns | ns | ns |
| ZT21-22 | 0.6358 | 0.4027 | 0.2554 | 0.0108 | 0.1027 | 0.0351 | ns | ns | ns |
| ZT22-23 | 0.1032 | 0.1278 | 0.6022 | 0.0018 | 0.0143 | 0.0180 | ns | ns | ns |
| ZT23-24 | 0.0353 | 0.1643 | 0.3655 | 0.0212 | 0.1027 | 0.0361 | * | ns | ns |
| ZT24-25 | 0.1111 | 0.0691 | 0.0423 | 0.1887 | 0.2912 | 0.0167 | ns | ns | * |
| ZT25-26 | 0.0023 | 0.0008 | 0.0002 | 0.0213 | 0.0116 | 0.0066 | * | * | ** |
| ZT26-27 | 0.0037 | 0.0059 | 0.0003 | >0.9999 | 0.9432 | 0.0291 | ns | ns | * |
| ZT27-28 | 0.0028 | 0.0587 | <0.0001 | 0.0008 | 0.3171 | 0.1213 | ** | ns | ns |
| ZT28-29 | 0.0002 | 0.0093 | <0.0001 | 0.0105 | 0.8054 | 0.0022 | * | ns | ns |

| | | | Group | | Time | | Interaction |
|---|---|---|---|---|---|---|---|

| | Data | DFn, DFd | F | p | F | p | F | p |
|---|---|---|---|---|---|---|---|---|
| Fig. 5D | Total sleep | 2, 120 | 10.69 | <0.0001 | 366.0 | <0.0001 | 3.9710 | <0.0001 |
| | Sleep episodes | 2, 120 | 1.6140 | 0.2034 | 32.36 | <0.0001 | 2.8770 | <0.0001 |
| | P(wake) | 2, 108 | 9.6600 | 0.0001 | 275.7 | <0.0001 | 2.8450 | <0.0001 |

Dunnett's multiple comparisons

| | c316/dTrpA1 vs c316/+ | | | c316/dTrpA1 vs dTrpA1/+ | | | Summary | | |
|---|---|---|---|---|---|---|---|---|---|
| | Total sleep | Sleep episodes | P(wake) | Total sleep | Sleep episodes | P(wake) | Total sleep | Sleep episodes | P(wake) |
| ZT0-1 | 0.3626 | 0.0391 | 0.8978 | 0.9918 | >0.9999 | 0.0111 | ns | ns | ns |
| ZT1-2 | 0.9545 | 0.9799 | 0.9973 | 0.5452 | 0.6784 | 0.1896 | ns | ns | ns |
| ZT2-3 | 0.9474 | 0.9727 | 0.0966 | 0.4805 | 0.6537 | 0.9961 | ns | ns | ns |
| ZT3-4 | 0.9316 | 0.8585 | 0.9857 | 0.1698 | 0.9464 | 0.0398 | ns | ns | ns |
| ZT4-5 | 0.4767 | 0.9982 | 0.9864 | 0.0028 | 0.3100 | 0.0016 | ns | ns | ns |
| ZT5-6 | 0.2708 | 0.6247 | 0.8270 | 0.0008 | 0.4900 | 0.0022 | ns | ns | ns |
| ZT6-7 | 0.4385 | 0.9366 | 0.7885 | <0.0001 | 0.9876 | 0.0631 | ns | ns | ns |
| ZT7-8 | 0.0021 | 0.0587 | 0.0113 | 0.0257 | 0.0587 | 0.077 | * | ns | ns |
| ZT8-9 | 0.5146 | 0.4169 | 0.8064 | 0.2170 | 0.5753 | 0.3378 | ns | ns | ns |
| ZT9-10 | 0.0518 | 0.0213 | 0.0871 | 0.7524 | 0.9972 | 0.8517 | ns | ns | ns |
| ZT10-11 | 0.0626 | 0.0877 | 0.0092 | 0.7672 | 0.7835 | 0.9346 | ns | ns | ns |
| ZT11-12 | 0.0121 | 0.1292 | 0.7628 | 0.9415 | 0.7839 | 0.2535 | ns | ns | ns |
| ZT12-13 | 0.0125 | 0.0010 | 0.0116 | 0.0118 | 0.0097 | 0.0027 | * | ** | * |
| ZT13-14 | 0.1297 | 0.3798 | 0.2562 | <0.0001 | 0.2680 | 0.0246 | ns | ns | ns |
| ZT14-15 | 0.9200 | 0.9719 | 0.1479 | <0.0001 | 0.0002 | 0.0124 | ns | ns | ns |
| ZT15-16 | 0.7227 | 0.6478 | 0.0932 | 0.0046 | 0.0021 | 0.9846 | ns | ns | ns |
| ZT16-17 | 0.9513 | 0.2125 | 0.3510 | 0.0581 | 0.0258 | 0.9999 | ns | ns | ns |
| ZT17-18 | 0.4718 | 0.2965 | 0.1772 | 0.9854 | 0.3905 | 0.8107 | ns | ns | ns |
| ZT18-19 | 0.4065 | 0.9803 | 0.1455 | 0.5511 | 0.0580 | 0.6864 | ns | ns | ns |
| ZT19-20 | 0.2573 | 0.1721 | 0.8705 | 0.6872 | 0.3212 | 0.6427 | ns | ns | ns |
| ZT20-21 | 0.9307 | 0.9430 | 0.8212 | 0.9755 | 0.5198 | 0.8757 | ns | ns | ns |
| ZT21-22 | 0.9996 | 0.8567 | 0.9889 | 0.9853 | 0.8890 | 0.9376 | ns | ns | ns |
| ZT22-23 | 0.6563 | 0.7556 | 0.8949 | 0.9631 | 0.9708 | 0.8445 | ns | ns | ns |
| ZT23-24 | 0.9722 | 0.8131 | 0.8383 | 0.9392 | 0.9815 | 0.9057 | ns | ns | ns |
| ZT24-25 | 0.3406 | 0.3063 | 0.2709 | 0.9494 | 0.9450 | 0.0613 | ns | ns | ns |
| ZT25-26 | 0.8390 | 0.9893 | 0.4518 | 0.4362 | 0.2941 | 0.1113 | ns | ns | ns |
| ZT26-27 | 0.9759 | 0.8060 | 0.9634 | 0.1469 | 0.1217 | 0.0861 | ns | ns | ns |
| ZT27-28 | 0.9128 | 0.7156 | 0.5508 | 0.0284 | 0.1679 | 0.001 | ns | ns | ns |
| ZT28-29 | 0.9916 | 0.6140 | 0.9656 | 0.0003 | 0.0173 | 0.0050 | ns | ns | ns |
| ZT29-30 | 0.2575 | 0.6352 | 0.7387 | 0.0025 | 0.8587 | 0.0085 | ns | ns | ns |
| ZT30-31 | 0.9980 | 0.7240 | 0.9954 | 0.0505 | 0.4597 | 0.0175 | ns | ns | ns |

| | | | Group | | Time | | Interaction | |
|---|---|---|---|---|---|---|---|---|
| | Data | DFn, DFd | F | p | F | p | F | p |
| Fig. 7H | Total sleep | 2, 76 | 8.388 | 0.0005 | 169.2 | <0.0001 | 3.7640 | <0.0001 |
| | Sleep episodes | 2, 76 | 4.602 | 0.0130 | 17.64 | <0.0001 | 2.9960 | <0.0001 |
| | P(wake) | 2, 76 | 3.148 | 0.0486 | 131.5 | <0.0001 | 2.7130 | <0.0001 |

Dunnett's multiple comparisons

| | MB299B/dTrpA1 vs MB299B/+ | | | MB299B/dTrpA1 vs dTrpA1/+ | | | Summary | | |
|---|---|---|---|---|---|---|---|---|---|
| | Total sleep | Sleep episodes | P(wake) | Total sleep | Sleep episodes | P(wake) | Total sleep | Sleep episodes | P(wake) |
| ZT0-1 | 0.9008 | 0.7471 | - | 0.3210 | 0.9972 | - | ns | ns | - |
| ZT1-2 | 0.1209 | 0.0056 | 0.0438 | 0.8648 | >0.9999 | 0.6962 | ns | ns | ns |
| ZT2-3 | 0.4323 | 0.2669 | 0.3740 | 0.9895 | 0.9957 | 0.8727 | ns | ns | ns |
| ZT3-4 | 0.9515 | 0.3079 | 0.8372 | 0.9240 | 0.9397 | 0.9880 | ns | ns | ns |
| ZT4-5 | 0.0994 | 0.5627 | 0.9481 | 0.0015 | 0.3609 | 0.9998 | ns | ns | ns |
| ZT5-6 | 0.1848 | 0.5773 | 0.3472 | <0.0001 | 0.1328 | 0.0002 | ns | ns | ns |
| ZT6-7 | 0.0675 | 0.0338 | 0.2039 | <0.0001 | 0.0343 | <0.0001 | ns | * | ns |
| ZT7-8 | 0.1650 | 0.9985 | 0.6571 | 0.1190 | 0.2494 | <0.0001 | ns | ns | ns |
| ZT8-9 | 0.1921 | 0.3105 | 0.9855 | 0.0015 | 0.1149 | 0.0515 | ns | ns | ns |
| ZT9-10 | 0.4766 | 0.1086 | 0.6598 | 0.8071 | 0.9734 | 0.9204 | ns | ns | ns |
| ZT10-11 | 0.9234 | 0.8170 | 0.8971 | 0.4374 | 0.1497 | 0.9152 | ns | ns | ns |
| ZT11-12 | 0.7611 | 0.9832 | 0.8104 | 0.4730 | 0.3304 | 0.6678 | ns | ns | ns |
| ZT12-13 | 0.4141 | 0.3131 | 0.3499 | 0.9966 | 0.5251 | 0.9479 | ns | ns | ns |
| ZT13-14 | 0.1063 | 0.1120 | 0.2001 | 0.0129 | 0.0788 | 0.0673 | ns | ns | ns |
| ZT14-15 | 0.8599 | 0.3156 | 0.3850 | 0.0029 | 0.0075 | 0.0057 | ns | ns | ns |
| ZT15-16 | 0.9441 | 0.0110 | 0.9094 | 0.0010 | 0.0010 | 0.0083 | ns | * | ns |
| ZT16-17 | 0.6404 | 0.0068 | 0.7756 | 0.0020 | 0.0001 | 0.0075 | ns | ** | ns |
| ZT17-18 | 0.0416 | 0.0037 | 0.0687 | 0.0002 | <0.0001 | <0.0001 | * | ** | ns |
| ZT18-19 | 0.0226 | 0.6235 | 0.4094 | 0.0004 | 0.0149 | 0.0097 | * | ns | ns |
| ZT19-20 | 0.0010 | 0.0007 | 0.0055 | 0.0008 | <0.0001 | 0.0009 | *** | *** | ** |
| ZT20-21 | 0.0016 | 0.0003 | 0.0020 | 0.0458 | <0.0001 | 0.0098 | * | *** | ** |

| | | | | | | | | | |
|---|---|---|---|---|---|---|---|---|---|
| ZT21-22 | 0.0097 | 0.0198 | 0.0025 | 0.0188 | 0.0016 | 0.0356 | * | * | * |
| ZT22-23 | 0.0584 | 0.0050 | 0.0526 | 0.0412 | 0.0375 | 0.1694 | ns | * | ns |
| ZT23-24 | 0.1141 | 0.7781 | 0.3733 | 0.1519 | 0.1133 | 0.1542 | ns | ns | ns |
| ZT24-25 | 0.6262 | 0.8891 | 0.9961 | 0.7042 | 0.9609 | 0.5279 | ns | ns | ns |
| ZT25-26 | 0.5513 | 0.8899 | 0.6711 | 0.4102 | 0.3740 | 0.9370 | ns | ns | ns |
| ZT26-27 | 0.7146 | 0.9669 | 0.9096 | 0.7726 | 0.7684 | 0.9991 | ns | ns | ns |
| ZT27-28 | 0.2084 | 0.2886 | 0.1680 | 0.9489 | 0.4794 | 0.9659 | ns | ns | ns |
| ZT28-29 | 0.1365 | 0.3233 | 0.5605 | 0.4382 | 0.8724 | 0.9521 | ns | ns | ns |
| ZT29-30 | 0.8415 | 0.9937 | 0.4147 | 0.1906 | 0.9938 | 0.3155 | ns | ns | ns |
| ZT30-31 | 0.4714 | 0.8791 | 0.7649 | 0.0561 | 0.9025 | 0.4026 | ns | ns | ns |

| | | | Group | | Time | | Interaction | |
|---|---|---|---|---|---|---|---|---|
| | Data | DFn, DFd | F | p | F | p | F | p |
| Fig. 7I | Total sleep | 2, 103 | 2.4680 | 0.0897 | 294.4 | <0.0001 | 9.8710 | <0.0001 |
| | Sleep episodes | 2, 103 | 8.9860 | 0.0003 | 19.13 | <0.0001 | 3.5190 | <0.0001 |
| | P(wake) | 2, 103 | 1.3280 | 0.2696 | 213.7 | <0.0001 | 9.2830 | <0.0001 |

Dunnett's multiple comparisons

| | MB299B/$shi^{ts1}$ vs MB299B/+ | | | MB299B/$shi^{ts1}$ vs $shi^{ts1}$/+ | | | Summary | | |
|---|---|---|---|---|---|---|---|---|---|
| | Total sleep | Sleep episodes | P(wake) | Total sleep | Sleep episodes | P(wake) | Total sleep | Sleep episodes | P(wake) |
| ZT0-1 | 0.5453 | 0.9967 | 0.9595 | 0.0046 | 0.1744 | 0.0008 | ns | ns | ns |
| ZT1-2 | 0.9548 | 0.9576 | 0.9559 | 0.4535 | 0.2436 | 0.0031 | ns | ns | ns |
| ZT2-3 | 0.8636 | 0.9626 | 0.0245 | 0.9724 | 0.9883 | 0.6837 | ns | ns | ns |
| ZT3-4 | 0.8184 | 0.9767 | 0.3053 | 0.8970 | 0.9319 | 0.7827 | ns | ns | ns |
| ZT4-5 | 0.8556 | 0.7185 | 0.8448 | 0.4840 | 0.9298 | 0.8280 | ns | ns | ns |
| ZT5-6 | 0.9404 | 0.9690 | 0.3321 | 0.0234 | 0.0760 | 0.0603 | ns | ns | ns |
| ZT6-7 | 0.1678 | 0.3358 | 0.3128 | <0.0001 | 0.0021 | 0.0216 | ns | ns | ns |
| ZT7-8 | 0.0003 | <0.0001 | 0.3743 | 0.9471 | 0.3120 | 0.0008 | ns | ns | ns |
| ZT8-9 | <0.0001 | 0.9511 | <0.0001 | 0.4955 | 0.6085 | 0.4937 | ns | ns | ns |
| ZT9-10 | >0.9999 | >0.9999 | 0.4020 | <0.0001 | 0.1663 | 0.0455 | ns | ns | ns |
| ZT10-11 | 0.0709 | 0.1667 | 0.3092 | 0.5090 | 0.0922 | <0.0001 | ns | ns | ns |
| ZT11-12 | 0.7331 | >0.9999 | 0.8686 | 0.5517 | 0.1070 | 0.0468 | ns | ns | ns |
| ZT12-13 | 0.5493 | 0.3579 | 0.0802 | 0.0011 | 0.0111 | 0.1171 | ns | ns | ns |
| ZT13-14 | 0.6080 | 0.0061 | 0.3821 | 0.0269 | 0.9540 | 0.0093 | ns | ns | ns |
| ZT14-15 | 0.3373 | 0.0355 | 0.3100 | 0.8050 | 0.3550 | 0.4580 | ns | ns | ns |
| ZT15-16 | 0.3016 | 0.0022 | 0.0626 | 0.9605 | 0.9818 | 0.8952 | ns | ns | ns |
| ZT16-17 | 0.0910 | 0.0106 | 0.0879 | 0.9966 | 0.3719 | 0.8301 | ns | ns | ns |
| ZT17-18 | 0.0504 | 0.0047 | 0.0209 | 0.9989 | 0.7792 | 0.3368 | ns | ns | ns |
| ZT18-19 | 0.0113 | 0.0060 | 0.0317 | 0.3184 | 0.4735 | 0.4365 | ns | ns | ns |
| ZT19-20 | 0.0149 | 0.0032 | 0.0095 | 0.2424 | >0.9999 | 0.1001 | ns | ns | ns |
| ZT20-21 | 0.2533 | 0.0386 | 0.0144 | 0.2765 | 0.7969 | 0.1505 | ns | ns | ns |
| ZT21-22 | 0.2597 | 0.3544 | 0.1959 | 0.9943 | 0.8940 | 0.9809 | ns | ns | ns |
| ZT22-23 | 0.0024 | 0.0394 | 0.0985 | 0.3160 | 0.6696 | 0.6003 | ns | ns | ns |
| ZT23-24 | 0.0170 | 0.0596 | 0.0096 | 0.1281 | 0.9342 | 0.0796 | ns | ns | ns |
| ZT24-25 | 0.8929 | 0.9648 | 0.4735 | 0.0238 | 0.0136 | 0.1013 | ns | ns | ns |
| ZT25-26 | 0.9853 | 0.9994 | 0.5683 | 0.0619 | 0.0246 | 0.0002 | ns | ns | ns |
| ZT26-27 | 0.9853 | 0.2475 | 0.8228 | 0.9812 | 0.2330 | 0.0617 | ns | ns | ns |
| ZT27-28 | 0.9771 | 0.7857 | 0.4495 | 0.2291 | 0.2123 | 0.7262 | ns | ns | ns |
| ZT28-29 | 0.3856 | 0.9034 | 0.9803 | 0.0176 | 0.2097 | 0.0461 | ns | ns | ns |
| ZT29-30 | 0.9851 | 0.6283 | 0.9927 | <0.0001 | 0.1686 | <0.0001 | ns | ns | ns |
| ZT30-31 | 0.9122 | 0.3080 | 0.9723 | <0.0001 | <0.0001 | <0.0001 | ns | ns | ns |

| | | | Group | | Time | | Interaction | |
|---|---|---|---|---|---|---|---|---|
| | Data | DFn, DFd | F | p | F | p | F | p |
| Fig. 7J | Total sleep | 2, 79 | 18.94 | <0.0001 | 178.8 | <0.0001 | 8.6940 | <0.0001 |
| | Sleep episodes | 2, 79 | 19.82 | <0.0001 | 16.39 | <0.0001 | 3.1660 | <0.0001 |
| | P(wake) | 2, 78 | 8.348 | 0.0005 | 137 | <0.0001 | 5.6740 | <0.0001 |

Dunnett's multiple comparisons

| | MB043B/dTrpA1 vs MB043B/+ | | | MB043B/dTrpA1 vs dTrpA1/+ | | | Summary | | |
|---|---|---|---|---|---|---|---|---|---|
| | Total sleep | Sleep episodes | P(wake) | Total sleep | Sleep episodes | P(wake) | Total sleep | Sleep episodes | P(wake) |
| ZT0-1 | 0.0012 | 0.0129 | - | 0.4400 | 0.7524 | - | ns | ns | - |
| ZT1-2 | 0.0032 | 0.0029 | 0.0187 | 0.0697 | 0.1516 | 0.4706 | ns | ns | ns |
| ZT2-3 | 0.0036 | 0.0019 | 0.0383 | 0.5696 | 0.8560 | 0.1921 | ns | ns | ns |
| ZT3-4 | 0.0023 | 0.0041 | 0.0498 | 0.1846 | 0.1069 | 0.5094 | ns | ns | ns |
| ZT4-5 | 0.8898 | 0.7974 | 0.0507 | <0.0001 | 0.9670 | 0.3653 | ns | ns | ns |
| ZT5-6 | 0.3382 | 0.0357 | 0.5267 | <0.0001 | 0.2234 | <0.0001 | ns | ns | ns |
| ZT6-7 | 0.0015 | 0.0618 | 0.0789 | <0.0001 | 0.0620 | <0.0001 | ** | ns | ns |
| ZT7-8 | 0.9979 | 0.3545 | 0.7467 | 0.0012 | 0.3976 | <0.0001 | ns | ns | ns |
| ZT8-9 | 0.0076 | 0.1832 | 0.2104 | <0.0001 | 0.9648 | 0.0025 | ** | ns | ns |

| | | | | | | | | | |
|---|---|---|---|---|---|---|---|---|---|
| ZT9-10 | 0.8015 | 0.0893 | 0.8603 | 0.0517 | 0.3482 | 0.0090 | ns | ns | ns |
| ZT10-11 | 0.0008 | 0.0024 | 0.1810 | 0.9587 | 0.8682 | 0.7601 | ns | ns | ns |
| ZT11-12 | 0.0279 | 0.0354 | 0.1140 | 0.7040 | 0.4804 | 0.9664 | ns | ns | ns |
| ZT12-13 | 0.5118 | 0.7019 | 0.5201 | 0.0638 | 0.4647 | 0.9425 | ns | ns | ns |
| ZT13-14 | 0.0491 | 0.0836 | 0.0259 | 0.0020 | 0.9500 | 0.0062 | * | ns | * |
| ZT14-15 | 0.0002 | 0.0038 | 0.0213 | 0.0002 | 0.0811 | 0.0093 | *** | ns | * |
| ZT15-16 | 0.0108 | 0.0147 | 0.0053 | 0.0003 | 0.0083 | 0.0026 | * | * | ** |
| ZT16-17 | 0.0155 | 0.0064 | 0.0830 | 0.0003 | 0.0011 | 0.0108 | * | ** | ns |
| ZT17-18 | 0.0028 | <0.0001 | 0.0059 | 0.0002 | 0.0002 | 0.0002 | ** | *** | ** |
| ZT18-19 | 0.0007 | 0.0014 | 0.0154 | 0.0002 | 0.0011 | 0.0014 | *** | ** | * |
| ZT19-20 | 0.0019 | 0.0019 | 0.0011 | 0.0002 | 0.0016 | 0.0006 | ** | ** | ** |
| ZT20-21 | 0.0006 | 0.0020 | 0.0034 | 0.0032 | 0.0034 | 0.0063 | ** | ** | ** |
| ZT21-22 | 0.0005 | 0.0046 | 0.0023 | 0.0019 | 0.0095 | 0.0076 | ** | ** | ** |
| ZT22-23 | 0.0002 | 0.0008 | 0.0010 | 0.0005 | 0.0298 | 0.0077 | *** | * | ** |
| ZT23-24 | 0.0007 | 0.0010 | 0.0019 | 0.0095 | 0.0182 | 0.0051 | ** | * | ** |
| ZT24-25 | 0.0112 | 0.0016 | 0.0078 | 0.9656 | 0.9991 | 0.0116 | ns | ns | * |
| ZT25-26 | 0.0065 | 0.0067 | 0.0952 | 0.6850 | 0.7371 | 0.5757 | ns | ns | ns |
| ZT26-27 | 0.0076 | 0.3405 | 0.0501 | 0.0352 | 0.4913 | 0.4102 | * | ns | ns |
| ZT27-28 | 0.0018 | 0.1981 | 0.0123 | 0.0628 | 0.9391 | 0.1087 | ns | ns | ns |
| ZT28-29 | 0.0089 | 0.4440 | 0.1439 | 0.8539 | 0.7427 | 0.8064 | ns | ns | ns |
| ZT29-30 | 0.0633 | 0.9574 | 0.0568 | 0.3596 | 0.4635 | 0.8512 | ns | ns | ns |
| ZT30-31 | 0.4448 | 0.4818 | 0.7767 | 0.7838 | 0.4160 | 0.2171 | ns | ns | ns |

| | | | Group | | Time | | Interaction | |
|---|---|---|---|---|---|---|---|---|
| | Data | DFn, DFd | F | p | F | p | F | p |
| | Total sleep | 2, 100 | 29.75 | <0.0001 | 214.3 | <0.0001 | 11.17 | <0.0001 |
| Fig. 7K | Sleep episodes | 2, 100 | 13.18 | <0.0001 | 18.11 | <0.0001 | 5.3760 | <0.0001 |
| | P(wake) | 2, 101 | 10.23 | <0.0001 | 166.8 | <0.0001 | 9.9810 | <0.0001 |

Dunnett's multiple comparisons

| | MB043B/$shi^{ts1}$ vs MB043B/+ | | | MB043B/$shi^{ts1}$ vs $shi^{ts1}$/+ | | | Summary | | |
|---|---|---|---|---|---|---|---|---|---|
| | Total sleep | Sleep episodes | P(wake) | Total sleep | Sleep episodes | P(wake) | Total sleep | Sleep episodes | P(wake) |
| ZT0-1 | 0.7629 | 0.8750 | 0.8152 | 0.0292 | 0.3556 | 0.0004 | ns | ns | ns |
| ZT1-2 | 0.2073 | 0.0745 | 0.9994 | 0.5546 | 0.3382 | 0.0505 | ns | ns | ns |
| ZT2-3 | 0.3581 | 0.5553 | 0.0235 | 0.4590 | 0.5238 | 0.1837 | ns | ns | ns |
| ZT3-4 | 0.0523 | 0.0154 | 0.2194 | 0.0142 | 0.0143 | 0.3313 | ns | * | ns |
| ZT4-5 | 0.0198 | 0.1146 | 0.0303 | <0.0001 | 0.4448 | 0.0404 | * | ns | * |
| ZT5-6 | <0.0001 | 0.1431 | 0.0077 | <0.0001 | 0.8487 | <0.0001 | **** | ns | ** |
| ZT6-7 | <0.0001 | 0.0808 | <0.0001 | <0.0001 | 0.7751 | <0.0001 | **** | ns | **** |
| ZT7-8 | 0.9826 | 0.6338 | <0.0001 | 0.0002 | >0.9999 | <0.0001 | ns | ns | **** |
| ZT8-9 | 0.1016 | 0.8709 | 0.0020 | 0.0145 | 0.6063 | 0.0553 | ns | ns | ns |
| ZT9-10 | 0.0085 | >0.9999 | 0.6787 | <0.0001 | 0.0324 | <0.0001 | ** | ns | ns |
| ZT10-11 | 0.3917 | 0.5212 | 0.1024 | 0.0981 | 0.5159 | <0.0001 | ns | ns | ns |
| ZT11-12 | 0.1553 | 0.1340 | 0.4362 | 0.3605 | 0.1312 | 0.2042 | ns | ns | ns |
| ZT12-13 | 0.4068 | 0.4866 | 0.7569 | 0.0564 | 0.0213 | 0.0500 | ns | ns | ns |
| ZT13-14 | 0.0387 | 0.0043 | 0.9123 | 0.0605 | 0.7237 | 0.9910 | ns | ns | ns |
| ZT14-15 | <0.0001 | 0.0005 | 0.0007 | 0.0007 | 0.3045 | 0.0307 | *** | ns | * |
| ZT15-16 | 0.0001 | <0.0001 | <0.0001 | <0.0001 | 0.2105 | <0.0001 | *** | ns | **** |
| ZT16-17 | 0.0008 | <0.0001 | <0.0001 | 0.0001 | 0.0009 | <0.0001 | *** | *** | **** |
| ZT17-18 | <0.0001 | 0.0003 | 0.0002 | <0.0001 | 0.0161 | <0.0001 | **** | * | *** |
| ZT18-19 | 0.0011 | <0.0001 | 0.0001 | <0.0001 | <0.0001 | <0.0001 | ** | **** | *** |
| ZT19-20 | 0.0025 | <0.0001 | 0.0017 | <0.0001 | 0.0126 | 0.0001 | ** | * | ** |
| ZT20-21 | <0.0001 | <0.0001 | <0.0001 | <0.0001 | <0.0001 | <0.0001 | **** | **** | **** |
| ZT21-22 | 0.0009 | 0.0002 | <0.0001 | <0.0001 | <0.0001 | <0.0001 | *** | *** | **** |
| ZT22-23 | 0.0019 | <0.0001 | 0.0002 | 0.0066 | 0.0006 | 0.0003 | ** | *** | *** |
| ZT23-24 | 0.0002 | <0.0001 | 0.0003 | <0.0001 | 0.0089 | 0.0002 | *** | ** | *** |
| ZT24-25 | 0.9226 | 0.6206 | 0.1322 | 0.1509 | 0.1250 | 0.0025 | ns | ns | ns |
| ZT25-26 | 0.4159 | 0.9544 | 0.8900 | 0.0498 | 0.0208 | 0.0055 | ns | ns | ns |
| ZT26-27 | 0.6479 | 0.9408 | 0.6301 | 0.8007 | 0.5577 | 0.1041 | ns | ns | ns |
| ZT27-28 | 0.1835 | 0.5260 | 0.2652 | 0.2042 | 0.4179 | 0.9925 | ns | ns | ns |
| ZT28-29 | 0.0001 | 0.8840 | 0.0050 | 0.1684 | 0.9925 | 0.5242 | ns | ns | ns |
| ZT29-30 | 0.0170 | 0.9084 | 0.0048 | <0.0001 | 0.6428 | 0.0360 | ns | ns | * |
| ZT30-31 | 0.0221 | 0.9918 | 0.0159 | <0.0001 | 0.0201 | <0.0001 | ns | ns | ns |

| | | | Group | | Time | | Interaction | |
|---|---|---|---|---|---|---|---|---|
| | Data | DFn, DFd | F | p | F | p | F | p |
| | Total sleep | 3, 149 | 10.14 | <0.0001 | 129.9 | <0.0001 | 11.29 | <0.0001 |
| Fig. 9A | Sleep episodes | 3, 149 | 26.05 | <0.0001 | 4.538 | <0.0001 | 3.516 | 0.0002 |
| | P(wake) | 3, 149 | 64.03 | <0.0001 | 72.10 | <0.0001 | 9.350 | <0.0001 |

Dunnett's multiple comparisons

| | VT064246, MB299B/$shi^{ts1}$ vs $shi^{ts1}$/+ | | | MB299B/$shi^{ts1}$ vs $shi^{ts1}$/+ | | | VT064246/$shi^{ts1}$ vs $shi^{ts1}$/+ | | | Summary | | |
|---|---|---|---|---|---|---|---|---|---|---|---|---|
| | Total sleep | Sleep episodes | P(wake) | Total sleep | Sleep episodes | P(wake) | Total sleep | Sleep episodes | P(wake) | Total sleep | Sleep episodes | P(wake) |
| ZT0-1 | 0.0012 | 0.0822 | 0.6939 | 0.4009 | 0.9999 | 0.7227 | 0.9699 | 0.9982 | 0.1135 | ns | ns | ns |
| ZT1-2 | 0.6433 | >0.9999 | 0.9528 | 0.9985 | 0.8127 | 0.9989 | 0.7121 | 0.9997 | 0.9997 | ns | ns | ns |
| ZT2-3 | 0.0585 | 0.7961 | 0.0255 | 0.3372 | 0.8271 | 0.8464 | 0.0792 | 0.4275 | 0.1163 | ns | ns | ns |
| ZT3-4 | 0.4668 | 0.1137 | >0.9999 | 0.5251 | 0.9315 | 0.9486 | 0.0362 | 0.9550 | 0.4602 | ns | ns | ns |
| ZT4-5 | <0.0001 | 0.0433 | <0.0001 | <0.0001 | 0.9898 | <0.0001 | <0.0001 | 0.9008 | <0.0001 | **** | ns | **** |
| ZT5-6 | 0.0962 | 0.9763 | <0.0001 | 0.9799 | 0.0140 | 0.6201 | 0.0334 | >0.9999 | 0.0002 | ns | ns | ns |
| ZT6-7 | 0.0001 | 0.4568 | <0.0001 | 0.7972 | 0.4453 | 0.5538 | 0.0206 | 0.2832 | 0.0031 | ns | ns | ns |
| ZT7-8 | <0.0001 | <0.0001 | <0.0001 | 0.004 | 0.1856 | 0.2448 | <0.0001 | 0.0015 | 0.0009 | ** | ns | ns |
| ZT8-9 | <0.0001 | 0.3101 | <0.0001 | <0.0001 | 0.9990 | <0.0001 | <0.0001 | 0.5269 | <0.0001 | **** | ns | **** |
| ZT9-10 | 0.8679 | 0.0001 | 0.0824 | 0.1871 | 0.0037 | 0.9995 | 0.5547 | 0.0090 | 0.9404 | ns | ** | ns |
| ZT10-11 | 0.0002 | <0.0001 | <0.0001 | 0.0005 | <0.0001 | <0.0001 | 0.0079 | 0.0034 | <0.0001 | ** | ** | **** |
| ZT11-12 | 0.0008 | <0.0001 | <0.0001 | 0.0026 | 0.0006 | <0.0001 | 0.0400 | 0.0169 | 0.0022 | * | * | ** |
| ZT12-13 | 0.0004 | 0.9910 | <0.0001 | <0.0001 | 0.7508 | <0.0001 | <0.0001 | 0.4758 | <0.0001 | *** | ns | **** |
| ZT13-14 | <0.0001 | 0.0002 | <0.0001 | <0.0001 | 0.0011 | <0.0001 | <0.0001 | 0.0021 | <0.0001 | **** | ** | **** |
| ZT14-15 | <0.0001 | <0.0001 | 0.0139 | <0.0001 | <0.0001 | 0.0003 | <0.0001 | 0.0046 | <0.0001 | **** | ** | * |
| ZT15-16 | 0.0002 | <0.0001 | 0.0066 | 0.0002 | 0.0045 | 0.0011 | <0.0001 | 0.0161 | <0.0001 | *** | * | ** |
| ZT16-17 | <0.0001 | <0.0001 | 0.0459 | 0.0001 | 0.0003 | 0.0153 | <0.0001 | 0.0006 | <0.0001 | *** | *** | * |
| ZT17-18 | 0.0022 | 0.0003 | 0.0495 | 0.0018 | 0.0974 | 0.019 | <0.0001 | 0.2057 | <0.0001 | ** | ns | * |
| ZT18-19 | 0.0033 | <0.0001 | 0.0241 | 0.0020 | <0.0001 | 0.0847 | <0.0001 | 0.0001 | <0.0001 | ** | *** | ns |
| ZT19-20 | <0.0001 | <0.0001 | 0.0859 | 0.0019 | 0.0003 | 0.4053 | <0.0001 | <0.0001 | <0.0001 | ** | *** | ns |
| ZT20-21 | 0.0016 | <0.0001 | 0.1985 | 0.0844 | 0.0276 | 0.3787 | <0.0001 | <0.0001 | 0.0082 | ns | * | ns |
| ZT21-22 | 0.0106 | 0.0003 | 0.6366 | 0.0897 | 0.0251 | 0.4458 | <0.0001 | <0.0001 | 0.0116 | ns | * | ns |
| ZT22-23 | 0.4068 | 0.0099 | 0.8556 | 0.1296 | 0.1362 | 0.3387 | 0.0011 | 0.0081 | 0.0084 | ns | ns | ns |
| ZT23-24 | 0.9757 | 0.0020 | <0.0001 | 0.2522 | 0.0193 | <0.0001 | 0.0263 | 0.0218 | <0.0001 | ns | * | **** |
| ZT24-25 | <0.0001 | <0.0001 | <0.0001 | 0.1589 | 0.0046 | <0.0001 | 0.0047 | 0.0138 | <0.0001 | ns | * | **** |
| ZT25-26 | <0.0001 | <0.0001 | <0.0001 | 0.0778 | 0.0008 | <0.0001 | 0.0495 | 0.8653 | <0.0001 | ns | ns | **** |
| ZT26-27 | <0.0001 | 0.0059 | <0.0001 | 0.0807 | 0.0037 | <0.0001 | 0.0115 | 0.4180 | 0.0002 | ns | ns | *** |
| ZT27-28 | <0.0001 | 0.0230 | <0.0001 | 0.0629 | 0.0066 | 0.535 | 0.0111 | 0.6860 | 0.6026 | ns | ns | ns |
| ZT28-29 | 0.0003 | 0.0188 | 0.011 | 0.9546 | 0.0928 | 0.724 | 0.8995 | 0.7298 | >0.9999 | ns | ns | ns |

| | | | Group | | Time | | Interaction | |
|---|---|---|---|---|---|---|---|---|
| | Data | DFn, DFd | F | p | F | p | F | p |
| Fig. 9B | Total sleep | 3, 116 | 1.191 | 0.3162 | 362.6 | ＜0.0001 | 5.321 | ＜0.0001 |
| | Sleep episodes | 3, 116 | 5.816 | 0.0010 | 24.25 | ＜0.0001 | 2.527 | ＜0.0001 |
| | P(wake) | 3, 116 | 13.07 | ＜0.0001 | 191.8 | ＜0.0001 | 5.645 | ＜0.0001 |

Dunnett's multiple comparisons

| | VT064246, MB299B/$shi^{ts1}$ vs $shi^{ts1}$/+ | | | MB299B/$shi^{ts1}$ vs $shi^{ts1}$/+ | | | VT064246/$shi^{ts1}$ vs $shi^{ts1}$/+ | | | Summary | | |
|---|---|---|---|---|---|---|---|---|---|---|---|---|
| | Total sleep | Sleep episodes | P(wake) | Total sleep | Sleep episodes | P(wake) | Total sleep | Sleep episodes | P(wake) | Total sleep | Sleep episodes | P(wake) |
| ZT0-1 | 0.0103 | 0.0017 | <0.0001 | 0.0082 | 0.0043 | <0.0001 | 0.0522 | 0.0556 | 0.0009 | ns | ns | *** |
| ZT1-2 | 0.0118 | 0.0007 | <0.0001 | 0.0037 | 0.0006 | <0.0001 | 0.3924 | 0.3516 | 0.1495 | ns | ns | ns |
| ZT2-3 | 0.2496 | 0.9481 | 0.4385 | 0.9134 | 0.9421 | 0.7102 | 0.0055 | 0.0327 | <0.0001 | ns | ns | ns |
| ZT3-4 | 0.1313 | 0.9997 | 0.0053 | 0.9615 | 0.6141 | 0.3147 | 0.0005 | 0.8393 | <0.0001 | ns | ns | ns |
| ZT4-5 | <0.0001 | 0.7066 | <0.0001 | <0.0001 | 0.5016 | <0.0001 | <0.0001 | 0.8397 | <0.0001 | **** | ns | **** |
| ZT5-6 | 0.5137 | 0.5190 | 0.2726 | 0.0048 | 0.0576 | 0.9944 | 0.5146 | >0.9999 | 0.0112 | ns | ns | ns |
| ZT6-7 | 0.2615 | 0.2026 | 0.7645 | 0.1905 | 0.0009 | 0.6368 | 0.6394 | 0.1354 | 0.5593 | ns | ns | ns |
| ZT7-8 | 0.0123 | 0.0077 | 0.1112 | 0.1023 | 0.0581 | 0.1758 | 0.4824 | 0.9989 | 0.5433 | ns | ns | ns |
| ZT8-9 | 0.7349 | 0.0093 | 0.8624 | >0.9999 | 0.6661 | 0.963 | 0.1297 | 0.0084 | 0.1596 | ns | ns | ns |
| ZT9-10 | 0.3777 | 0.1718 | 0.0706 | 0.4230 | 0.1025 | 0.1385 | 0.0136 | 0.0331 | 0.0006 | ns | ns | ns |
| ZT10-11 | 0.1639 | 0.0055 | 0.2396 | 0.4143 | 0.0769 | 0.3056 | 0.1275 | 0.0316 | 0.0731 | ns | ns | ns |
| ZT11-12 | 0.0035 | 0.0013 | <0.0001 | 0.0051 | 0.0085 | <0.0001 | 0.1701 | 0.1295 | 0.0001 | ns | ns | *** |
| ZT12-13 | 0.9002 | 0.9165 | 0.8841 | 0.2025 | 0.1386 | 0.0027 | 0.9968 | 0.1032 | 0.6822 | ns | ns | ns |
| ZT13-14 | 0.3590 | 0.0660 | 0.4135 | 0.0027 | 0.0305 | 0.0062 | 0.0162 | 0.0035 | 0.1486 | ns | ns | ns |
| ZT14-15 | 0.3260 | 0.1938 | 0.7823 | 0.0125 | 0.0062 | 0.437 | 0.2132 | 0.1447 | 0.7811 | ns | ns | ns |
| ZT15-16 | 0.0681 | 0.1197 | 0.6239 | 0.0057 | 0.0677 | 0.5947 | 0.0302 | 0.0703 | 0.8581 | ns | ns | ns |
| ZT16-17 | 0.3349 | 0.8176 | 0.9227 | 0.1069 | 0.0465 | 0.763 | 0.4191 | 0.1614 | 0.9277 | ns | ns | ns |
| ZT17-18 | 0.7686 | 0.8747 | 0.9730 | 0.3623 | 0.0955 | 0.8591 | 0.7458 | 0.4303 | 0.9661 | ns | ns | ns |
| ZT18-19 | 0.2418 | 0.3409 | 0.8494 | 0.0706 | 0.0501 | 0.9412 | 0.1038 | 0.3003 | 0.6700 | ns | ns | ns |
| ZT19-20 | 0.1315 | 0.9081 | 0.8608 | 0.1567 | 0.6220 | 0.9717 | 0.1203 | 0.9979 | 0.9467 | ns | ns | ns |
| ZT20-21 | 0.2492 | 0.7762 | 0.9562 | 0.9919 | 0.3888 | >0.9999 | 0.6921 | 0.9825 | >0.9999 | ns | ns | ns |
| ZT21-22 | 0.4072 | 0.9972 | 0.7979 | 0.8373 | 0.9045 | >0.9999 | 0.9271 | 0.7844 | 0.9960 | ns | ns | ns |
| ZT22-23 | 0.9986 | 0.5470 | 0.9999 | 0.8669 | 0.9988 | 0.9958 | 0.8884 | 0.9867 | 0.9966 | ns | ns | ns |
| ZT23-24 | <0.0001 | 0.0252 | 0.3332 | 0.0073 | 0.9617 | 0.6343 | <0.0001 | 0.9089 | 0.4822 | ** | ns | ns |
| ZT24-25 | 0.0004 | <0.0001 | <0.0001 | 0.0002 | 0.0006 | <0.0001 | 0.0147 | 0.0716 | 0.0035 | * | ns | ** |
| ZT25-26 | 0.0239 | 0.0014 | <0.0001 | 0.0025 | <0.0001 | <0.0001 | 0.6768 | 0.0498 | 0.1390 | ns | * | ns |

| | | | | | | | | | | | | |
|---|---|---|---|---|---|---|---|---|---|---|---|---|
| ZT26-27 | 0.9932 | 0.1633 | 0.7489 | 0.2377 | 0.0081 | 0.2974 | 0.3914 | 0.7054 | 0.0420 | ns | ns | ns |
| ZT27-28 | 0.0007 | 0.9996 | 0.0018 | 0.3792 | 0.3802 | 0.3490 | 0.0005 | 0.5244 | <0.0001 | ns | ns | ns |
| ZT28-29 | 0.0041 | 0.8965 | 0.0254 | 0.0534 | 0.4718 | 0.2205 | <0.0001 | 0.9933 | <0.0001 | ns | ns | ns |

| | | | Group | | Time | | Interaction | |
|---|---|---|---|---|---|---|---|---|
| | Data | DFn, DFd | F | p | F | p | F | p |
| | Total sleep | 2, 121 | 5.4550 | 0.0054 | 163.3 | ＜0.0001 | 4.773 | ＜0.0001 |
| Fig. 9C | Sleep episodes | 2, 121 | 46.12 | ＜0.0001 | 53.63 | ＜0.0001 | 2.890 | ＜0.0001 |
| | P(wake) | 2, 121 | 28.78 | ＜0.0001 | 111.8 | ＜0.0001 | 4.606 | ＜0.0001 |

THIP — Dunnett's multiple comparisons

| | MB299B/dTrpA1 vs MB299B/+ | | | MB299B/dTrpA1 vs dTrpA1/+ | | | Summary | | |
|---|---|---|---|---|---|---|---|---|---|
| | Total sleep | Sleep episodes | P(wake) | Total sleep | Sleep episodes | P(wake) | Total sleep | Sleep episodes | P(wake) |
| ZT0-1 | >0.9999 | >0.9999 | >0.9999 | >0.9999 | >0.9999 | >0.9999 | ns | ns | ns |
| ZT1-2 | <0.0001 | 0.0007 | <0.0001 | 0.0015 | 0.0015 | <0.0001 | ** | ** | **** |
| ZT2-3 | 0.0025 | 0.0020 | 0.0002 | 0.9165 | 0.0001 | 0.0018 | ns | ** | ** |
| ZT3-4 | 0.4794 | 0.0039 | 0.4226 | 0.2734 | <0.0001 | 0.0002 | ns | ** | ns |
| ZT4-5 | 0.1300 | 0.8895 | 0.1198 | 0.9998 | 0.0203 | 0.3675 | ns | ns | ns |
| ZT5-6 | 0.0066 | 0.0002 | 0.0003 | 0.9625 | <0.0001 | 0.8478 | ns | *** | ns |
| ZT6-7 | 0.1271 | <0.0001 | 0.1878 | 0.9319 | 0.0007 | 0.5117 | ns | *** | ns |
| ZT7-8 | 0.0053 | 0.0196 | 0.0102 | 0.1474 | 0.0003 | 0.1118 | ns | * | ns |
| ZT8-9 | 0.8029 | 0.2057 | 0.6601 | 0.0047 | <0.0001 | 0.0051 | ns | ns | ns |
| ZT9-10 | 0.1210 | <0.0001 | 0.1673 | 0.7838 | <0.0001 | 0.0794 | ns | **** | ns |
| ZT10-11 | 0.2984 | 0.4497 | 0.8538 | 0.1240 | 0.0025 | 0.8052 | ns | ns | ns |
| ZT11-12 | 0.1363 | 0.9995 | 0.986 | 0.0353 | 0.7733 | 0.9802 | ns | ns | ns |
| ZT12-13 | 0.8437 | 0.0020 | 0.9424 | 0.0054 | 0.1270 | 0.0187 | ns | ns | ns |
| ZT13-14 | 0.9803 | 0.9990 | 0.9752 | 0.2276 | 0.6382 | 0.8253 | ns | ns | ns |
| ZT14-15 | 0.2842 | 0.6901 | 0.928 | 0.5218 | 0.9982 | 0.9938 | ns | ns | ns |
| ZT15-16 | 0.1458 | 0.5454 | 0.8994 | 0.0301 | 0.1401 | 0.6789 | ns | ns | ns |
| ZT16-17 | 0.9067 | 0.9110 | 0.9992 | 0.7096 | 0.5752 | 0.8705 | ns | ns | ns |
| ZT17-18 | 0.8000 | 0.5803 | 0.9404 | 0.9212 | >0.9999 | 0.9924 | ns | ns | ns |
| ZT18-19 | 0.9836 | 0.9874 | 0.9978 | 0.5325 | 0.9044 | 0.9406 | ns | ns | ns |
| ZT19-20 | 0.7899 | 0.9843 | 0.9936 | 0.6374 | 0.9273 | 0.9577 | ns | ns | ns |
| ZT20-21 | 0.0596 | 0.5384 | 0.9297 | 0.1306 | 0.4757 | 0.6704 | ns | ns | ns |
| ZT21-22 | 0.3002 | 0.8908 | 0.8979 | 0.2139 | 0.7441 | 0.8780 | ns | ns | ns |
| ZT22-23 | 0.9757 | 0.9758 | 0.9983 | 0.8121 | 0.9554 | 0.9193 | ns | ns | ns |
| ZT23-24 | 0.1441 | 0.6972 | 0.8504 | 0.1902 | 0.9993 | 0.5056 | ns | ns | ns |
| ZT24-25 | <0.0001 | 0.9833 | <0.0001 | 0.0002 | 0.8403 | <0.0001 | *** | ns | **** |
| ZT25-26 | 0.1190 | 0.7561 | 0.0314 | 0.0024 | 0.9860 | <0.0001 | ns | ns | * |
| ZT26-27 | 0.9528 | 0.5715 | 0.5897 | 0.0185 | 0.3839 | <0.0001 | ns | ns | ns |
| ZT27-28 | 0.1630 | 0.0010 | 0.0131 | 0.0295 | 0.0009 | <0.0001 | ns | *** | * |
| ZT28-29 | 0.2208 | 0.7238 | 0.2136 | 0.5167 | 0.1863 | 0.2903 | ns | ns | ns |
| ZT29-30 | 0.9525 | 0.2923 | 0.8716 | 0.0403 | 0.0289 | 0.0206 | ns | ns | ns |
| ZT30-31 | 0.7100 | 0.8909 | 0.8372 | 0.0642 | 0.0258 | 0.0041 | ns | ns | ns |

| | | | Group | | Time | | Interaction | |
|---|---|---|---|---|---|---|---|---|
| | Data | DFn, DFd | F | p | F | p | F | p |
| | Total sleep | 2, 120 | 11.28 | ＜0.0001 | 187.6 | ＜0.0001 | 4.921 | ＜0.0001 |
| Fig. 9D | Sleep episodes | 2, 120 | 136.6 | ＜0.0001 | 24.47 | ＜0.0001 | 2.663 | ＜0.0001 |
| | P(wake) | 2, 120 | 68.03 | ＜0.0001 | 115.5 | ＜0.0001 | 3.822 | ＜0.0001 |

No THIP — Dunnett's multiple comparisons

| | MB299B/dTrpA1 vs MB299B/+ | | | MB299B/dTrpA1 vs dTrpA1/+ | | | Summary | | |
|---|---|---|---|---|---|---|---|---|---|
| | Total sleep | Sleep episodes | P(wake) | Total sleep | Sleep episodes | P(wake) | Total sleep | Sleep episodes | P(wake) |
| ZT0-1 | 0.0000 | >0.9999 | >0.9999 | 0.0000 | >0.9999 | >0.9999 | ns | ns | ns |
| ZT1-2 | <0.0001 | <0.0001 | <0.0001 | 0.3155 | 0.0098 | 0.1434 | ns | ** | ns |
| ZT2-3 | 0.3188 | 0.0621 | 0.1293 | 0.9686 | 0.0345 | 0.2843 | ns | ns | ns |
| ZT3-4 | 0.4801 | 0.1031 | 0.499 | 0.5186 | 0.1391 | >0.9999 | ns | ns | ns |
| ZT4-5 | 0.0058 | 0.0044 | 0.0177 | 0.3284 | 0.0375 | 0.9048 | ns | * | ns |
| ZT5-6 | 0.0052 | 0.0004 | 0.0120 | 0.2461 | 0.0003 | 0.3825 | ns | *** | ns |
| ZT6-7 | 0.0349 | 0.1035 | 0.0179 | 0.5165 | 0.8308 | 0.3524 | ns | ns | ns |
| ZT7-8 | 0.3903 | 0.5888 | 0.5023 | <0.0001 | 0.1743 | <0.0001 | ns | ns | ns |
| ZT8-9 | 0.0003 | 0.0939 | 0.0016 | <0.0001 | <0.0001 | <0.0001 | *** | ns | ** |
| ZT9-10 | 0.4517 | 0.6421 | 0.6328 | 0.1320 | 0.1189 | 0.1644 | ns | ns | ns |
| ZT10-11 | 0.5145 | 0.2046 | 0.9002 | 0.9614 | 0.2560 | 0.7398 | ns | ns | ns |
| ZT11-12 | 0.8946 | 0.8603 | 0.7208 | 0.7045 | 0.7044 | 0.5085 | ns | ns | ns |
| ZT12-13 | 0.3118 | 0.3101 | 0.0777 | 0.2076 | 0.8004 | 0.0322 | ns | ns | ns |
| ZT13-14 | 0.0044 | 0.0002 | 0.0090 | 0.0002 | 0.0005 | 0.0019 | ** | *** | ** |
| ZT14-15 | 0.0070 | 0.0154 | 0.0187 | 0.1159 | 0.4901 | 0.0699 | ns | ns | ns |

| | | | | | | | | | |
|---|---|---|---|---|---|---|---|---|---|
| ZT15-16 | 0.0894 | 0.1660 | 0.2314 | 0.0439 | 0.6076 | 0.1956 | ns | ns | ns |
| ZT16-17 | 0.2326 | 0.4380 | 0.5854 | 0.1302 | 0.8774 | 0.5638 | ns | ns | ns |
| ZT17-18 | 0.2170 | 0.7168 | 0.6225 | 0.2571 | 0.8930 | 0.6692 | ns | ns | ns |
| ZT18-19 | 0.3937 | >0.9999 | 0.8572 | 0.8981 | 0.9924 | 0.9642 | ns | ns | ns |
| ZT19-20 | 0.2576 | 0.6006 | 0.6305 | 0.6617 | 0.9979 | 0.8228 | ns | ns | ns |
| ZT20-21 | 0.0276 | 0.8190 | 0.2093 | 0.3351 | 0.7864 | 0.5658 | ns | ns | ns |
| ZT21-22 | 0.3564 | 0.9404 | 0.7808 | 0.9997 | 0.8333 | 0.9989 | ns | ns | ns |
| ZT22-23 | 0.9545 | 0.7168 | 0.9888 | 0.3941 | 0.4378 | 0.7727 | ns | ns | ns |
| ZT23-24 | 0.3011 | 0.4537 | 0.8086 | 0.6479 | 0.5679 | 0.9384 | ns | ns | ns |
| ZT24-25 | 0.6425 | 0.7526 | 0.0360 | 0.1782 | 0.5018 | 0.0042 | ns | ns | * |
| ZT25-26 | 0.0145 | 0.9453 | 0.0014 | 0.0987 | 0.0664 | 0.0074 | ns | ns | ** |
| ZT26-27 | 0.0069 | 0.6050 | <0.0001 | 0.4597 | 0.0752 | 0.0924 | ns | ns | ns |
| ZT27-28 | 0.0110 | 0.1354 | 0.0008 | 0.1130 | 0.0263 | 0.0120 | ns | ns | * |
| ZT28-29 | 0.0002 | 0.8988 | <0.0001 | 0.0033 | 0.0454 | <0.0001 | ** | ns | **** |
| ZT29-30 | <0.0001 | 0.5663 | <0.0001 | 0.0004 | 0.0084 | <0.0001 | *** | ns | **** |
| ZT30-31 | <0.0001 | 0.0666 | <0.0001 | <0.0001 | 0.0007 | <0.0001 | **** | ns | **** |

| | | | Group | | Time | | Interaction | |
|---|---|---|---|---|---|---|---|---|
| | Data | DFn, DFd | F | p | F | p | F | p |
| Fig. 11A | Total sleep | 2, 86 | 28.77 | ＜0.0001 | 75.90 | ＜0.0001 | 6.104 | ＜0.0001 |
| | Sleep episodes | 2, 86 | 13.09 | ＜0.0001 | 5.620 | ＜0.0001 | 3.299 | ＜0.0001 |
| | P(wake) | 2, 86 | 33.61 | ＜0.0001 | 61.77 | ＜0.0001 | 5.769 | ＜0.0001 |

Dunnett's multiple comparisons

| | MB310C/dTrpA1 vs MB310C/+ | | | MB310C/dTrpA1 vs dTrpA1/+ | | | Summary | | |
|---|---|---|---|---|---|---|---|---|---|
| | Total sleep | Sleep episodes | P(wake) | Total sleep | Sleep episodes | P(wake) | Total sleep | Sleep episodes | P(wake) |
| ZT0-1 | 0.9209 | 0.8212 | 0.9727 | >0.9999 | 0.9125 | 0.3328 | ns | ns | ns |
| ZT1-2 | 0.9739 | 0.9054 | 0.8915 | 0.7835 | 0.3355 | 0.6966 | ns | ns | ns |
| ZT2-3 | 0.9282 | 0.8102 | 0.9535 | 0.7731 | 0.2023 | 0.5547 | ns | ns | ns |
| ZT3-4 | 0.9991 | 0.7525 | 0.9885 | 0.9569 | 0.5942 | 0.9750 | ns | ns | ns |
| ZT4-5 | 0.9772 | 0.2479 | 0.8653 | 0.9943 | 0.7517 | 0.9871 | ns | ns | ns |
| ZT5-6 | 0.2201 | 0.1629 | 0.0612 | 0.0243 | 0.0002 | 0.0073 | ns | ns | ns |
| ZT6-7 | 0.8267 | 0.1489 | 0.9766 | 0.7558 | 0.951 | 0.843 | ns | ns | ns |
| ZT7-8 | 0.9615 | 0.8731 | 0.9956 | 0.0278 | 0.0037 | 0.0045 | ns | ns | ns |
| ZT8-9 | 0.9966 | 0.7818 | 0.2757 | 0.0719 | <0.0001 | 0.0007 | ns | ns | ns |
| ZT9-10 | 0.9700 | 0.8157 | 0.7109 | 0.0825 | 0.0001 | 0.0036 | ns | ns | ns |
| ZT10-11 | 0.9608 | 0.7226 | 0.5655 | 0.6760 | 0.2394 | 0.1232 | ns | ns | ns |
| ZT11-12 | >0.9999 | 0.9985 | 0.9268 | 0.6941 | 0.1747 | 0.1854 | ns | ns | ns |
| ZT12-13 | 0.4999 | 0.4009 | 0.0879 | 0.7045 | 0.5522 | 0.7301 | ns | ns | ns |
| ZT13-14 | 0.0144 | 0.9545 | 0.0398 | 0.2463 | 0.2266 | 0.2253 | ns | ns | ns |
| ZT14-15 | 0.2824 | 0.9829 | 0.3489 | 0.1192 | 0.2082 | 0.1915 | ns | ns | ns |
| ZT15-16 | 0.0708 | 0.4962 | 0.2753 | 0.1845 | 0.3273 | 0.4015 | ns | ns | ns |
| ZT16-17 | 0.0443 | 0.3957 | 0.1712 | 0.3006 | 0.1052 | 0.5685 | ns | ns | ns |
| ZT17-18 | 0.2605 | 0.2765 | 0.5275 | 0.1293 | 0.3194 | 0.3575 | ns | ns | ns |
| ZT18-19 | 0.0608 | 0.4502 | 0.4503 | 0.1545 | 0.0556 | 0.2011 | ns | ns | ns |
| ZT19-20 | 0.1506 | 0.5139 | 0.7401 | 0.0511 | 0.0082 | 0.1124 | ns | ns | ns |
| ZT20-21 | 0.2076 | 0.4447 | 0.3850 | 0.1263 | 0.0309 | 0.4961 | ns | ns | ns |
| ZT21-22 | 0.4032 | 0.5139 | 0.4121 | 0.1451 | 0.2461 | 0.5181 | ns | ns | ns |
| ZT22-23 | 0.2872 | 0.8730 | 0.4268 | 0.9456 | 0.0178 | 0.8179 | ns | ns | ns |
| ZT23-24 | 0.4082 | 0.8873 | 0.6733 | 0.5687 | 0.0393 | 0.8829 | ns | ns | ns |
| ZT24-25 | 0.5076 | 0.8320 | 0.9224 | 0.9954 | 0.7415 | 0.7698 | ns | ns | ns |
| ZT25-26 | 0.324 | 0.1284 | 0.4005 | 0.4397 | 0.5522 | 0.4096 | ns | ns | ns |
| ZT26-27 | 0.1700 | 0.3164 | 0.3728 | 0.4213 | 0.9889 | 0.5227 | ns | ns | ns |
| ZT27-28 | 0.1106 | 0.1572 | 0.1624 | 0.6258 | 0.5215 | 0.6858 | ns | ns | ns |
| ZT28-29 | 0.0240 | 0.3211 | 0.0181 | 0.4736 | 0.5011 | 0.2201 | ns | ns | ns |
| ZT29-30 | 0.0164 | 0.2984 | 0.0353 | 0.4617 | 0.9510 | 0.2879 | ns | ns | ns |
| ZT30-31 | 0.0032 | 0.9220 | 0.0962 | 0.4974 | 0.7618 | 0.1488 | ns | ns | ns |

| | | | Group | | Time | | Interaction | |
|---|---|---|---|---|---|---|---|---|
| | Data | DFn, DFd | F | p | F | p | F | p |
| Fig. 11B | Total sleep | 2, 120 | 30.75 | ＜0.0001 | 75.57 | ＜0.0001 | 8.833 | ＜0.0001 |
| | Sleep episodes | 2, 120 | 34.41 | ＜0.0001 | 8.902 | ＜0.0001 | 2.992 | ＜0.0001 |
| | P(wake) | 2, 120 | 47.69 | ＜0.0001 | 73.51 | ＜0.0001 | 9.187 | ＜0.0001 |

Dunnett's multiple comparisons

| | MB310C/$shi^{ts1}$ vs MB310C/+ | | | MB310C/$shi^{ts1}$ vs $shi^{ts1}$/+ | | | Summary | | |
|---|---|---|---|---|---|---|---|---|---|
| | Total sleep | Sleep episodes | P(wake) | Total sleep | Sleep episodes | P(wake) | Total sleep | Sleep episodes | P(wake) |
| ZT0-1 | 0.8249 | 0.9125 | 0.6284 | 0.6978 | 0.8573 | 0.1424 | ns | ns | ns |
| ZT1-2 | 0.9926 | 0.9731 | 0.9487 | 0.8521 | 0.7287 | 0.9031 | ns | ns | ns |
| ZT2-3 | 0.9967 | 0.1758 | 0.8049 | 0.9925 | 0.7195 | 0.3323 | ns | * | ns |

| | | | | | | | | | |
|---|---|---|---|---|---|---|---|---|---|
| ZT3-4 | 0.6008 | 0.1189 | 0.2749 | 0.7758 | 0.2411 | 0.963 | ns | ns | ns |
| ZT4-5 | 0.7972 | 0.0041 | 0.4624 | 0.1953 | 0.2762 | 0.1483 | ns | ns | ns |
| ZT5-6 | 0.5537 | 0.4074 | 0.253 | 0.6152 | 0.0177 | 0.6434 | ns | ns | ns |
| ZT6-7 | 0.0018 | 0.8219 | 0.0022 | 0.7723 | 0.2663 | 0.6300 | ns | ns | ns |
| ZT7-8 | 0.7803 | 0.8340 | 0.9477 | 0.7120 | 0.9266 | 0.8216 | ns | ns | ns |
| ZT8-9 | 0.2172 | 0.0012 | 0.1801 | <0.0001 | 0.0374 | <0.0001 | ns | * | ns |
| ZT9-10 | 0.0408 | 0.0011 | <0.0001 | <0.0001 | 0.0309 | <0.0001 | * | * | **** |
| ZT10-11 | 0.3039 | 0.0038 | 0.0014 | 0.0017 | 0.0109 | <0.0001 | ns | * | ** |
| ZT11-12 | 0.6036 | 0.0226 | 0.0022 | 0.7658 | 0.6119 | 0.7890 | ns | ns | ns |
| ZT12-13 | <0.0001 | 0.0005 | <0.0001 | 0.0001 | 0.2813 | <0.0001 | *** | ns | **** |
| ZT13-14 | <0.0001 | 0.7580 | <0.0001 | <0.0001 | 0.0634 | <0.0001 | **** | ns | **** |
| ZT14-15 | <0.0001 | 0.0089 | <0.0001 | <0.0001 | 0.1252 | <0.0001 | **** | ns | **** |
| ZT15-16 | <0.0001 | 0.0214 | <0.0001 | <0.0001 | 0.0002 | <0.0001 | **** | * | **** |
| ZT16-17 | <0.0001 | 0.0136 | <0.0001 | <0.0001 | 0.0093 | <0.0001 | **** | * | **** |
| ZT17-18 | <0.0001 | 0.1862 | <0.0001 | <0.0001 | 0.1265 | <0.0001 | **** | ns | **** |
| ZT18-19 | <0.0001 | 0.0023 | <0.0001 | <0.0001 | 0.0044 | <0.0001 | **** | ** | **** |
| ZT19-20 | <0.0001 | 0.0015 | <0.0001 | <0.0001 | <0.0001 | <0.0001 | **** | ** | **** |
| ZT20-21 | <0.0001 | 0.0025 | <0.0001 | <0.0001 | <0.0001 | <0.0001 | **** | ** | **** |
| ZT21-22 | 0.0025 | 0.0215 | 0.008 | 0.0009 | <0.0001 | 0.0124 | ** | * | * |
| ZT22-23 | 0.1899 | 0.0152 | 0.2984 | 0.1442 | 0.0027 | 0.3396 | ns | * | ns |
| ZT23-24 | 0.0331 | 0.0110 | 0.0879 | 0.0361 | 0.0026 | 0.2144 | * | * | ns |
| ZT24-25 | 0.9927 | 0.7842 | 0.5996 | 0.8293 | 0.5952 | 0.2000 | ns | ns | ns |
| ZT25-26 | 0.9907 | 0.5504 | 0.8577 | 0.8694 | 0.9369 | 0.6275 | ns | ns | ns |
| ZT26-27 | 0.9713 | 0.1650 | 0.7899 | 0.8366 | 0.7714 | 0.6709 | ns | ns | ns |
| ZT27-28 | 0.9775 | 0.6379 | 0.7312 | 0.6895 | 0.8043 | 0.6911 | ns | ns | ns |
| ZT28-29 | 0.9393 | 0.8604 | 0.9958 | 0.7253 | 0.4614 | 0.6374 | ns | ns | ns |

| | | | Group | | Time | | Interaction | |
|---|---|---|---|---|---|---|---|---|
| | Data | DFn, DFd | F | p | F | p | F | p |
| | Total sleep | 2, 93 | 62.96 | <0.0001 | 227.6 | <0.0001 | 4.132 | <0.0001 |
| Fig. 11C | Sleep episodes | 2, 93 | 32.60 | <0.0001 | 15.14 | <0.0001 | 1.679 | <0.0001 |
| | P(wake) | 2, 93 | 79.54 | <0.0001 | 187.9 | <0.0001 | 4.915 | <0.0001 |

Dunnett's multiple comparisons

| | MB310C/dTrpA1 vs MB310C/+ | | | MB310C/dTrpA1 vs dTrpA1/+ | | | Summary | | |
|---|---|---|---|---|---|---|---|---|---|
| | Total sleep | Sleep episodes | P(wake) | Total sleep | Sleep episodes | P(wake) | Total sleep | Sleep episodes | P(wake) |
| ZT0-1 | 0.3981 | 0.7024 | 0.5048 | 0.0008 | 0.0020 | <0.0001 | ns | ns | ns |
| ZT1-2 | 0.4935 | 0.9603 | 0.0265 | <0.0001 | 0.0030 | <0.0001 | ns | ns | * |
| ZT2-3 | 0.0195 | 0.2692 | 0.0164 | <0.0001 | 0.0473 | <0.0001 | * | ns | * |
| ZT3-4 | 0.1281 | 0.2693 | 0.0536 | 0.9248 | 0.0064 | 0.9872 | ns | ns | ns |
| ZT4-5 | 0.0006 | 0.9603 | 0.0118 | 0.7230 | 0.7803 | 0.9252 | ns | ns | ns |
| ZT5-6 | 0.8092 | 0.3274 | 0.7552 | 0.8155 | 0.7803 | 0.9717 | ns | ns | ns |
| ZT6-7 | 0.8855 | 0.0256 | 0.7206 | 0.5477 | 0.1750 | 0.5827 | ns | ns | ns |
| ZT7-8 | 0.0159 | 0.0631 | <0.0001 | 0.7433 | 0.9133 | 0.9996 | ns | ns | ns |
| ZT8-9 | 0.0212 | 0.0831 | <0.0001 | 0.6106 | 0.8520 | 0.9039 | ns | ns | ns |
| ZT9-10 | 0.1337 | 0.1384 | 0.0005 | 0.5963 | 0.3926 | 0.1859 | ns | ns | ns |
| ZT10-11 | 0.4672 | 0.1079 | 0.0279 | 0.7160 | 0.9603 | 0.4684 | ns | ns | ns |
| ZT11-12 | 0.6810 | 0.6214 | 0.4731 | 0.9636 | >0.9999 | 0.8818 | ns | ns | ns |
| ZT12-13 | 0.2161 | 0.9603 | 0.1588 | 0.6810 | 0.0631 | 0.9781 | ns | ns | ns |
| ZT13-14 | 0.0007 | 0.6214 | 0.0206 | 0.1254 | >0.9999 | 0.2154 | ns | ns | ns |
| ZT14-15 | 0.0141 | 0.2692 | 0.1063 | 0.0009 | 0.0064 | 0.0402 | * | ns | ns |
| ZT15-16 | 0.0305 | 0.4644 | 0.2660 | 0.0133 | 0.4644 | 0.2675 | * | ns | ns |
| ZT16-17 | 0.3516 | 0.9603 | 0.7082 | 0.5339 | 0.9133 | 0.8570 | ns | ns | ns |
| ZT17-18 | 0.5068 | 0.9603 | 0.7770 | 0.9727 | 0.9899 | >0.9999 | ns | ns | ns |
| ZT18-19 | 0.9727 | 0.4644 | 0.9797 | 0.9923 | 0.7024 | 0.9855 | ns | ns | ns |
| ZT19-20 | 0.7365 | 0.8520 | 0.9385 | 0.9667 | 0.9603 | 0.9858 | ns | ns | ns |
| ZT20-21 | 0.4351 | 0.1750 | 0.7562 | 0.6317 | 0.6214 | 0.7603 | ns | ns | ns |
| ZT21-22 | 0.9938 | 0.0631 | 0.9469 | 0.9568 | 0.8520 | 0.9959 | ns | ns | ns |
| ZT22-23 | 0.7020 | 0.5413 | 0.8929 | 0.4164 | >0.9999 | 0.8509 | ns | ns | ns |
| ZT23-24 | 0.8092 | 0.7024 | 0.9290 | 0.9495 | 0.7803 | 0.9978 | ns | ns | ns |
| ZT24-25 | 0.0055 | 0.0043 | <0.0001 | <0.0001 | <0.0001 | <0.0001 | ** | ** | **** |
| ZT25-26 | 0.0005 | 0.7025 | 0.0004 | <0.0001 | 0.0001 | <0.0001 | *** | ns | *** |
| ZT26-27 | 0.6388 | 0.3925 | 0.4132 | <0.0001 | 0.2184 | <0.0001 | ns | ns | ns |
| ZT27-28 | 0.0039 | 0.6214 | 0.0789 | 0.0060 | 0.0256 | 0.009 | ** | ns | ns |
| ZT28-29 | 0.0035 | 0.7022 | 0.0077 | 0.9418 | 0.7025 | 0.9923 | ns | ns | ns |
| ZT29-30 | 0.0003 | 0.2186 | 0.0061 | 0.0297 | 0.6214 | 0.1137 | * | ns | ns |
| ZT30-31 | <0.0001 | >0.9999 | 0.0011 | 0.0028 | 0.9899 | 0.0090 | ** | ns | ** |

| | | | Group | | Time | | Interaction | |
|---|---|---|---|---|---|---|---|---|
| | Data | DFn, DFd | F | p | F | p | F | p |

| | Data | DFn, DFd | F | p | F | p | F | p |
|---|---|---|---|---|---|---|---|---|
| Fig. 11D | Total sleep | 2, 124 | 394.4 | ＜0.0001 | 249.6 | ＜0.0001 | 7.587 | ＜0.0001 |
| | Sleep episodes | 2, 124 | 54.30 | ＜0.0001 | 21.67 | ＜0.0001 | 3.888 | ＜0.0001 |
| | P(wake) | 2, 124 | 315.5 | ＜0.0001 | 246.7 | ＜0.0001 | 9.371 | ＜0.0001 |

Dunnett's multiple comparisons

| | MB310C/$shi^{ts1}$ vs MB310C/+ | | | MB310C/$shi^{ts1}$ vs $shi^{ts1}$/+ | | | Summary | | |
|---|---|---|---|---|---|---|---|---|---|
| | Total sleep | Sleep episodes | P(wake) | Total sleep | Sleep episodes | P(wake) | Total sleep | Sleep episodes | P(wake) |
| ZT0-1 | <0.0001 | 0.1314 | <0.0001 | 0.4594 | 0.3620 | <0.0001 | ns | ns | **** |
| ZT1-2 | <0.0001 | 0.9305 | <0.0001 | 0.6375 | 0.3515 | 0.0138 | ns | ns | * |
| ZT2-3 | <0.0001 | 0.9919 | <0.0001 | 0.1509 | 0.3771 | 0.6089 | ns | ns | ns |
| ZT3-4 | <0.0001 | 0.4696 | <0.0001 | <0.0001 | 0.8699 | <0.0001 | **** | ns | **** |
| ZT4-5 | <0.0001 | 0.0003 | <0.0001 | <0.0001 | 0.0879 | 0.0003 | **** | ns | *** |
| ZT5-6 | 0.0919 | <0.0001 | 0.3379 | 0.0002 | 0.0170 | 0.2468 | ns | * | ns |
| ZT6-7 | 0.1917 | 0.9919 | 0.4836 | 0.1244 | 0.8319 | 0.3470 | ns | ns | ns |
| ZT7-8 | 0.0365 | 0.1050 | 0.2912 | 0.4234 | 0.0889 | 0.7433 | ns | ns | ns |
| ZT8-9 | <0.0001 | 0.0502 | <0.0001 | <0.0001 | 0.0018 | <0.0001 | **** | ns | **** |
| ZT9-10 | <0.0001 | 0.0119 | <0.0001 | <0.0001 | 0.0039 | <0.0001 | **** | * | **** |
| ZT10-11 | <0.0001 | 0.2420 | <0.0001 | 0.0050 | 0.3356 | <0.0001 | ** | ns | **** |
| ZT11-12 | <0.0001 | 0.1995 | <0.0001 | 0.8301 | 0.6909 | 0.4675 | ns | ns | ns |
| ZT12-13 | <0.0001 | 0.6102 | <0.0001 | <0.0001 | 0.2493 | <0.0001 | **** | ns | **** |
| ZT13-14 | <0.0001 | <0.0001 | 0.0002 | <0.0001 | <0.0001 | 0.0001 | **** | **** | *** |
| ZT14-15 | <0.0001 | <0.0001 | 0.0112 | <0.0001 | <0.0001 | 0.002 | **** | **** | * |
| ZT15-16 | 0.0006 | 0.0031 | 0.0577 | 0.0007 | 0.0002 | 0.0524 | *** | ** | ns |
| ZT16-17 | <0.0001 | 0.0062 | 0.0217 | <0.0001 | 0.0053 | 0.0265 | **** | ** | * |
| ZT17-18 | 0.0198 | 0.0086 | 0.2010 | 0.0163 | 0.0039 | 0.1824 | * | ** | ns |
| ZT18-19 | 0.0062 | 0.0007 | 0.0936 | 0.0037 | 0.0011 | 0.0909 | ** | ** | ns |
| ZT19-20 | 0.4957 | 0.2903 | 0.8552 | 0.3081 | 0.0185 | 0.7265 | ns | ns | ns |
| ZT20-21 | 0.3988 | 0.1314 | 0.7413 | 0.3662 | 0.0469 | 0.6987 | ns | ns | ns |
| ZT21-22 | 0.9763 | 0.8202 | 0.9925 | 0.9581 | 0.9219 | 0.9977 | ns | ns | ns |
| ZT22-23 | 0.6599 | 0.3446 | 0.8807 | 0.7810 | 0.9491 | 0.9537 | ns | ns | ns |
| ZT23-24 | 0.9980 | 0.9683 | 0.9986 | >0.9999 | >0.9999 | 0.9980 | ns | ns | ns |
| ZT24-25 | <0.0001 | 0.4046 | <0.0001 | 0.8635 | 0.7294 | 0.0451 | ns | ns | * |
| ZT25-26 | <0.0001 | 0.0022 | <0.0001 | 0.0491 | 0.4102 | 0.3330 | * | ns | ns |
| ZT26-27 | <0.0001 | 0.9683 | <0.0001 | <0.0001 | 0.0576 | 0.0013 | **** | ns | ** |
| ZT27-28 | <0.0001 | 0.0829 | <0.0001 | <0.0001 | 0.8104 | <0.0001 | **** | ns | **** |
| ZT28-29 | <0.0001 | 0.0022 | <0.0001 | <0.0001 | 0.4197 | <0.0001 | **** | ns | **** |

| | | | Group | | Time | | Interaction | |
|---|---|---|---|---|---|---|---|---|
| | Data | DFn, DFd | F | p | F | p | F | p |
| Figure S1A | Total sleep | 2, 122 | 3.663 | 0.0285 | 542.2 | ＜0.0001 | 4.285 | ＜0.0001 |
| | Sleep episodes | 2, 122 | 3.395 | 0.0367 | 25.91 | ＜0.0001 | 2.238 | ＜0.0001 |
| | P(wake) | 2, 109 | 8.903 | 0.0003 | 270.2 | ＜0.0001 | 3.895 | ＜0.0001 |

| | c316/dTrpA1 vs c316/+ | | | c316/dTrpA1 vs dTrpA1/+ | | | Summary | | |
|---|---|---|---|---|---|---|---|---|---|
| | Total sleep | Episodes | P(wake) | Total sleep | Episodes | P(wake) | Total sleep | Episodes | P(wake) |
| ZT0-1 | - | - | 0.2729 | - | - | 0.9731 | - | - | ns |
| ZT1-2 | 0.0416 | 0.0449 | 0.0251 | 0.9944 | 0.7676 | 0.8389 | ns | ns | ns |
| ZT2-3 | 0.0238 | 0.0079 | 0.0661 | 0.6571 | 0.6346 | 0.9494 | ns | ns | ns |
| ZT3-4 | 0.9595 | 0.8212 | 0.7928 | 0.9883 | 0.9244 | 0.7709 | ns | ns | ns |
| ZT4-5 | 0.6517 | 0.2818 | 0.6146 | 0.7396 | 0.9968 | 0.1197 | ns | ns | ns |
| ZT5-6 | 0.4275 | 0.9773 | 0.9887 | 0.0490 | 0.7132 | 0.0131 | ns | ns | ns |
| ZT6-7 | 0.2991 | 0.4860 | 0.7434 | 0.0009 | 0.7980 | 0.9988 | ns | ns | ns |
| ZT7-8 | 0.0009 | 0.6351 | 0.2941 | ＜0.0001 | 0.1021 | 0.0313 | *** | ns | ns |
| ZT8-9 | 0.3708 | 0.8332 | 0.9429 | 0.9635 | 0.0576 | 0.1104 | ns | ns | ns |
| ZT9-10 | 0.9755 | 0.0910 | 0.3739 | 0.0975 | 0.3645 | 0.0268 | ns | ns | ns |
| ZT10-11 | 0.3291 | 0.0958 | 0.0237 | 0.1601 | 0.4534 | 0.1541 | ns | ns | ns |
| ZT11-12 | 0.7080 | 0.3258 | 0.7644 | 0.0099 | 0.0085 | 0.0033 | ns | ns | ns |
| ZT12-13 | 0.6876 | 0.7817 | 0.0611 | 0.0070 | 0.3628 | <0.0001 | ns | ns | ns |
| ZT13-14 | 0.0471 | 0.2383 | 0.264 | ＜0.0001 | 0.0007 | 0.0115 | * | ns | ns |
| ZT14-15 | 0.0525 | 0.1282 | 0.0689 | 0.0241 | 0.0020 | 0.0302 | ns | ns | ns |
| ZT15-16 | 0.1835 | 0.9479 | 0.0906 | 0.1267 | 0.3157 | 0.1077 | ns | ns | ns |
| ZT16-17 | 0.0482 | 0.2292 | 0.2923 | 0.6146 | 0.0307 | 0.3538 | ns | ns | ns |
| ZT17-18 | 0.6288 | 0.8345 | 0.8326 | 0.7372 | 0.7873 | 0.5124 | ns | ns | ns |
| ZT18-19 | 0.7262 | 0.9503 | 0.2319 | 0.8776 | 0.9609 | 0.7318 | ns | ns | ns |
| ZT19-20 | 0.0933 | 0.1098 | 0.5267 | 0.8844 | 0.9567 | 0.5553 | ns | ns | ns |
| ZT20-21 | 0.4664 | 0.7729 | 0.2610 | 0.9652 | ＞0.9999 | 0.3811 | ns | ns | ns |
| ZT21-22 | 0.0684 | 0.2100 | 0.0467 | 0.3909 | 0.0835 | 0.4988 | ns | ns | ns |
| ZT22-23 | 0.1829 | 0.2046 | 0.2136 | 0.1383 | 0.1650 | 0.0132 | ns | ns | ns |
| ZT23-24 | 0.2747 | 0.5565 | 0.4119 | 0.0209 | 0.3919 | 0.0067 | ns | ns | ns |
| ZT24-25 | 0.1774 | 0.0525 | 0.0154 | 0.9825 | 0.8761 | 0.4027 | ns | ns | ns |

| | | | | | | | | | |
|---|---|---|---|---|---|---|---|---|---|
| ZT25-26 | 0.2638 | 0.2641 | 0.0279 | 0.5260 | 0.3123 | 0.5431 | ns | ns | ns |
| ZT26-27 | 0.9986 | 0.9976 | 0.2889 | 0.1335 | 0.1367 | 0.7433 | ns | ns | ns |
| ZT27-28 | 0.9416 | 0.9997 | 0.3050 | 0.8498 | 0.7472 | 0.6701 | ns | ns | ns |
| ZT28-29 | 0.6998 | 0.4309 | 0.0702 | 0.5270 | 0.5210 | 0.5182 | ns | ns | ns |
| ZT29-30 | 0.2991 | 0.2914 | 0.0647 | 0.3147 | 0.8440 | 0.1195 | ns | ns | ns |
| ZT30-31 | 0.3330 | 0.7330 | 0.1081 | 0.0336 | 0.8739 | 0.3067 | ns | ns | ns |

| | Data | DFn, DFd | Group | | Time | | Interaction | |
|---|---|---|---|---|---|---|---|---|
| | | | F | p | F | p | F | p |
| Figure S2A | Total sleep | 2, 83 | 12.16 | ＜0.0001 | 180.6 | ＜0.0001 | 4.2060 | ＜0.0001 |
| | Sleep episodes | 2, 83 | 10.90 | ＜0.0001 | 15.79 | ＜0.0001 | 2.8380 | ＜0.0001 |
| | P(wake) | 2, 84 | 20.33 | ＜0.0001 | 158.3 | ＜0.0001 | 4.5350 | ＜0.0001 |

Dunnett's multiple comparisons

| | MB299B/dTrpA1 vs MB299B/+ | | | MB299B/dTrpA1 vs dTrpA1/+ | | | Summary | | |
|---|---|---|---|---|---|---|---|---|---|
| | Total sleep | Sleep episodes | P(wake) | Total sleep | Sleep episodes | P(wake) | Total sleep | Sleep episodes | P(wake) |
| ZT0-1 | 0.9271 | 0.8150 | 0.9569 | 0.0025 | 0.0032 | 0.2554 | ns | ns | ns |
| ZT1-2 | 0.0380 | 0.0005 | 0.5383 | 0.3502 | 0.3906 | 0.0436 | ns | ns | ns |
| ZT2-3 | 0.0345 | 0.3404 | 0.0151 | 0.6377 | 0.4221 | 0.1726 | ns | ns | ns |
| ZT3-4 | ＜0.0001 | 0.0091 | 0.0008 | 0.4268 | 0.2148 | 0.5099 | ns | ns | ns |
| ZT4-5 | 0.0026 | 0.1054 | 0.0008 | 0.2269 | 0.4578 | 0.5939 | ns | ns | ns |
| ZT5-6 | ＜0.0001 | 0.8922 | ＜0.0001 | 0.0062 | 0.6089 | 0.0024 | ** | ns | ** |
| ZT6-7 | ＜0.0001 | 0.5051 | ＜0.0001 | 0.0188 | 0.5812 | 0.2621 | * | ns | ns |
| ZT7-8 | 0.6767 | 0.9945 | 0.0488 | 0.0081 | 0.7022 | 0.0039 | ns | ns | * |
| ZT8-9 | 0.0005 | 0.0533 | 0.0046 | 0.1411 | 0.9933 | 0.1829 | ns | ns | ns |
| ZT9-10 | 0.3437 | 0.5029 | 0.0261 | 0.2293 | 0.0146 | 0.1745 | ns | ns | ns |
| ZT10-11 | 0.5884 | 0.3553 | 0.3099 | 0.0125 | 0.1773 | 0.0407 | ns | ns | ns |
| ZT11-12 | 0.0227 | 0.0746 | 0.0095 | 0.1394 | 0.2117 | 0.0179 | ns | ns | ns |
| ZT12-13 | 0.4153 | 0.4524 | 0.0143 | 0.0007 | 0.9532 | 0.0985 | ns | ns | ns |
| ZT13-14 | 0.8556 | 0.8430 | 0.7427 | 0.0148 | 0.8993 | 0.0020 | ns | ns | ns |
| ZT14-15 | 0.5386 | 0.6505 | 0.7164 | 0.0193 | 0.0172 | 0.0207 | ns | ns | ns |
| ZT15-16 | 0.6468 | 0.0463 | 0.7030 | 0.0118 | 0.0470 | 0.0025 | ns | * | ns |
| ZT16-17 | 0.3764 | 0.4060 | 0.1597 | 0.0034 | 0.0234 | 0.0109 | ns | ns | ns |
| ZT17-18 | 0.9319 | 0.6260 | 0.7423 | 0.2286 | 0.9402 | 0.0132 | ns | ns | ns |
| ZT18-19 | 0.8229 | 0.9259 | 0.9782 | 0.5236 | 0.3345 | 0.4937 | ns | ns | ns |
| ZT19-20 | 0.9463 | 0.3941 | 0.9300 | 0.4978 | 0.8826 | 0.6145 | ns | ns | ns |
| ZT20-21 | 0.9414 | 0.5798 | 0.8807 | 0.3993 | 0.6656 | 0.3446 | ns | ns | ns |
| ZT21-22 | 0.3539 | 0.9949 | 0.9476 | 0.7408 | 0.6482 | 0.4334 | ns | ns | ns |
| ZT22-23 | 0.2121 | 0.5568 | 0.2526 | 0.1485 | 0.0365 | 0.5154 | ns | ns | ns |
| ZT23-24 | 0.1292 | 0.0414 | 0.0952 | 0.0139 | ＞0.9999 | 0.0437 | ns | ns | ns |
| ZT24-25 | 0.8655 | 0.7913 | 0.6770 | 0.0139 | 0.0274 | 0.5071 | ns | ns | ns |
| ZT25-26 | 0.9087 | 0.7469 | 0.7692 | 0.0170 | 0.0940 | 0.0012 | ns | ns | ns |
| ZT26-27 | 0.1199 | 0.8552 | 0.0069 | 0.0027 | 0.0055 | 0.1198 | ns | ns | ns |
| ZT27-28 | 0.0468 | 0.1457 | 0.0236 | 0.1713 | 0.0057 | 0.0069 | ns | ns | ns |
| ZT28-29 | 0.0013 | 0.0871 | 0.0115 | 0.9998 | 0.8844 | 0.3795 | ns | ns | ns |
| ZT29-30 | 0.0036 | 0.9752 | 0.0010 | 0.6750 | 0.6980 | 0.9888 | ns | ns | ns |
| ZT30-31 | 0.1281 | 0.8489 | ＜0.0001 | 0.9690 | 0.0034 | 0.9888 | ns | ns | ns |

| | Data | DFn, DFd | Group | | Time | | Interaction | |
|---|---|---|---|---|---|---|---|---|
| | | | F | p | F | p | F | p |
| Figure S2B | Total sleep | 2, 87 | 0.7374 | 0.4813 | 196.3 | ＜0.0001 | 2.6170 | ＜0.0001 |
| | Sleep episodes | 2, 87 | 9.0790 | 0.0003 | 20.05 | ＜0.0001 | 0.8662 | 0.7583 |
| | P(wake) | 2, 87 | 1.4220 | 0.2467 | 181.0 | ＜0.0001 | 2.6300 | ＜0.0001 |

Dunnett's multiple comparisons

| | MB043B/dTrpA1 vs MB043B/+ | | | MB043B/dTrpA1 vs dTrpA1/+ | | | Summary | | |
|---|---|---|---|---|---|---|---|---|---|
| | Total sleep | Sleep episodes | P(wake) | Total sleep | Sleep episodes | P(wake) | Total sleep | Sleep episodes | P(wake) |
| ZT0-1 | 0.0542 | 0.0947 | 0.0895 | 0.0083 | 0.0087 | 0.8686 | ns | ns | ns |
| ZT1-2 | 0.0773 | 0.2427 | 0.5187 | 0.1517 | 0.1117 | 0.1746 | ns | ns | ns |
| ZT2-3 | 0.3812 | 0.7501 | 0.1942 | 0.1885 | 0.3468 | 0.0902 | ns | ns | ns |
| ZT3-4 | 0.8887 | 0.9643 | 0.8913 | 0.5610 | 0.4246 | 0.2496 | ns | ns | ns |
| ZT4-5 | 0.1860 | 0.6133 | 0.3141 | 0.3599 | 0.2616 | 0.9949 | ns | ns | ns |
| ZT5-6 | 0.0257 | 0.3878 | 0.0367 | 0.0074 | 0.3989 | 0.0040 | * | ns | * |
| ZT6-7 | 0.4763 | 0.5832 | 0.3301 | 0.1268 | 0.7228 | 0.4315 | ns | ns | ns |
| ZT7-8 | 0.4915 | 0.9954 | 0.9840 | 0.1722 | 0.8730 | 0.0294 | ns | ns | ns |
| ZT8-9 | 0.9269 | 0.0304 | 0.2414 | 0.2756 | 0.7988 | 0.2011 | ns | ns | ns |
| ZT9-10 | 0.0431 | 0.1218 | 0.1413 | 0.5599 | 0.0637 | 0.3395 | ns | ns | ns |
| ZT10-11 | 0.0397 | 0.2360 | 0.1278 | 0.0063 | 0.0454 | 0.5039 | * | ns | ns |
| ZT11-12 | 0.3379 | 0.7433 | 0.4231 | 0.0322 | 0.2603 | 0.0010 | ns | ns | ns |
| ZT12-13 | 0.0010 | 0.0923 | 0.0232 | 0.0099 | 0.6299 | 0.0261 | ** | ns | * |

| | | | | | | | | | |
|---|---|---|---|---|---|---|---|---|---|
| ZT13-14 | 0.3856 | ＞0.9999 | 0.1911 | 0.5492 | 0.0433 | 0.3573 | ns | ns | ns |
| ZT14-15 | 0.8210 | 0.9296 | 0.9701 | 0.9751 | 0.6404 | 0.7050 | ns | ns | ns |
| ZT15-16 | 0.2657 | 0.4844 | 0.8947 | 0.3767 | 0.6163 | 0.4584 | ns | ns | ns |
| ZT16-17 | 0.0167 | 0.7103 | 0.1198 | 0.0211 | 0.5542 | 0.2998 | * | ns | ns |
| ZT17-18 | 0.3970 | 0.7614 | 0.1917 | 0.5029 | 0.3777 | 0.2287 | ns | ns | ns |
| ZT18-19 | 0.4264 | 0.4994 | 0.2790 | 0.8455 | 0.1745 | 0.9340 | ns | ns | ns |
| ZT19-20 | 0.8153 | 0.8355 | 0.9636 | 0.7050 | 0.0930 | 0.9907 | ns | ns | ns |
| ZT20-21 | 0.4155 | 0.5319 | 0.8338 | 0.1225 | 0.6983 | 0.2679 | ns | ns | ns |
| ZT21-22 | 0.7446 | 0.9676 | 0.9880 | 0.6336 | 0.9920 | 0.3069 | ns | ns | ns |
| ZT22-23 | 0.6136 | 0.4624 | 0.9295 | 0.1035 | 0.9415 | 0.2829 | ns | ns | ns |
| ZT23-24 | 0.3903 | 0.2161 | 0.5840 | 0.3318 | 0.5054 | 0.1491 | ns | ns | ns |
| ZT24-25 | 0.3200 | 0.8207 | 0.0750 | 0.2397 | 0.6446 | 0.7277 | ns | ns | ns |
| ZT25-26 | 0.1311 | 0.9551 | 0.3670 | 0.1416 | 0.8683 | 0.0840 | ns | ns | ns |
| ZT26-27 | 0.3947 | 0.9924 | 0.1422 | 0.0485 | 0.1254 | 0.1170 | ns | ns | ns |
| ZT27-28 | 0.7008 | ＞0.9999 | 0.4673 | 0.2948 | 0.0098 | 0.0488 | ns | ns | ns |
| ZT28-29 | 0.1018 | 0.3762 | 0.1138 | 0.9999 | 0.1909 | 0.7851 | ns | ns | ns |
| ZT29-30 | 0.1215 | 0.0680 | 0.0808 | 0.5694 | 0.1893 | 0.9435 | ns | ns | ns |
| ZT30-31 | 0.0503 | 0.1580 | 0.0209 | 0.7486 | 0.2126 | 0.8009 | ns | ns | ns |

| | | | Group | | Time | | Interaction | |
|---|---|---|---|---|---|---|---|---|
| | Data | DFn, DFd | F | p | F | p | F | p |
| Figure S2C | Total sleep | 2, 98 | 1.0650 | 0.3487 | 222.1 | ＜0.0001 | 3.7180 | ＜0.0001 |
| | Sleep episodes | 2, 98 | 2.5120 | 0.0863 | 32.28 | ＜0.0001 | 1.9580 | ＜0.0001 |
| | P(wake) | 2, 98 | 0.8850 | 0.4160 | 181.4 | ＜0.0001 | 5.3520 | ＜0.0001 |

| | | | Group | | Time | | Interaction | |
|---|---|---|---|---|---|---|---|---|
| | Data | DFn, DFd | F | p | F | p | F | p |
| Figure S2D | Total sleep | 2, 101 | 6.0000 | 0.0034 | 167.4 | ＜0.0001 | 5.2050 | ＜0.0001 |
| | Sleep episodes | 2, 101 | 13.20 | ＜0.0001 | 17.65 | ＜0.0001 | 2.5730 | ＜0.0001 |
| | P(wake) | 2, 104 | 2.194 | 0.1166 | 135.3 | ＜0.0001 | 7.3347 | ＜0.0001 |

Dunnett's multiple comparisons

| | MB043B/$shi^{ts1}$ vs MB043B/+ | | | MB043B/$shi^{ts1}$ vs $shi^{ts1}$/+ | | | Summary | | |
|---|---|---|---|---|---|---|---|---|---|
| | Total sleep | Sleep episodes | P(wake) | Total sleep | Sleep episodes | P(wake) | Total sleep | Sleep episodes | P(wake) |
| ZT0-1 | 0.1800 | 0.4486 | 0.0048 | 0.0103 | 0.0085 | 0.1223 | ns | ns | ns |
| ZT1-2 | 0.9957 | 0.1574 | 0.5541 | 0.1346 | 0.4120 | 0.0026 | ns | ns | ns |
| ZT2-3 | 0.8998 | 0.0948 | 0.2824 | 0.6483 | 0.5005 | 0.8577 | ns | ns | ns |
| ZT3-4 | 0.9987 | 0.0428 | 0.9950 | 0.0070 | 0.0336 | 0.2015 | ns | * | ns |
| ZT4-5 | 0.1434 | 0.0262 | 0.5763 | <0.0001 | 0.8067 | 0.0006 | ns | ns | ns |
| ZT5-6 | 0.3662 | 0.9111 | 0.0273 | <0.0001 | 0.0406 | <0.0001 | ns | ns | * |
| ZT6-7 | 0.0409 | 0.5251 | 0.7056 | <0.0001 | 0.9842 | <0.0001 | * | ns | ns |
| ZT7-8 | 0.0334 | 0.0232 | 0.0003 | 0.8532 | 0.3051 | <0.0001 | ns | ns | *** |
| ZT8-9 | 0.0390 | 0.0486 | 0.0054 | 0.9213 | 0.9037 | 0.3300 | ns | ns | ns |
| ZT9-10 | 0.6141 | 0.7464 | 0.8216 | 0.1060 | 0.8155 | 0.6512 | ns | ns | ns |
| ZT10-11 | 0.8941 | 0.9994 | 0.5373 | 0.6583 | 0.0818 | 0.0220 | ns | ns | ns |
| ZT11-12 | 0.2796 | 0.9921 | 0.7696 | 0.0028 | 0.0009 | 0.0023 | ns | ns | ns |
| ZT12-13 | 0.7142 | 0.8039 | 0.6344 | 0.9829 | 0.7743 | 0.2227 | ns | ns | ns |
| ZT13-14 | 0.1356 | 0.2098 | 0.5681 | 0.0129 | 0.0274 | 0.2156 | ns | ns | ns |
| ZT14-15 | 0.0098 | 0.2677 | 0.0161 | 0.0082 | 0.0048 | 0.0168 | ** | ns | * |
| ZT15-16 | 0.1530 | 0.0158 | 0.0192 | 0.0623 | 0.0155 | 0.0329 | ns | * | * |
| ZT16-17 | 0.0273 | 0.0110 | 0.0684 | 0.0072 | 0.0029 | 0.0475 | * | * | ns |
| ZT17-18 | 0.0531 | 0.1402 | 0.1046 | 0.0115 | 0.0153 | 0.0087 | ns | ns | ns |
| ZT18-19 | 0.0609 | 0.0210 | 0.1614 | 0.0004 | 0.0002 | 0.0038 | ns | * | ns |
| ZT19-20 | 0.5793 | 0.0212 | 0.3301 | 0.0489 | 0.0513 | 0.0029 | ns | ns | ns |
| ZT20-21 | 0.7019 | 0.2464 | 0.9120 | 0.0485 | 0.0173 | 0.3255 | ns | ns | ns |
| ZT21-22 | 0.9867 | 0.5995 | 0.7299 | 0.0279 | 0.0600 | 0.0086 | ns | ns | ns |
| ZT22-23 | 0.9994 | 0.8742 | 0.8509 | 0.6676 | 0.1044 | 0.3311 | ns | ns | ns |
| ZT23-24 | 0.9922 | 0.1545 | 0.4956 | 0.7800 | 0.2270 | 0.9768 | ns | ns | ns |
| ZT24-25 | 0.0029 | 0.1390 | 0.5547 | 0.0019 | 0.0072 | 0.9952 | ** | ns | ns |
| ZT25-26 | 0.0096 | 0.8586 | 0.0225 | 0.0065 | 0.0013 | <0.0001 | ** | ns | * |
| ZT26-27 | 0.0539 | 0.3434 | 0.2088 | 0.5070 | 0.4646 | 0.0339 | ns | ns | ns |
| ZT27-28 | 0.1591 | 0.3726 | 0.2383 | 0.1718 | 0.2547 | 0.9173 | ns | ns | ns |
| ZT28-29 | 0.7048 | 0.2134 | 0.1502 | 0.0149 | 0.8512 | 0.1055 | ns | ns | ns |
| ZT29-30 | 0.8512 | 0.1013 | 0.9681 | 0.0451 | 0.0135 | 0.0077 | ns | ns | ns |
| ZT30-31 | 0.6151 | 0.6842 | 0.9183 | 0.0398 | 0.2023 | 0.0615 | ns | ns | ns |
